# Rule-based meta-analysis reveals the major role of PB2 in influencing influenza A virus virulence in mice

**DOI:** 10.1101/556647

**Authors:** Fransiskus Xaverius Ivan, Chee Keong Kwoh

## Abstract

**Background:** Influenza A virus (IAV) poses threats to human health and life. Many individual studies have been carried out in mice to uncover the viral factors responsible for the virulence of IAV infections. Virus adaptation through serial lung-to-lung passaging and reverse genetic engineering and mutagenesis approaches have been widely used in the studies. Nonetheless, a single study may not provide enough confident about virulence factors, hence combining several studies for a meta-analysis is desired to provide better views.

**Methods:** Virulence information of IAV infections and the corresponding virus and mouse strains were documented from literature. Using the mouse lethal dose 50, time series of weight loss or percentage of survival, the virulence of the infections was classified as avirulent or virulent for two-class problems, and as low, intermediate or high for three-class problems. On the other hand, protein sequences were decoded from the corresponding IAV genomes or reconstructed manually from other proteins according to mutations mentioned in the related literature. IAV virulence models were then learned from various datasets containing IAV proteins whose amino acids at their aligned position and the corresponding two-class or three-class virulence labels. Three proven rule-based learning approaches, i.e., OneR, JRip and PART, and additionally random forest were used for modelling, and top protein sites and synergy between protein sites were identified from the models.

**Results:** More than 500 records of IAV infections in mice whose viral proteins could be retrieved were documented. The BALB/C and C57BL/6 mouse strains and the H1N1, H3N2 and H5N1 viruses dominated the infection records. PART models learned from full or subsets of datasets achieved the best performance, with moderate averaged model accuracies ranged from 65.0% to 84.4% and from 54.0% to 66.6% for two-class and three-class datasets that utilized all records of aligned IAV proteins, respectively. Their averaged accuracies were comparable or even better than the averaged accuracies of random forest models and should be preferred based on the Occam’s razor principle. Interestingly, models based on a dataset that included all IAV strains achieved a better averaged accuracy when host information was taken into account. For model interpretation, we observed that although many sites in HA were highly correlated with virulence, PART models based on sites in PB2 could compete against and were often better than PART models based on sites in HA. Moreover, PART had a high preference to include sites in PB2 when models were learned from datasets containing concatenated alignments of all IAV proteins. Several sites with a known contribution to virulence were found as the top protein sites, and site pairs that may synergistically influence virulence were also uncovered.

**Conclusion:** Modelling the virulence of IAV infections is a challenging problem. Rule-based models generated using only viral proteins are useful for its advantage in interpretation, but only achieve moderate performance. Development of more advanced machine learning approaches that learn models from features extracted from both viral and host proteins must be considered for future works.

## Introduction

Influenza A virus (IAV) is a member of the family *Orthomyxoviridae* that circulates in humans, mammals and birds. The genome of the virus consists of 8 single-stranded, negative-sense viral RNA segments encoding at least 12 proteins that make up its proteome (**Table 1**). The surface glycoproteins HA and NA proteins play a role in the entry into a host cell and exit from the host cell, respectively. Each viral RNA is packaged with multiple copies of NP protein and an RNA polymerase complex that comprises PA, PB1 and PB2 proteins, to form a rod-like ribonucleoprotein complex [1]. The RNA polymerase complex plays a role in both transcription and replication of the viral genomes. The M1 protein mediates virion assembly, while the M2 protein forms a proton channel that is required for viral entry. The NS1 and NS2 proteins are multifunctional. For examples, NS1 is well known to inhibit interferon related activities (reviewed in [2]), while NS2 has been implicated in mediating the nuclear export of RNP complexes and the recruiting ATPase for efficient viral exit (reviewed in [3]). PB1-F2 and PA-X proteins are non-essential and encoded by a +1 alternate open reading frame in the PB1 and PA, respectively. PB1-F2 and PA-X play a role in IAV pathogenesis [4, 5].

**Table 1.** IAV segments and their encoded proteins

| Segment | Protein 1 (p1) | Protein 2 (p2) |
| --- | --- | --- |
| 1 – PB2 | RNA polymerase B2 (PB2) |  |
| 2 – PB1 | RNA polymerase B1 (PB1) | Non-structural protein PB1-F2 |
| 3 – PA | RNA polymerase A (PA) | Non-structural protein PA-X |
| 4 – HA | Hemagglutinin (HA) |  |
| 5 – NP | Nucleoprotein (NP) |  |
| 6 – NA | Neuraminidase (NA) |  |
| 7 – M | Matrix protein 1 (M1) | Matrix protein 2 (M2; also known as ion channel protein) |
| 8 – NS | Non-structural protein 1 (NS1) | Non-structural protein 2 (NS2; also known as nuclear export protein (NEP)) |

The HA and NA determine the subtype of IAV. To date, 18 HA (H1-H18) and 11 NA (N1-N11) have been identified. IAV of H1N1, H2N2, and H3N2 subtypes have been responsible for five pandemics of severe human respiratory diseases in the last 100 years, i.e., the 1918 Spanish Influenza (H1N1), 1957 Asian Influenza (H2N2), 1968 Hong Kong (H3N2), 1977 Russian Influenza (H1N1), and 2009 Swine-Origin Influenza (H1N1) pandemics. The pandemic strains continuously spread among humans and cause recurrent, seasonal epidemics. In the last few years, the seasonal human IAVs were mainly dominated by the 1968’s H3N2 and 2009’s H1N1 strains. In addition to epidemic and pandemic strains, several IAV subtypes have also caused human infections, including the H5N1, H5N6, H6N1, H7N2, H7N3, H7N7, H7N9, H9N2, and H10N8 avian influenza viruses [6, 7]. Among them, the H5N1 and H7N9 subtypes have raised a major public health concern due to their ability to cause human outbreaks with high fatality rate (about 60% (www.who.int) and 39% [8], respectively). Overall, IAV poses a threat to human health and life, and therefore further understanding about the virus is needed for a better surveillance and counteractive measures against it.

Many aspects of IAV and the disease it causes have been investigated in mice since the animals are not only cost-effective and easy to handle, but also available in various inbred, transgenic, and knockout strains. Moreover, the genomes of various inbred mice have been recently available. Mice have also allowed us to uncover host and viral molecular determinants of IAV virulence. Early outcome of IAV study in mice was the revelation of the protective role of interferon-induced gene Mx1 against the virus [9]. Recently, the gene has been shown to inhibit the assembly of functional viral ribonucleoprotein complex of IAV [10]. In the last 50 years, the importance of many more host genes in influenza pathogenesis has been discovered through experiments in mice, including RIG-I, IFITM3, TNF and IL-1R genes (reviewed in [11, 12]). Nonetheless, one limitation of the existing approaches in investigating host molecular determinants involved in IAV virulence is that it has not yet taken into account the contribution of allelic variation to differential host responses.

In contrast, the influence of variations in viral genes to IAV virulence have been investigated in a number of ways. These included the generation of mouse-adapted IAVs through serial lung-to-lung passaging and recombinant IAVs harboring specific mutations using plasmid-based reverse genetic techniques combined with mutagenesis approaches. The application of these techniques has provided various insights about viral mutations involved in IAV virulence. For example, the increased virulence of IAV during its adaptation in mice has been associated with mutations in the region 190-helix, 220-loop and 130-loop, which surround the receptor-binding site in the HA protein (reviewed in [13]). Mutations in PB2 have also been considered to play a significant role in the increased IAV virulence in mice, which include mutations E627K and D701N that are considered as general markers for IAV virulence in mice [11]. Interestingly, a single mutation N66S in the accessory protein PB1-F2 could also contribute to increased virulence [14]. Mutations in multiple sites of a specific viral protein and mutations in multiple genes have also been shown to have a synergistic effect on IAV virulence in mice. For example, synergistic effect of dual mutations S224P and N383D in PA led to increased polymerase activity and has been considered as a hallmark for natural adaptation of H1N1 and H5N1 viruses to mammals [15]. Another example is the synergistic action of two mutations D222G and K163E in HA and one mutation F35L in PA of pandemic 2009 influenza A/H1N1 virus that causes lethality in the infected mice [16]. Furthermore, virulence may not only be encoded at protein level, but also at nucleotide level. In a very recent study, synonymous codons were interestingly able to give rise different virulence levels [17].

The confidence of contribution of viral protein sites to the virulence of influenza infections could be better investigated through a meta-analysis approach, which is a systematic amalgamation of results from individual studies. Such approach, to our knowledge, has only been carried out using a Bayesian graphical model to investigate the viral protein sites important for virulence of influenza A/H5N1 in mammals [18]. Nevertheless, a meta-analysis approach using Naive Bayes approach at viral nucleotide level has recently been carried out to demonstrate the contribution of synonymous nucleotide mutations to IAV virulence [17]. In this paper we present a meta-analysis of viral protein sites that determine the virulence of infections with any subtype of IAV; however, instead of any mammal, we focus on the infections in mice. Our meta-analysis approach utilized rule-based machine learnings and random forest to predict IAV virulence from datasets we created. The creation of the datasets involved: (*i*) documentation of the virulence of infections involving particular IAV and mouse strains, (*ii*) classification of virulence levels, and (*iii*) collection of the corresponding IAV proteins. For learning IAV virulence models, each column of the alignments was considered as a feature vector and the virulence levels as a target vector. When host information was considered, the amino acids in the columns were tagged with a symbol representing the corresponding mouse strain. The models were developed using either all records in the datasets or records for a specific mouse strain or influenza subtype, and using concatenated alignments of all IAV proteins or an individual alignment of particular IAV proteins. Top protein sites and synergy between protein sites were then examined for some biological interpretations.

## Methods

### Collection of IAV infections in mice with virulence information

We collected journal publications containing virulence information of IAV infection in non-transgenic and non-knock-out inbred mice. Each unique infection involving specific IAV strain and specific mouse strain and with known value of MLD50 was recorded. Infections without MLD50 values but whose time series of weight loss or percentage of survival of infected mice per infection dose could be estimated from the relevant figures, were also recorded and used to estimate the lower or upper bound of MLD50; few of them were used to estimate the exact MLD50 using the Reed and Muench method [19]. Various units for MLD50, which include the plaque forming unit (PFU), focus forming unit (FFU), egg infectious dose (EID50), tissue culture infectious dose (TCID50), and cell culture infectious dose (CCID50), were assumed to measure the same quantity.

Next, the levels of virulence were categorized into two classes, i.e., avirulent and virulent. If the MLD50 of an infection is >10E6.0 (regardless of its unit), then the infection is considered avirulent; otherwise, virulent. When the class of an infection cannot be determined from the lower or upper bound of MLD50, then the following rules were used:

### RULE 1

An infection is avirulent if:

i. the infection dose between 10E4.0 and 10E6.0 leads to <15% average weight loss;
ii. the infection dose ≥10E5.0 does not kill any mouse; or
iii. the infection dose between 10E3.0 and 10E4.0 leads to ≤10% average weight loss.

### RULE 2

An infection is virulent if:

i. the infection dose ≤10E5.0 leads to ≥15% average weight loss;
ii. the infection dose ≤10E3.0 leads to ≥10% average weight loss; or
iii. the infection dose ≤10E3.5 kills ≥10% mice.

The levels of virulence were also categorized into three classes: low, intermediate and high virulence. If the MLD50 >10E6.0, then the infection is considered low virulence; if the MLD50 ≤10E3.0, then the infection is considered high virulence; otherwise, intermediate virulence. When the class of an infection cannot be determined from the lower or upper bound of MLD50, then the following rules were used:

### RULE 3

An infection is low virulence if it is considered avirulent (as given in the two class labelling).

### RULE 4

An infection is intermediate virulence if:

i. the infection dose <10E4.0 leads to ≥10% average weight loss;
ii. the infection dose between 10E4.0 and 10E5.0 leads to ≥15% average weight loss; or
iii. the infection dose between 10E5.0 and 10E6.0 leads to ≥20% average weight loss.

### RULE 5

An infection is high virulence if:

i. the infection dose ≤10E6.0 kills ≥80% mice or leads to ≥25% average weight loss; or
ii. the infection dose ≤10E1.0 kills ≥20% mice.

Following this, multiple records for infection involving specific IAV and mouse strains were reduced into a single record (**Table S2**) by the following procedure (termed as **RULE 6**):

i. Specify the majority class of the three-class virulence assignment for those records; when no majority, consider the class that is more or the most virulent.
ii. Select the record with:

- the highest lower bound of MLD50 value when only lower bound of MLD50 values presented;
- the lowest exact or upper bound of MLD50 value when they are available; but when the highest lower bound of MLD50 value is lower than this value, then calculate the average of those two values and assign the virulence class as described previously.

This procedure selects a record that has the more or most virulent information among the records (with the majority class if it can be determined), except when only lower bound of MLD50 values are available. Note that when applying this procedure, the recombinants of naturally occurring or wild-type IAV strains were considered representing the wild-type version. In a similar fashion, we applied this procedure to reduce multiple records for infection of a specific IAV strain in different mouse strains into a single record (**Table S3**).

### Collection of related genomes and main proteins

IAV strains found in the literature were searched online by their name, and their nucleotide sequences were collected from GenBank or GISAID EpiFlu databases. A number of sequences were obtained from the authors directly. When the genomic segments of a particular virus were incomplete, the HA and/or NA of the virus were BLASTed against GenBank database and the top virus hit whose complete genomes were available was used to extrapolate the incomplete genome (**Table S4**). Considering the closeness between their names, the genome of influenza A/Turkey/15/2006(H5N1) was used to represent the genome of influenza A/Turkey/13/2006(H5N1) that was not available. Furthermore, we extrapolated partial IAV sequences by using the closest complete IAV sequence identified by BLAST (**Table S5**). Then, the reassortant viruses reported in the literature were reconstructed using relevant genomic segments. Following the collection of IAV genomes, the 12 IAV proteins were obtained by identifying their coding sequence regions using Influenza Virus Sequence Annotation Tool available at the NCBI Influenza Virus Resource and then translating them into proteins according to standard genetic code. Some proteins, mainly for mutant viruses, were generated from existing proteins according to the list of amino acid differences at various sites reported in the literature. Note that some IAVs were represented by different versions of genomes or sets of proteins, but the reassortant or mutant viruses were mainly reconstructed from one of the versions.

### Machine learning approaches for IAV virulence prediction

Three rule-based machine learning approaches, i.e., OneR, JRip and PART that are available in RWeka [20], and random forest (RF) available in randomForest package for R [21] were explored to develop predictive models for IAV virulence. Various input datasets were considered (see the first section of results), but in general, the input datasets consisted of IAV proteins that have been aligned with muscle package [22] and their target virulence class. The datasets included either the alignments of all IAV proteins or an individual alignment of particular IAV proteins. Each column in the alignment that contained more than one symbol was considered as a single feature vector – H3 and N2 numberings were used to label the position in the alignments of HA and NA, respectively. Input datasets that incorporated the host strain information, where each amino acid in the alignments was tagged with a symbol indicating associated host strain, were also considered. For each input dataset, each learning algorithm and each of two-class and three-class virulence groupings, rule-based and RF models were learned independently 100 times. In each iteration, the dataset was balanced by reducing the size of the bigger (biggest) class to the size of the smaller (smallest) class through sampling without replacement. To develop a learning model, 60% of the records (rows of the alignment) from each virulent class were used as training data, while the rest were used as test data. Performance metrics that included accuracy, macro-averaged precision and macro-averaged recall were calculated to evaluate the models.

### Visualization, statistical analyses and site rankings

The concatenated alignments of IAV proteins were visualized in 3D Cartesian coordinates. For this, a matrix of pairwise distances from concatenated protein alignments was computed using Fitch similarity matrix and then the Kruskal’s non-metric multidimensional scaling available in R’s MASS package [23] was applied to place each record of concatenated protein sequences in a 3D space.

The correlations between sites in the alignment and the target virulence class were measured using the Benjamini-Hochberg adjusted p-values of the chi-square test of independence. The –log(adjusted p-value) of the test over the sites of each IAV protein was visualized with a line plot.

Wilcoxon signed-rank sum test was used to test the null hypothesis that the median of the accuracy of 100 models learned independently is equal to the accuracy of zero rule learner (which assigns predicted class to the majority class in the training set) and to test the null hypothesis that the median of the accuracy of one learner is greater than that of another learner. The p-values of the tests were adjusted using the Bonferroni method.

Following 100 independent learnings from two-class and three-class IV datasets, the protein sites from models learned using each algorithm were ranked. For OneR, the sites were ranked according to their frequency of being selected for the models; for JRip and PART, the sites were ranked according to their averaged contribution to the accuracy of learned models; and for RF, the sites were ranked according to their contribution to the averaged mean decrease in accuracy. For PART models, we also ranked the site pairs according to their averaged contribution to the accuracy of learned models and visualized the synergistic graph arises from the top 50 site pairs using igraph package for R software [24].

## Results

### Datasets for modelling IAV virulence

The steps in creating benchmark datasets for modeling IAV virulence is summarized in Fig. 1. Initially, a dataset containing 637 records of IAV infections in mice, where the full or incomplete genome of the IAVs could be retrieved from public sequence databases and the virulence class of the infection could be identified, was created according to information available in 84 journal publications (**Table S1**). Of those records, 502 records have their MLD50 provided in the literature. Following **RULE 6** (see Methods), multiple records involving specific IAV and mouse strains were reduced into a single record (**Table S2**). This produced a new dataset containing 555 records and named as Mouse-IAV Virulence (MIVir) dataset. Using the same rule, the MIVir dataset was further reduced to a dataset containing 489 records of IAV virulence across different mouse strains and named as IAV Virulence (IVir) dataset.

**Fig. 1.**
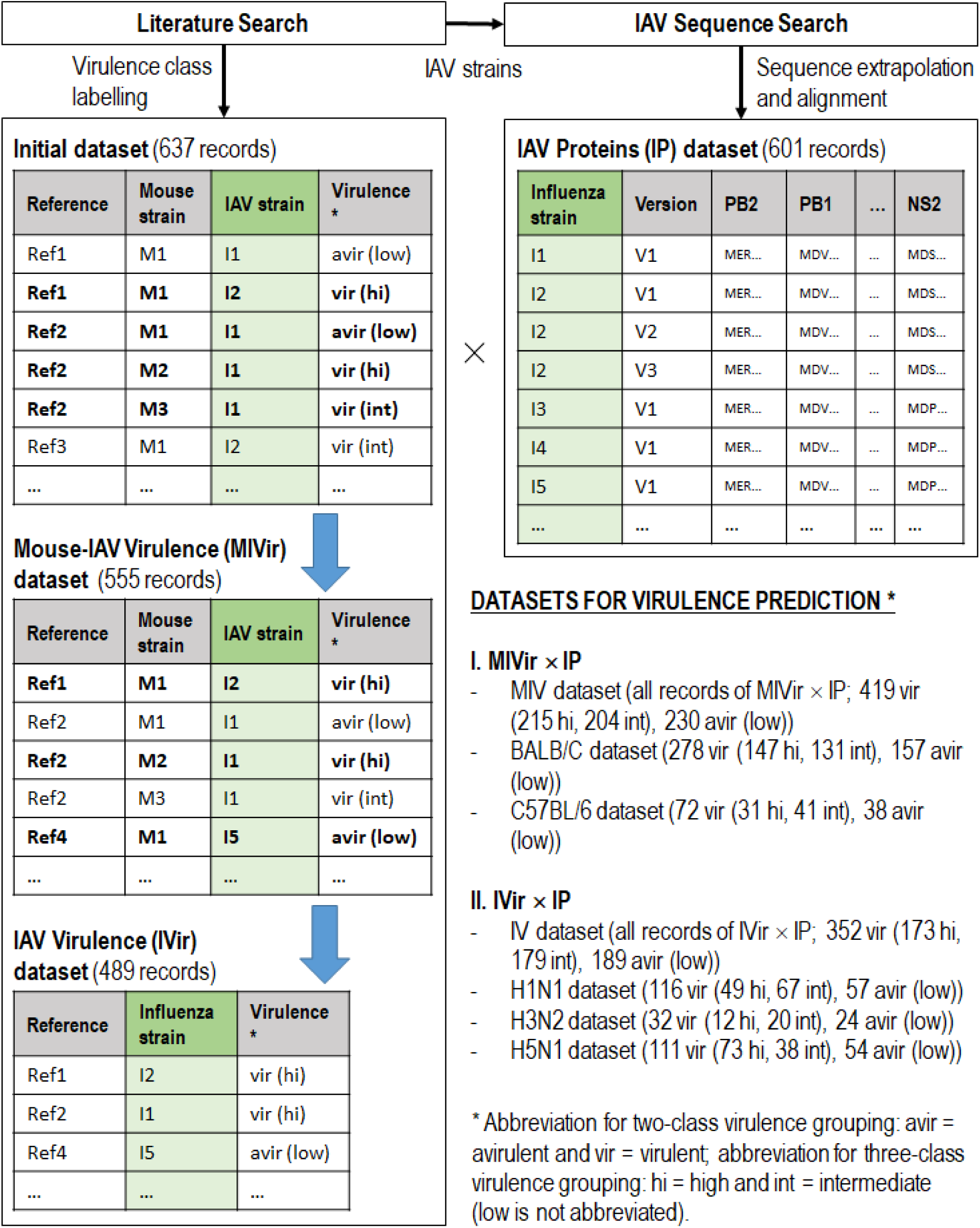
Creation of benchmark datasets for IAV virulence prediction. Note that for simplicity, only the two-class and three-class virulence labels are illustrated in the table, while original or estimate of LD50 is not shown.

The MIVir and IVir datasets were then joined with another dataset containing the 12 IAV proteins whose amino acids in their aligned position (IAV Proteins (IP) dataset), producing MIVir × IP and IVir × IP datasets, respectively. The keys for joining the dataset were the IAV strains listed in MIVir or IVir dataset. Once again, note that some virus strains were represented by multiple records in IP dataset and some proteins were generated from extrapolated genomes.

The breakdowns of the joined datasets are shown in Fig. 1, and more detailed breakdowns of MIVir × IP are shown in Table 2. As shown in the figure and table, the final datasets were mainly dominated by experiments involving BALB/C and C57BL/6 mice and IAV subtypes H1N1, H3N2 and H5N1. Much lesser mouse strains in the records included the 129S1/SvImJ, 129S1/SvPasCrlVr, A/J, C3H, CAST/EiJ, CBA/J, CD-1, DBA/2, FVB/NJ, ICR, NOD/ShiLtJ, NZO/HILtJ, PWK/PhJ, SJL/JOrlCrl, and WSB/EiJ mice, while much lesser IAV subtypes included the H1N2, H3N8, H5N2, H5N5, H5N6, H5N8, H6N1, H7N1, H7N2, H7N3, H7N7, H7N9 and H9N2. Subsets of MIVir × IP dataset used for virulence prediction included dataset containing all records (named as MIV dataset) and datasets containing records of infections in BALB/C and C57BL/6 mice (BALB/C and C57BL/6 datasets, respectively); while subsets of IVir × IP dataset used for virulence prediction included dataset containing all records (IV dataset) and datasets containing infections with H1N1, H3N2 and H5N1 viruses (H1N1, H3N2 and H5N1 datasets, respectively).

**Table 2.** Cross-tabulation between mouse strains and IAV subtypes in MIV dataset. The number at the top in each cell corresponds to the number of records of relevant infections, and the number of cases for each of three-virulent class, i.e., high, intermediate and low virulence, are shown in order in the bracket. The number of virulent cases for two-class virulence grouping is the sum of the number of high and intermediate virulence cases, while the number of avirulent cases equals to the number of low virulence cases.

| Mouse strain | IAV subtype |  |  |  | Total |
| --- | --- | --- | --- | --- | --- |
|  | H1N1 | H3N2 | H5N1 | Others |  |
| BALB/C | 123<br>(35/40/48) | 14<br>(4/2/8) | 162<br>(69/40/53) | 136<br>(39/49/48) | 435<br>(147/131/157) |
| C57BL/6 | 61<br>(14/34/13) | 17<br>(1/2/14) | 6<br>(6/0/0) | 26<br>(10/5/11) | 110<br>(31/41/38) |
| CD-1 | 0<br>(0/0/0) | 34<br>(5/16/13) | 0<br>(0/0/0) | 0<br>(0/0/0) | 34<br>(5/16/13) |
| DBA/2 | 21<br>(14/5/2) | 15<br>(2/5/8) | 0<br>(0/0/0) | 6<br>(2/2/2) | 42<br>(18/12/12) |
| Others | 19<br>(9/3/7) | 7<br>(5/0/2) | 1<br>(0/0/1) | 1<br>(0/1/0) | 28<br>(14/4/10) |
| Total | 224<br>(72/82/70) | 87<br>(17/25/45) | 169<br>(75/40/54) | 169<br>(51/57/61) | 649<br>(215/204/230) |

### Visualization of IV dataset

For an initial view of the IAV sequences being used for virulence prediction, the 3D MDS plot that visualizes the level of similarity between concatenated alignments of IAV proteins in the IV dataset is presented in Fig. 2. While the clusters of dominant IAV subtypes can be easily observed in the plot, separation between virulence classes is lack and this illustrates the challenge in the prediction.

**Fig. 2.**
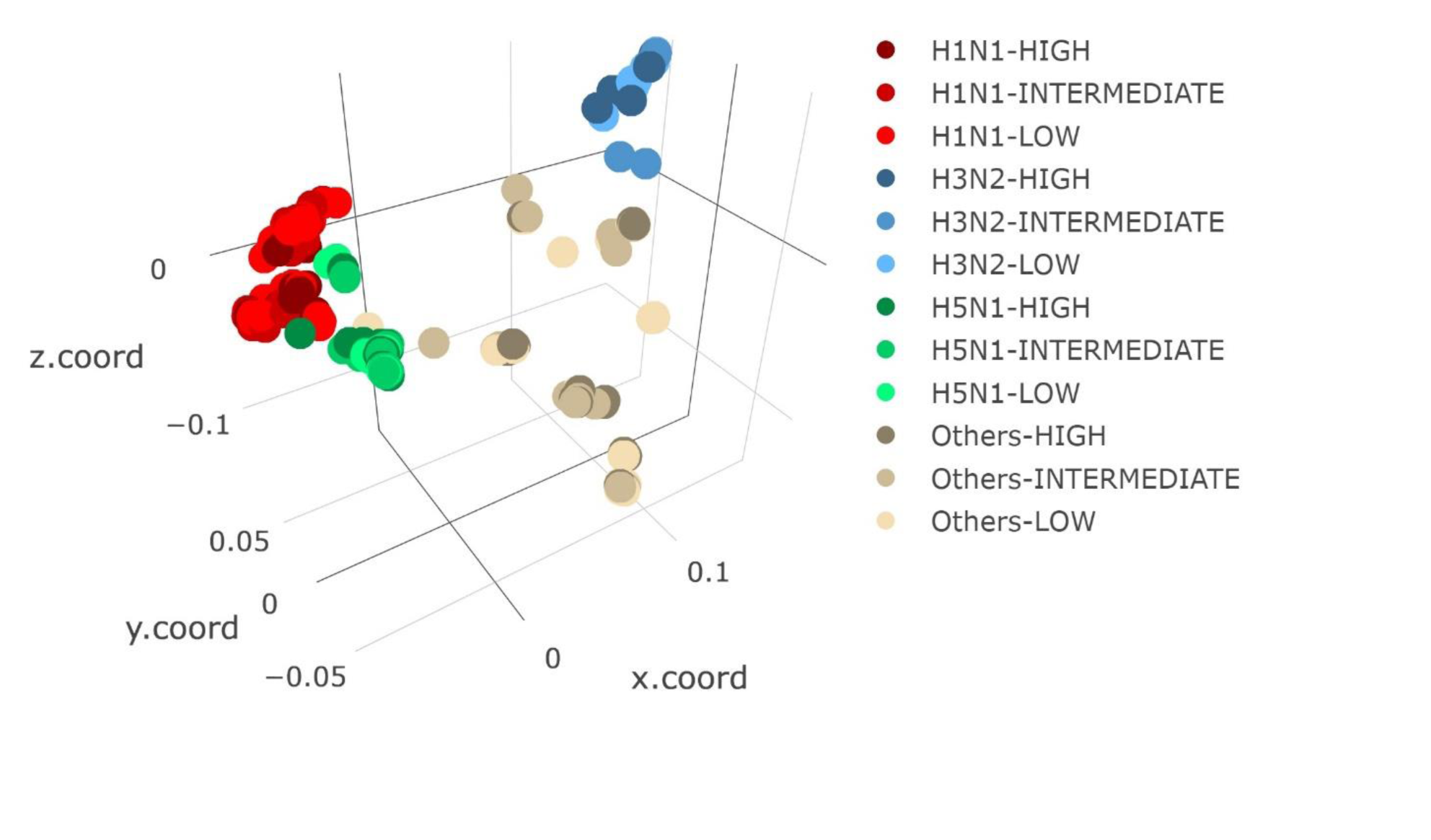
Three-dimensional MDS plot of concatenated alignments of IAV proteins. Each data point representing a record of concatenated IAV proteins is colored based on the subtype and three-class virulence label of associated virus in three-class IV dataset.

In addition, the correlation between each site and the target virulence in the dataset was also measured using the adjusted p-value of the chi-square test of independence. The line plots showing the –log(adjusted p-value) over the alignment sites of each IAV protein and each of two-class and three-class virulence groupings are given in Fig. 3. Overall, HA has many more sites that have a significant correlation with the target virulence (adjusted p-value <0.05), i.e., 72 and 283 sites for two-class and three-class virulence grouping, respectively. On the other hand, M2 has the least numbers of significant sites, i.e., 1 and 4 for two-class and three-class virulence, respectively. The numbers of significant sites for other proteins and for two-class and three-class virulence grouping, respectively, are as follows: 26 and 44 for PB2, 6 and 30 for PB1, 14 and 33 for PA, 19 and 40 for NP, 19 and 167 for NA, 4 and 10 for M1, 18 and 32 for NS1, 3 and 30 for PB1-F2, 6 and 26 for PA-X, and 3 and 5 for NS2. Interestingly, while PB2, PA, NP, M1, NS1 and NS2 have their number of significant sites for three-class virulence about twice the number of significant sites for two-class virulence, the PB1, HA, NA, PB1-F2 and PA-X have a much higher fold increase in the number of significant sites.

**Fig. 3.**
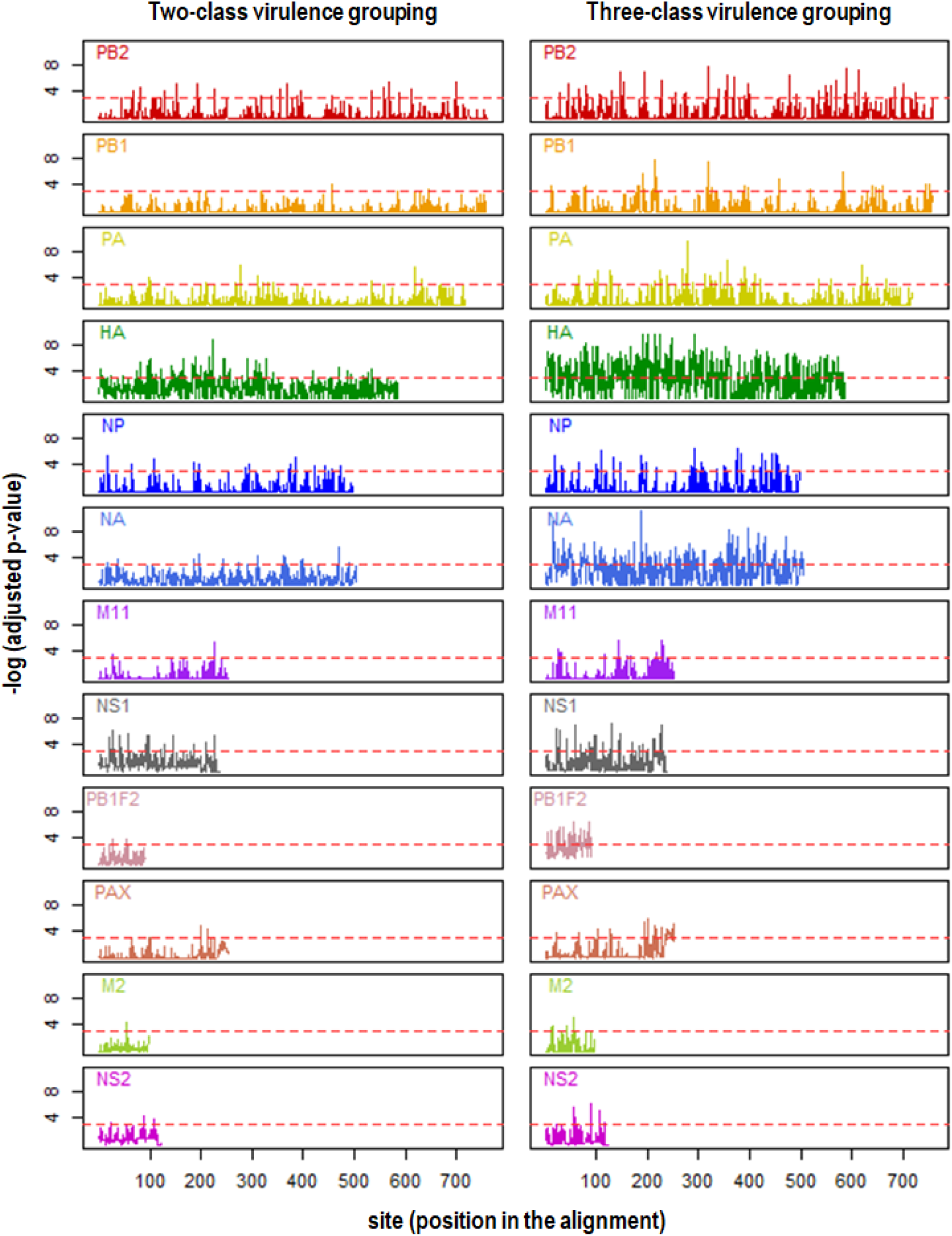
Correlations between sites in the protein alignment and their target virulence in the (A) two-class and (B) three-class IV datasets. The Benjamini-Hochberg adjusted p-values of the chi-square test for independence between sites and their target virulence are used as a measure of the correlation. The red dashed horizontal line in each plot refers to the threshold for the significance of the tests (adjusted p-value <0.05).

### Performance of rule-based models for IAV virulence

Here we focus on the application of OneR, JRip and PART algorithms on MIV, BALB/C, C57BL/6, IV, H1N1 and H3N2 datasets in developing rule-based models for IAV virulence. Table 3 highlights the performance of OneR, JRip and PART on various two-class and three-class datasets with concatenated protein alignments, while examples of the output models and their summary (for H1N1) are presented in **Table S6**. Overall, in terms of their accuracy, precision and recall (but we mainly focus on the accuracy in the rest of the paper), PART models always outperformed OneR and JRip, while JRip were almost always better than OneR (the only case where OneR outperformed JRip was on the three-class classification problem for H3N2). Nonetheless, PART had many more rules compared to JRip and OneR. For example, on IV dataset, PART had on average 10.67 and 46.97 rules per model for two-class and three-class virulence grouping, respectively; while JRip had on average 3.89 and 4.55, respectively, and OneR always had 1 rule.

**Table 3.** Accuracy, macro-averaged precision and macro-averaged recall of models generated by OneR, JRip and PART from various input datasets containing concatenated alignments of IAV proteins. For each cell, the number at the top is the mean of performance values calculated from 100 models learned independently; while the number in the bracket is related standard deviation.

|  | Accuracy (%) |  |  | Macro-averaged Precision (%) |  |  | Macro-averaged Recall (%) |  |  |
| --- | --- | --- | --- | --- | --- | --- | --- | --- | --- |
|  | OneR | JRip | PART | OneR | JRip | PART | OneR | JRip | PART |
| **Two-class virulence grouping** |  |  |  |  |  |  |  |  |  |
| MIV | 58.6<br>(3.6) | 58.8<br>(5.9) | **71.8**<br>(3.8) | 59.1<br>(3.8) | 59.9<br>(6.8) | **72.2**<br>(3.8) | 58.6<br>(3.6) | 58.8<br>(5.9) | **71.8**<br>(3.8) |
| BALB/C | 54.6<br>(3.8) | 57.5<br>(5.5) | **70.6**<br>(4.8) | 55.1<br>(4.3) | 58.3<br>(6.4) | **71.0**<br>(4.9) | 54.6<br>(3.8) | 57.5<br>(5.5) | **70.6**<br>(4.8) |
| C57BL/6 | 70.7<br>(7.9) | 73.4<br>(7.4) | **74.3**<br>(7.1) | 72.6<br>(8.6) | 75.0<br>(7.5) | **75.4**<br>(7.1) | 70.7<br>(7.9) | 73.4<br>(7.4) | **74.3**<br>(7.1) |
| IV | 55.2<br>(4.0) | 60.4<br>(6.1) | **72.4**<br>(4.0) | 55.8<br>(4.4) | 61.2<br>(6.5) | **72.8**<br>(4.1) | 55.2<br>(4.0) | 60.4<br>(6.1) | **72.4**<br>(4.0) |
| H1N1 | 58.7<br>(6.0) | 59.2<br>(6.3) | **65.0**<br>(7.5) | 61.8<br>(8.0) | 61.9<br>(8.1) | **65.8**<br>(7.6) | 58.7<br>(6.0) | 59.2<br>(6.3) | **65.0**<br>(7.5) |
| H3N2 | 72.1<br>(9.2) | 80.7<br>(11.5) | **84.4**<br>(8.4) | 79.4<br>(8.8) | 84.1<br>(9.7) | **86.5**<br>(7.4) | 72.1<br>(9.2) | 80.7<br>(11.5) | **84.4**<br>(8.4) |
| H5N1 | 57.3<br>(6.4) | 64.9<br>(8.1) | **72.4**<br>(6.9) | 62.1<br>(10.6) | 67.2<br>(8.8) | **73.3**<br>(7.3) | 57.3<br>(6.4) | 64.9<br>(8.1) | **72.4**<br>(6.9) |
| **Three-class virulence grouping** |  |  |  |  |  |  |  |  |  |
| MIV | 45.7<br>(2.6) | 44.5<br>(3.4) | **60.2**<br>(3.0) | 46.6<br>(3.1) | 52.8<br>(5.3) | **60.3**<br>(2.9) | 45.7<br>(2.6) | 44.5<br>(3.4) | **60.2**<br>(3.0) |
| BALB/C | 39.8<br>(3.5) | 42.1<br>(4.2) | **55.4**<br>(3.5) | 40.7<br>(4.8) | 49.1<br>(6.9) | **55.5**<br>(3.5) | 39.8<br>(3.5) | 42.1<br>(4.2) | **55.4**<br>(3.5) |
| C57BL/6 | 60.4<br>(5.8) | 61.9<br>(7.2) | **66.6**<br>(7.5) | 65.6<br>(7.6) | 66.3<br>(7.1) | **68.6**<br>(7.8) | 60.4<br>(5.8) | 61.9<br>(7.2) | **66.6**<br>(7.5) |
| IV | 42.1<br>(3.2) | 42.5<br>(3.3) | **56.3**<br>(3.5) | 43.4<br>(4.4) | 47.9<br>(6.5) | **56.6**<br>(3.5) | 42.1<br>(3.2) | 42.5<br>(3.3) | **56.3**<br>(3.5) |
| H1N1 | 43.3<br>(5.0) | 44.0<br>(7.1) | **54.6**<br>(6.6) | 48.4<br>(8.2) | 50.3<br>(9.7) | **55.5**<br>(7.0) | 43.3<br>(5.0) | 44.0<br>(7.1) | **54.6**<br>(6.6) |
| H3N2 | 47.9<br>(8.9) | 43.0<br>(9.5) | **60.9**<br>(11.7) | 61.4<br>(17.1) | 59.3<br>(14.6) | **64.4**<br>(13.6) | 47.9<br>(8.9) | 43.0<br>(9.5) | **60.9**<br>(11.7) |
| H5N1 | 38.0<br>(5.8) | 42.1<br>(6.9) | **54.0**<br>(7.5) | 39.7<br>(8.6) | 47.6<br>(10.6) | **55.1**<br>(7.8) | 38.0<br>(5.8) | 42.1<br>(6.9) | **54.0**<br>(7.5) |

Table 3 also shows that incorporating host information improved the accuracy of three-class virulence grouping but not for two-class virulence grouping – the mean accuracies of PART models on three-class MIV and IV datasets were 60.2% and 56.3%, respectively, but they were about the same for two-class virulence grouping, i.e., 71.8% for MIV dataset and 72.4% for IV dataset. Furthermore, when a specific host strain was considered, we can see that a rule-based model was easier to learn from C57BL/6 dataset than BALB/C dataset; and when a specific IAV subtype was considered, H3N2 dataset was easier to learn than H1N1 and H5N1 datasets. However, it ought to be noted that the standard deviations for C57BL/6 and H3N2 datasets were higher than the rest, and that aggregating all mouse and/or virus strains gave the smallest standard deviation while keeping accuracy competitive.

The accuracy distribution per learning algorithm per input dataset derived from MIV and IV datasets over 100 models learned independently is shown in Fig. 4, while the accuracy distribution per learning algorithm per input dataset derived from BALB/C, C57BL/6, H1N1 and H3N2 is shown in **Fig. S1** and **Fig. S2**. Once again, we can observe that PART models often outperformed OneR and JRip, and OneR occasionally outperformed JRip. Of interest, models trained on input dataset containing concatenated protein alignments were often better than the ones trained on input containing an alignment of a particular type of IAV protein. Nonetheless, models trained on a particular protein alignment usually achieved averaged accuracies significantly higher than those given by zero rules. The accuracies of models based on alignment of PB2 and/or HA were usually higher than the accuracies of models based on alignment of other proteins. For some cases, the models based only on PB2 or HA could even achieve accuracies as good as those given by the models based on concatenated protein alignments (see the accuracies of models based on PB2 for two-class and three-class H3N2 datasets, PB2 for two-class H5N1 dataset, and HA for two-class H1N1 dataset in **Fig. S2**).

**Fig. 4.**
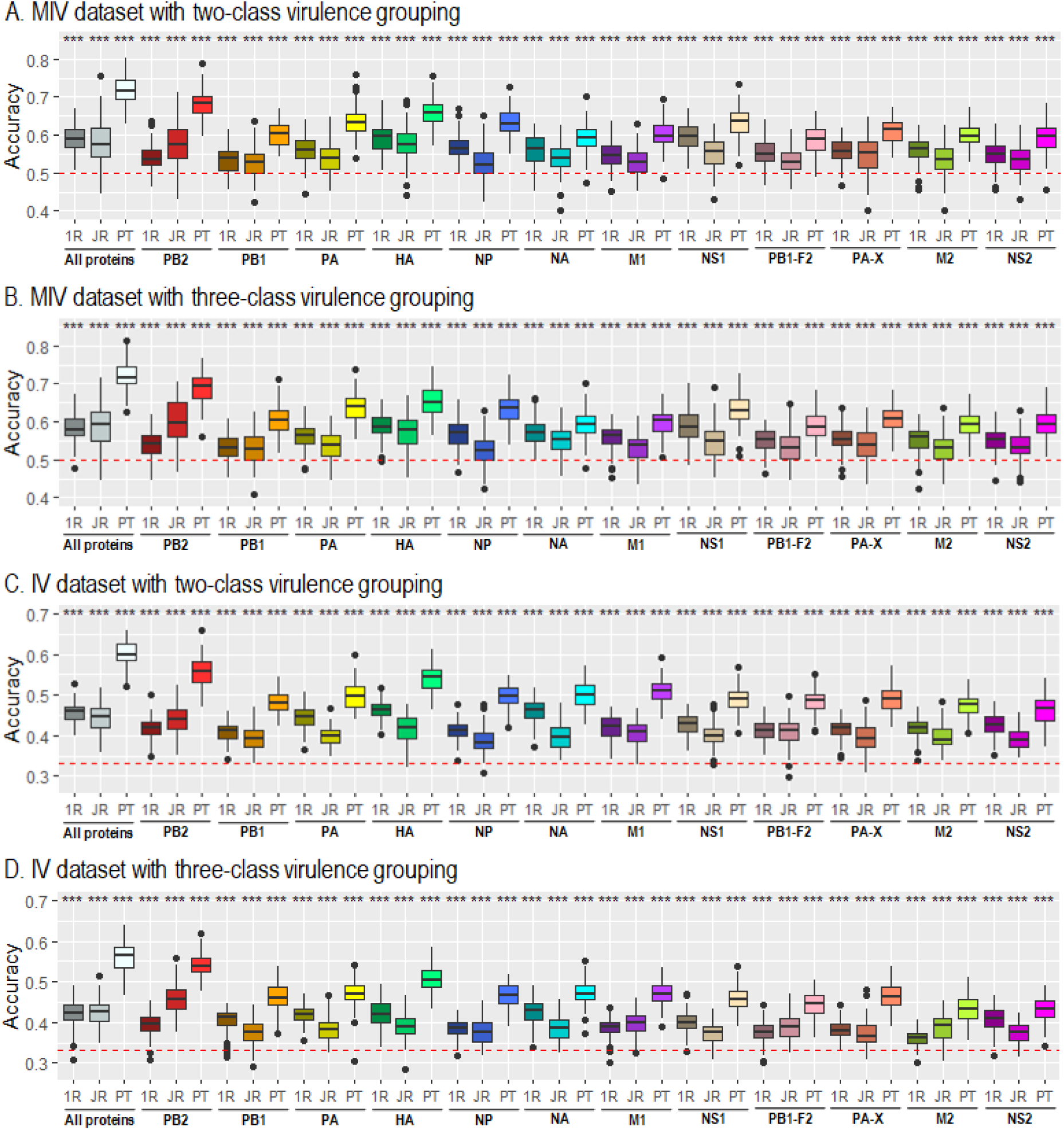
Accuracy distribution of 100 models learned independently from two-class and three-class MIV (A and B, respectively) and IV (C and D, respectively) datasets using OneR, JRip and PART. The datasets contain either the concatenated alignments or an individual alignment of IAV proteins. Wilcoxon signed-rank sum test is used to test the null hypothesis that the median of the accuracy is equal to the accuracy of zero rule learner (represented by the red dashed horizontal line). The level of significance of each test is flagged by the stars: * adjusted p-value <0.05, ** adjusted p-value <0.01 and *** adjusted p-value <0.001.

Finally, we noted that RF models did not outperform PART models. In about 50% of the cases, PART even gave significantly better accuracies than RF (**Fig. S3**). Nonetheless, the site importance ranking output by RF could provide valuable insights and hence, RF models were further explored.

### Top sites and synergy between sites for IAV virulence

As the performance of the models generated by a specific learning algorithm varied from one independent learning to another, the models themselves tended to vary a lot. This demonstrated the influence of selected training data. Hence, rather than inspecting the model one by one, it is more interesting to investigate individual sites that were frequently included in learned models or considered to have more impacts in the models. For this, the OneR’s single site model and RF’s site importance ranking naturally suit the purpose. For JRip and PART, we calculated the averaged contribution of each site to the accuracy of learned models. Table 4 summarizes the sites selected by OneR (ordered by their frequency; sites that were selected once are not shown), top 20 sites by JRip and PART (ordered by their averaged contribution to the accuracy of learned models), and top 20 influential sites by RF (ordered by the averaged mean decrease in accuracy) following 100 independent learnings from both two-class and three-class IV datasets containing concatenated protein alignments.

**Table 4.**
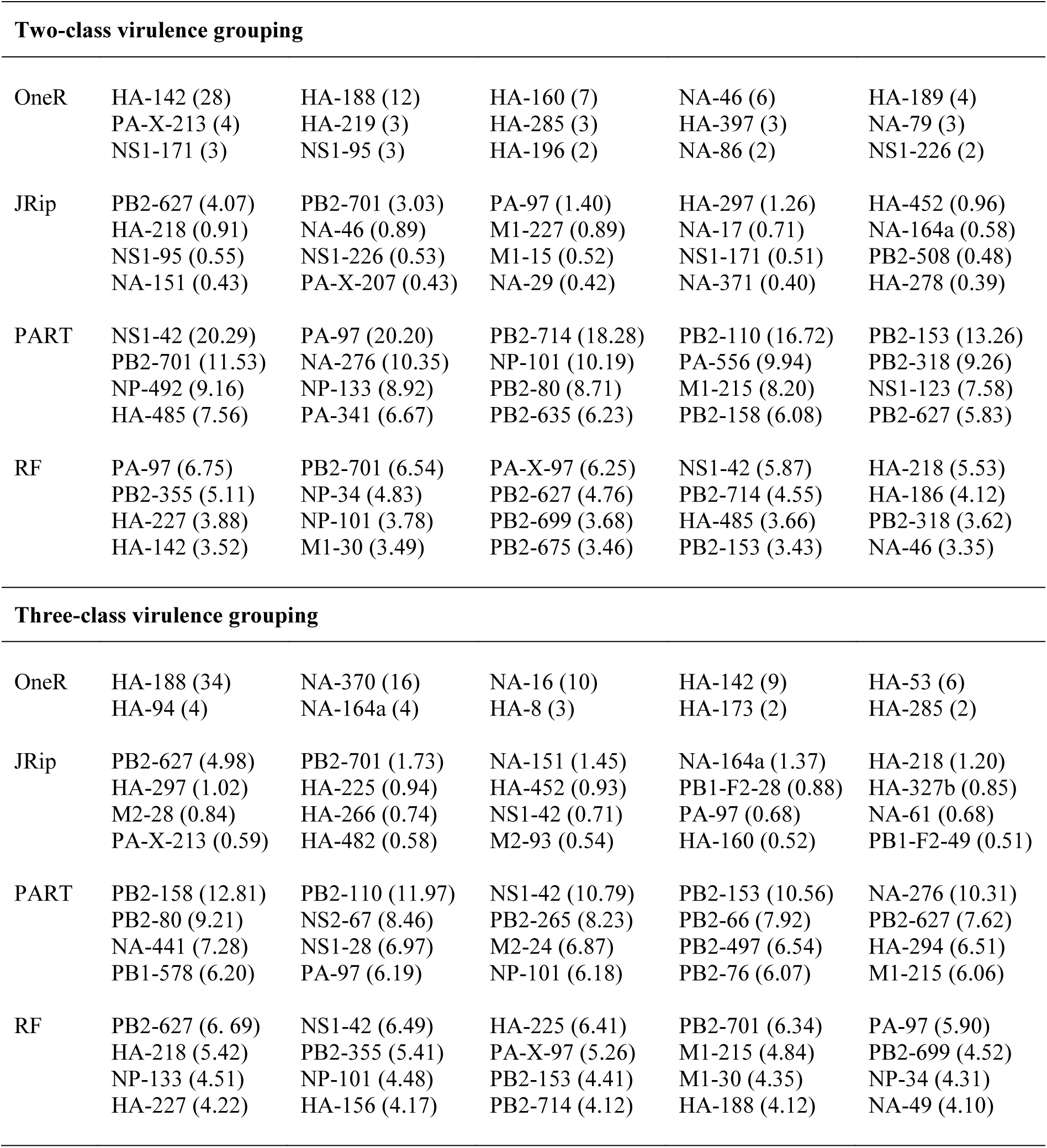
Top sites for modelling IAV virulence based on models generated from two-class and three-class IV datasets. For OneR, the numbers in brackets are the frequency of the corresponding site being selected in the models; for JRip and PART, they are the averaged contribution of the corresponding site to accuracy (in percent); and for random forest (RF), they are the averaged mean decrease in accuracy attributed to the corresponding site. Each number was calculated following 100 independent learnings from two-class or three-class IV dataset. For OneR, only sites with frequency >1 are shown, while for JRip, PART and RF, only top 20 sites are shown.

Overall, for the top sites in Table 4, OneR and JRip preferred sites in HA and NA, PART had a high preference towards sites in PB2, and RF pointed out more sites in PB2 and HA were important. In terms of their consistency in selecting sites for two-class and three-class virulence models, RF was the most consistent (15 shared sites), followed by PART (10 shared sites), JRip (8 shared sites) and finally OneR (only 4 sites). Furthermore, no site was shared by all four learners for either two-class or three-class virulence grouping; but there were few sites shared by combinations of three learners: PB2-627, PB2-701, PA-97 and NA-46 for two-class virulence grouping, and PB2-627, PA-97 and NS1-42 for three-class virulence grouping.

In addition to analyzing individual sites, it is also interesting to investigate the synergy between sites that determine IAV virulence. The rule-based models given by JRip and PART serve this purpose, but here we limit to PART models that gave the highest accuracy. For this, in similar way to the identification of top individual sites, we extracted the averaged contribution of each pair of sites appearing in each rule in PART models to the overall accuracy. The synergistic networks arising from top 50 site pairs in PART models learned from two-class and three-class IV datasets are shown in Fig. 5A and 5B, respectively. As shown, the sites in both cases are interestingly fully connected and mainly involved sites in PB2. Top 4 sites that had highest degree (number of connections) for two-class virulence grouping included PB2-714 (degree = 14), PA-97 (13), NS1-42 (10) and PB2-701 (7), and interestingly, the pairing between top two sites PB2-714 and PA-97 had the highest contribution to accuracy. On the other hand, sites that have highest degree for three-class virulence grouping included PB2-110 (15), PB2-158 (13), NS1-42 (10) and PB2-153 (9), and the pairing between PB2-153 and NS1-42 had the highest contribution to accuracy.

**Fig. 5.**
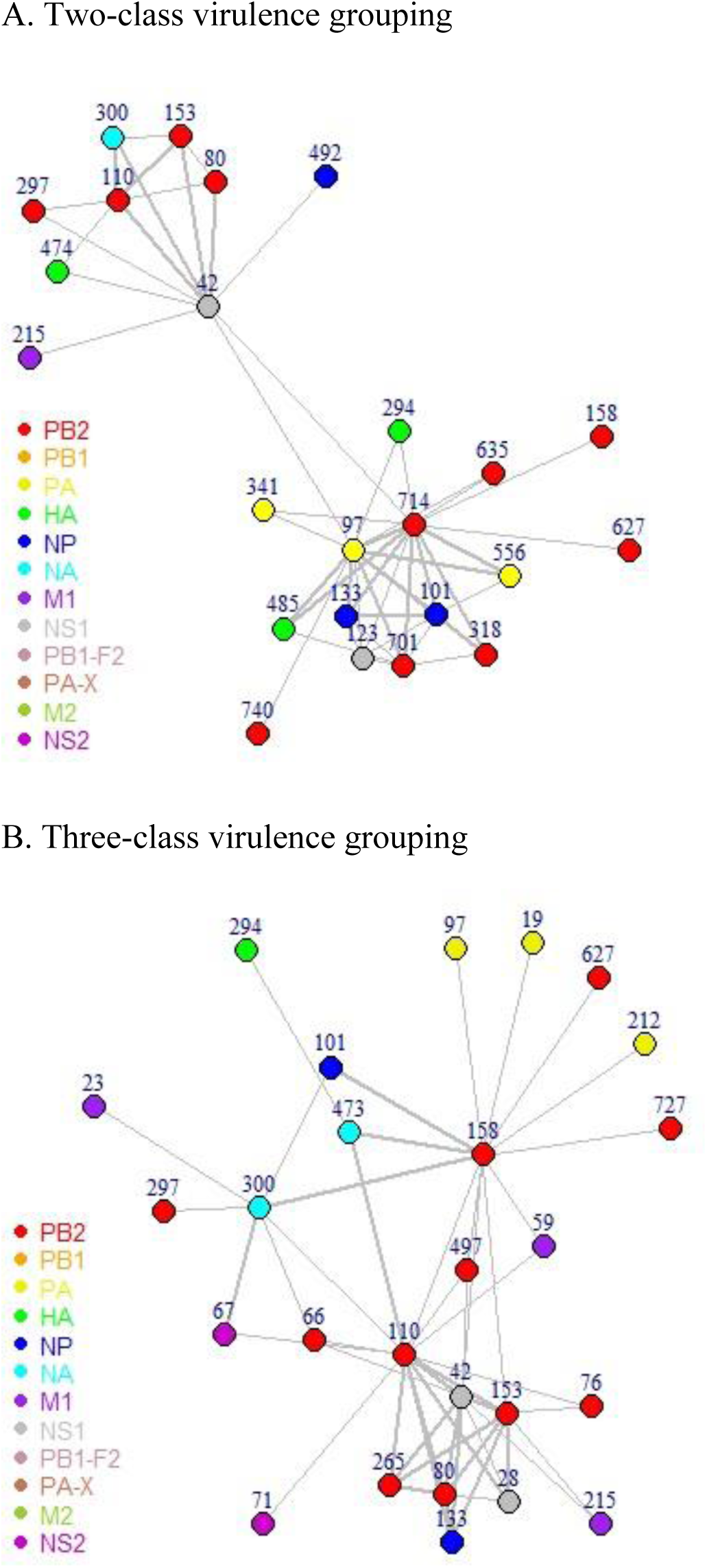
Synergistic graphs between protein sites that are based on models generated by PART from (A) two-class and (B) three-class IV datasets containing concatenated alignments of IAV proteins. The nodes in the graph are the sites in IAV proteins – the proteins are encoded by color and the site numbers are written above the nodes. Two sites are connected by an edge if they appear in the top 50 site pairs that have the highest contribution to accuracy. The thickness of an edge indicates the level of contribution of the corresponding site pair to accuracy of PART models.

## Discussions

In this influenza study, we systematically and extensively searched literature, collected infection records involving specific mouse and IAV strains, noted their virulence, classified the virulence level (the various units of infection dose were assumed to measure the same quantity and the MLD50 thresholds 10E3.0 and 10E6.0 for virulence classification follow the thresholds used by WHO when the infection doses measured with EID50 [25]), and obtained related IAV proteins in order to develop predictive virulence models of IAV infections. Furthermore, we proposed a number of procedures to tackle various missing data. For virulence, the MLD50 value is the ultimate information we looked for; but in its absence, time series of weight loss or percentage of survival of infected mice were utilized to infer the lower or upper bound of MLD50 and subsequently, to label the virulence class. For IAV genomes, when the genomes were incomplete or contained partial sequences, extrapolation was performed using the closest genome relative identified with BLAST. These pre-processing works were done manually and ambiguity occasionally occurred. Hence, caution must be taken when dealing with the datasets and improvement in the pre-processing approach may be considered for future works. Alternatively, efforts in improving the current practice of storing IAV virulence information by research community such that it eases its reusability could be encouraged, e.g., by creating an online database that accepts submissions of IAV virulence related data and provides high quality tables or figures of their input data that can be added into their manuscript.

Despite the limitations of the datasets due to the ways in handling missing MLD50, partial sequences and incomplete genomes, and also a recent critic of using LD50 as a virulence measure [26], the models learned from the datasets could provide insights about IAV virulence across mouse and virus strains. Rule-based models were chosen since their output can be easily interpreted and are congruent with the current practice in investigating IAV virulence experimentally. Three rule-based learning approaches were employed: OneR, JRip and PART. OneR approach outputs a single site model that gives the highest accuracy [27]; JRip and PART considers multiple sites and they construct a set of decision rules using different strategy. While JRip mainly uses separate-and-conquer algorithms [28], PART combines separate-and-conquer strategy and partial decision trees [29]. For a comparison in the performance, we also explored the RF approach [30] in modelling IAV virulence.

For the models and their performance, we first noted that OneR mainly selected sites in HA and NA for its single site models, and the OneR models could give significantly better averaged accuracies than the zero rule models. Among the sites, some have known functions while some others are not yet characterized. For example, site 188 is known to be located at the helix 190 that surrounds the receptor-binding site in the HA protein and thus it affects host specificity [31], while site 142 has not yet been well studied even though it was frequently selected as the top OneR classifier. Nonetheless, JRip and PART generated multiple site models that almost always gave better accuracies than OneR models for any specific IAV protein. Of interest, PART not only outperformed OneR and JRip, but also RF in 50% of the tested cases. Moreover, higher accuracy generally could be achieved by PART when considering all IAV proteins at once. These results demonstrate a synergistic between sites within a single protein and sites in different proteins; in other words, the polygenic nature of IAV virulence in mice. This is consistent with the observations from various experimental studies, such as the ones that demonstrate intra-protein synergy in PB2 [32–37], PA [15], and NS1 [38, 39], and inter-protein synergy that involves combinations of PB2, PB1, PA, HA or NA [16, 40-46].

Further inspection on PART models across different IAV strains using IV dataset revealed that although HA had many more sites correlated with virulence, PB2 seemed to play more important role in determining IAV virulence. This was in agreement with the RF’s site importance ranking. In terms of their accuracy, PART models based on PB2 alone could compete against or were even better than PART models based on HA; except when modelling the virulence of H1N1 virus alone, PART models based on HA from two-class datasets were more superior (see **Fig. S2A**). Moreover, PART models based on all IAV proteins have a high preference towards sites in PB2, and many sites in PB2 were also considered as the most important features for RF models (Table 4). Fig. 5 that shows synergistic graphs for two-class and three-class virulence grouping further clearly demonstrate this. Investigations on MIV dataset and datasets for specific IAV or mouse strain also revealed the dominance of PB2 in most of the cases (data not shown). When sites in PB2 did not dominate, the sites in HA dominated, such as in the case for two-class H1N1 dataset.

The critical role of PB2 in determining virulence in mice have been indeed highlighted for various strains, including H3N2 [44, 47], H5N1 [32-34, 48, 49], H5N8 [36, 50], H7N9 [51–55], H9N2 [35, 37, 55, 56] and H10N8 [55]. Among the top 20 sites in PB2 for PART models, sites 627 and 701 have been repeatedly shown to affect IAV virulence in mammals including mice. Site 627 is considered critical for efficient replication, while site 701 influences polymerase activity via its interaction with the nuclear import factor importin α that mediates the transport of proteins into nucleus [57]. Other top sites in PB2 are also known to contribute to virulence. For examples, site 714 (top 20 for two-class IV dataset) influences replication efficiency and IAV virulence in mice in combination with site 701 [33, 58, 59]; site 66 (top 20 for three-class IV dataset) sets a prerequisite for acquiring virulence [60]; and site 158 (top 20 for two-class and three-class IV dataset; specifically, top one for three-class) strongly influences the virulence of both pandemic H1N1 and H5 influenza viruses in mice [61]. Experimental evidence for the contribution of other top sites in PB2 to virulence, e.g., sites 80, 110 and 153, are still none to our knowledge. On the other hand, some other sites not in the top list have been shown to play a role in dictating virulence, e.g., sites 147, 339 and 588 that can synergize to give rise a higher level of virulence [34].

Next, the synergistic graph for two-class virulence grouping interestingly presents a clustering of two subgraphs for sites in PART virulence models, with sites PB2-714, PA-97 and NS1-42 act as a bottleneck (a node with high betweenness centrality, i.e., having many shortest paths going through it) connecting the two subgraphs. Furthermore, when three-class was considered, the synergistic graph containing top site pairs concentrated and expanded in the subnetwork that included sites PB2-80, PB2-110, PB2-153, PB2-297, NA-300, NS1-42, and M1-215. This may indicate a greater role of these sites in sensitizing the virulence level of IAV infections. For example, site 42 within the RNA-binding domain of NS1 influences the capability of the protein in binding double-stranded RNA and it determines the degree of pathogenicity in mice [62]. This site also influences the activation of IRF3 and regulation of host interferon response, which subsequently influences the efficiency of viral replication [63]. Another site that has been experimentally explored is site 215 in M1, which also contributes to the degree of IAV virulence [64].

Overall, PART, with its approach that combines separate-and-conquer strategy and partial decision tree, has been a suitable method to generate sequence-based virulence models that are not only considerably good in performance, but also provides interpretable information. But here, rather than relying on a single model developed from a single training dataset, the information was extracted from 100 models learned independently from different training datasets. While bias due to imbalanced classes were resolved by under-sampling to obtain balanced classes, the iterations might help reducing bias due to over-sampling of a particular mouse or IAV strain. Furthermore, we also noted from the confusion matrix that PART models tended to misclassify the avirulent (or less virulent) strains as virulent (or more virulent) ones rather than misclassify the virulent (more virulent) strains as avirulent (or less virulent) ones. In practice, this is preferred since classifying the virulent (more virulent) strains as avirulent (less virulent) ones is a worse decision that can cost lives.

In terms of their accuracy, PART models achieved moderate performance for various datasets being learned. The average accuracy over 100 models ranged between 65.0% and 84.4% (15.0% - 34.4% above baseline) for two-class datasets that utilized all IAV proteins, and between 54.0% and 66.6% (20.7% - 33.3% above baseline) for three-class datasets (see Table 3). Learning from subsets of specific mouse or IAV strains revealed that some strains were easier while others were harder to learn. Of interest, while the average accuracies were relatively the same for full two-class datasets regardless the host information was included or not, some significant improvement (3.9% in increase of accuracy) was observed when incorporating host information for full three-class dataset. Thus, using learning approaches that further incorporate host information shall be encouraged, especially since several laboratory experiments have demonstrated the importance of host genetic backgrounds in determining IAV virulence [65–71]. In particular, with the availability of genomes and proteomes of various mouse strains, sophisticated methods that are based on host-pathogen protein-protein interactions might be of interest. If successful, an implementation of such methods may be translated to human cases and other diseases to improve our understanding about disease mechanisms, establish a foundation for future personalized medicine, and provide a better surveillance. Nevertheless, the development of the approaches will be more fruitful if there is a significant increase in the number of influenza experiments carried out with mouse and IAV strains that are still limited in their number of studies.

In summary, we have developed benchmark datasets for IAV virulence and explored rule-based and RF approaches for modelling IAV virulence. To our knowledge, the datasets have been the biggest aggregation of IAV infections in mice, and the number of the infection records can still grow. The creation of these benchmark datasets will be beneficial for further understanding the molecular principles underlying influenza mechanisms since mice have been a major animal model for influenza. In the current study, we utilized the datasets to assess predictabilities of IAV virulence for specific and across mouse and IAV strains, and identify top proteins sites and synergy between protein sites that contribute to IAV virulence. Overall, our study confirmed the polygenic nature of IAV virulence, with several sites in PB2 playing more dominant roles. Not only sites that are well known as IAV virulence markers, e.g. 627, 701 and 714, but also some other sites in PB2 not yet known influencing virulence were identified. Nonetheless, modelling virulence is in fact a very challenging problem due to the nature of complex interactions that underlie the phenotype, which involve not only viral factors, but also host factors. Hence, future works shall incorporate more host information, especially the host proteomic data that now widely available for various mouse strains. Applying different machine learning approaches and protein features, and posing virulence modelling as a regression problem that predicts LD50 shall also be considered.

## Acknowledgements

The project is supported by AcRF Tier 2 Grant MOE2014-T2-2-023, Ministry of Education, Singapore.

