## Supplements for "Rule-based meta-analysis reveals the major role of PB2 in influencing influenza A virus virulence in mice"

**Table S1.** Initial dataset for influenza A virus (IAV) infections in mice with virulence information.

| Reference | Host strain | Influenza strain | LD50 point estimate | LD50 lower bound | LD50 upper bound | Infection unit | Notes on LD50 | Two-class virulence level | Three-class virulence level |
|---|---|---|---|---|---|---|---|---|---|
| Ainai *et al*., 2015 [1] | BALB/C | A/Narita/1/2009/MDCK(H1N1) | | 5 | | TCID50 | Value given | Avirulent | LOW |
| | BALB/C | A/Narita/1/2009/MDCK15(H1N1) | | 6 | | TCID50 | Value given | Avirulent | LOW |
| | BALB/C | A/Narita/1/2009/Egg(H1N1) | 5.2 | | | TCID50 | Value given | Virulent | INTERMEDIATE |
| | BALB/C | A/Narita/1/2009/Egg15(H1N1) | 5.9 | | | TCID50 | Value given | Virulent | INTERMEDIATE |
| | BALB/C | A/Narita/1/2009/Mouse15(H1N1) | 3.91 | | | TCID50 | Value given | Virulent | INTERMEDIATE |
| Belser *et al*., 2007 [2] | BALB/C | A/Netherlands/219/2003(H7N7) | 2.5 | | | EID50 | Value given | Virulent | HIGH |
| | BALB/C | A/Netherlands/230/2003(H7N7) | | 7 | | EID50 | Value given | Avirulent | LOW |
| | BALB/C | A/chicken/Netherlands/1/2003(H7N7) | | 7 | | EID50 | Value given | Avirulent | LOW |
| | BALB/C | A/NewYork/107/2003(H7N2) | | 7 | | EID50 | Value given | Avirulent | LOW |
| | BALB/C | A/chicken/CT/260413-2/2003(H7N2) | | 7 | | EID50 | Value given | Avirulent | LOW |
| Belser *et al*., 2010 [3] | BALB/C | A/California/04/2009(H1N1) | | 6 | | EID50 | Value given | Avirulent | LOW |
| | BALB/C | A/Texas/15/2009(H1N1) | | 6 | | EID50 | Value given | Avirulent | LOW |
| | BALB/C | A/Mexico/4108/2009(H1N1) | | 6 | | EID50 | Value given | Avirulent | LOW |
| | BALB/C | A/Mexico/4482/2009(H1N1) | | 6 | | EID50 | Value given | Avirulent | LOW |
| | BALB/C | A/Mexico/InDRE4487/2009(H1N1) | | 6 | | EID50 | Value given | Avirulent | LOW |
| | BALB/C | A/NewJersey/8/1976(H1N1) | | 6 | | EID50 | Value given | Avirulent | LOW |
| | BALB/C | A/Ohio/02/2007(H1N1) | 5.8 | | | EID50 | Value given | Virulent | INTERMEDIATE |
| | BALB/C | A/SouthCarolina/1/1918(H1N1) | 3.5 | | | EID50 | Value given | Virulent | INTERMEDIATE |
| | BALB/C | A/VietNam/1203/2004(H5N1) | 1.3 | | | EID50 | Value given | Virulent | HIGH |
| Bi *et al*., 2015 [4] | BALB/C | rA/Anhui/1/2013(H7N9) | 7.83 | | | EID50 | Value given | Avirulent | LOW |
| | BALB/C | rA/chicken/Shandong/lx1023/2007(H9N2) | | 9.37 | | EID50 | Value given | Avirulent | LOW |
| | BALB/C | rA/SH1023(2345678)/AH1(1)(H9N2) | 5.5 | | | EID50 | Value given | Virulent | INTERMEDIATE |
| | BALB/C | rA/SH1023(1345678)/AH1(2)(H9N2) | | 9.37 | | EID50 | Value given | Avirulent | LOW |
| | BALB/C | rA/SH1023(1245678)/AH1(3)(H9N2) | | 10.45 | | EID50 | Value given | Avirulent | LOW |
| | BALB/C | rA/SH1023(1234678)/AH1(5)(H9N2) | | 9.95 | | EID50 | Value given | Avirulent | LOW |
| | BALB/C | rA/SH1023(1234568)/AH1(7)(H9N2) | 8.48 | | | EID50 | Value given | Avirulent | LOW |
| | BALB/C | rA/SH1023(1234567)/AH1(8)(H9N2) | | 9.7 | | EID50 | Value given | Avirulent | LOW |
| | BALB/C | rA/SH1023(2345678)/AH1(1)-627E(H9N2) | 7.12 | | | EID50 | Value given | Avirulent | LOW |
| | BALB/C | rA/Anhui/1/PB2-627E/2013(H7N9) | | 7.62 | | EID50 | Value given | Avirulent | LOW |
| Blazejewska *et al*., 2011 [5] | C57BL/6 | A/PR8M/1934(H1N1) | | 3.3 | | PFU | LD50_LB or LD50_UB is based on survival rate | Virulent | INTERMEDIATE |

| | | | | | | | | | |
|---|---|---|---|---|---|---|---|---|---|
| | C57BL/6 | A/PR8F/1934(H1N1) | | | 3.3 | PFU | LD50_LB or LD50_UB is based on survival rate | Virulent | HIGH |
| | C57BL/6 | A/hvPR8/1934(H1N1) | | | 3.3 | PFU | LD50_LB or LD50_UB is based on survival rate | Virulent | HIGH |
| | C57BL/6 | rA/PR8M(1235678)/PR8F(4)(H1N1) | | | 3.3 | PFU | LD50_LB or LD50_UB is based on survival rate | Virulent | HIGH |
| | DBA/2 | A/PR8M/1934(H1N1) | | | 3.3 | PFU | LD50_LB or LD50_UB is based on survival rate | Virulent | HIGH |
| | DBA/2 | A/PR8F/1934(H1N1) | | | 3.3 | PFU | LD50_LB or LD50_UB is based on survival rate | Virulent | HIGH |
| | DBA/2 | A/hvPR8/1934(H1N1) | | | 3.3 | PFU | LD50_LB or LD50_UB is based on survival rate | Virulent | HIGH |
| Bodewes *et al*., 2011 [6] | C57BL/6 | rA/PR8(1234678)/NL94(5)(H1N1) | | 3 | | TCID50 | LD50_LB or LD50_UB is based on weight loss | Avirulent | LOW |
| | C57BL/6 | rA/PR8(1234678)/NL94(5)/G384R(H1N1) | | 3 | | TCID50 | LD50_LB or LD50_UB is based on weight loss | Virulent | INTERMEDIATE |
| Casalegno *et al*., 2014 [7] | BALB/C | A/Lyon/969/2009(H1N1) | 3.2 | | | TCID50 | Value given; It is written as MID50 in the paper | Virulent | INTERMEDIATE |
| | BALB/C | A/StEtienne/1139/2010(H1N1) | 4.2 | | | TCID50 | Value given; It is written as MID50 in the paper | Virulent | INTERMEDIATE |
| | BALB/C | A/StEtienne/1691/2009(H1N1) | 2.7 | | | TCID50 | Value given; It is written as MID50 in the paper | Virulent | HIGH |
| | BALB/C | A/Limoges/1159/2010(H1N1) | 3.15 | | | TCID50 | Value given; It is written as MID50 in the paper | Virulent | INTERMEDIATE |
| | BALB/C | A/LaReunion/803/2010(H1N1) | 2.95 | | | TCID50 | Value given; It is written as MID50 in the paper | Virulent | HIGH |
| | BALB/C | A/Lyon/52.16/2010(H1N1) | 2.7 | | | TCID50 | Value given; It is written as MID50 in the paper | Virulent | HIGH |
| | BALB/C | A/Lyon/1.12/2011(H1N1) | 3 | | | TCID50 | Value given; It is written as MID50 in the paper | Virulent | HIGH |
| Chen *et al*., 2007 [8] | BALB/C | rA/HongKong/483/1997(H5N1) | 1.9 | | | EID50 | Value given | Virulent | HIGH |
| | BALB/C | rA/HongKong/486/1997(H5N1) | 5.98 | | | EID50 | Value given | Virulent | INTERMEDIATE |
| | BALB/C | rA/HK486/15mts(H5N1) | 4.78 | | | EID50 | Value given | Virulent | INTERMEDIATE |
| | BALB/C | rA/HK486/5mts(H5N1) | 4.78 | | | EID50 | Value given | Virulent | INTERMEDIATE |
| | BALB/C | rA/HK486/PB2-6mts(H5N1) | 6 | | | EID50 | Value given | Virulent | INTERMEDIATE |
| | BALB/C | rA/HK486/M-3mts(H5N1) | 6.17 | | | EID50 | Value given | Avirulent | LOW |
| | BALB/C | rA/HK486/NA-3mts(H5N1) | 6.25 | | | EID50 | Value given | Avirulent | LOW |
| | BALB/C | rA/HK486/PB1-3mts(H5N1) | 7.83 | | | EID50 | Value given | Avirulent | LOW |
| | BALB/C | rA/HK486/PB2-627K(H5N1) | 2.25 | | | EID50 | Value given | Virulent | HIGH |
| | BALB/C | rA/HK483(2345678)/HK486(1)(H5N1) | 6.25 | | | EID50 | Value given | Avirulent | LOW |
| | BALB/C | rA/HK486(123578)/HK483(46)(H5N1) | 3.9 | | | EID50 | Value given | Virulent | INTERMEDIATE |
| | BALB/C | rA/HK486(1235678)/HK483(4)(H5N1) | 4.88 | | | EID50 | Value given | Virulent | INTERMEDIATE |
| | BALB/C | rA/HK486(1234578)/HK483(6)(H5N1) | 5.75 | | | EID50 | Value given | Virulent | INTERMEDIATE |
| Choi *et al*., 2017 [9] | C57BL/6 | A/mallard/Korea/W452/2014(H5N8) | | 4 | | PFU | LD50_LB or LD50_UB is based on survival rate | Avirulent | LOW |
| | C57BL/6 | A/ma452-G1-1/2014(H5N8) | | | 4 | PFU | LD50_LB or LD50_UB is based on survival rate | Virulent | HIGH |
| | C57BL/6 | A/ma452-G3-1/2014(H5N8) | | | 4 | PFU | LD50_LB or LD50_UB is based on survival rate | Virulent | HIGH |
| | C57BL/6 | A/ma452-G3-2/2014(H5N8) | | | 4 | PFU | LD50_LB or LD50_UB is based on survival rate | Virulent | HIGH |
| | C57BL/6 | A/ma452-G4-1/2014(H5N8) | | | 4 | PFU | LD50_LB or LD50_UB is based on survival rate | Virulent | HIGH |
| | C57BL/6 | A/environment/Korea/W468/2014(H5N8) | | 4 | | PFU | LD50_LB or LD50_UB is based on survival rate | Avirulent | LOW |

| | | | | | | | | | |
|---|---|---|---|---|---|---|---|---|---|
| | C57BL/6 | A/ma468-G1-1/2014(H5N8) | | | 4 | PFU | LD50_LB or LD50_UB is based on survival rate | Virulent | HIGH |
| | C57BL/6 | A/ma468-G1-2/2014(H5N8) | | | 4 | PFU | LD50_LB or LD50_UB is based on survival rate | Virulent | HIGH |
| | C57BL/6 | A/ma468-G2-1/2014(H5N8) | | | 4 | PFU | LD50_LB or LD50_UB is based on survival rate | Virulent | HIGH |
| | C57BL/6 | A/ma468-G2-2/2014(H5N8) | | | 4 | PFU | LD50_LB or LD50_UB is based on survival rate | Virulent | HIGH |
| | C57BL/6 | A/ma468-G2-3/2014(H5N8) | | 4 | | PFU | LD50_LB or LD50_UB is based on survival rate | Avirulent | LOW |
| | C57BL/6 | A/ma468-G4-2/2014(H5N8) | | | 4 | PFU | LD50_LB or LD50_UB is based on survival rate | Virulent | HIGH |
| | BALB/C | A/mallard/Korea/W452/2014(H5N8) | 7.5 | | | PFU | Value given | Avirulent | LOW |
| | BALB/C | A/ma452-G1-1/2014(H5N8) | 1 | | | PFU | Value given | Virulent | HIGH |
| | BALB/C | A/ma452-G3-1/2014(H5N8) | 2 | | | PFU | Value given | Virulent | HIGH |
| | BALB/C | A/ma452-G3-2/2014(H5N8) | 0.5 | | | PFU | Value given | Virulent | HIGH |
| | BALB/C | A/ma452-G4-1/2014(H5N8) | 1.3 | | | PFU | Value given | Virulent | HIGH |
| | BALB/C | A/environment/Korea/W468/2014(H5N8) | 7.3 | | | PFU | Value given | Avirulent | LOW |
| | BALB/C | A/ma468-G1-1/2014(H5N8) | 1.7 | | | PFU | Value given | Virulent | HIGH |
| | BALB/C | A/ma468-G1-2/2014(H5N8) | 1 | | | PFU | Value given | Virulent | HIGH |
| | BALB/C | A/ma468-G2-1/2014(H5N8) | 0.7 | | | PFU | Value given | Virulent | HIGH |
| | BALB/C | A/ma468-G2-2/2014(H5N8) | 1.5 | | | PFU | Value given | Virulent | HIGH |
| | BALB/C | A/ma468-G2-3/2014(H5N8) | 4.8 | | | PFU | Value given | Virulent | INTERMEDIATE |
| | BALB/C | A/ma468-G4-2/2014(H5N8) | 0.5 | | | PFU | Value given | Virulent | HIGH |
| Cline *et al*., 2011 [10] | BALB/C | A/California/04/2009(H1N1) | | 5 | | TCID50 | Value given | Virulent | INTERMEDIATE |
| | BALB/C | A/HongKong/483/1997(H5N1) | 1.5 | | | TCID50 | Value given | Virulent | HIGH |
| | BALB/C | rA/CA04(2345678)/HK483(1)(H1N1) | | 3 | | TCID50 | LD50_LB or LD50_UB is based on survival rate | Avirulent | LOW |
| | BALB/C | rA/CA04(1345678)/HK483(2)(H1N1) | | 3 | | TCID50 | LD50_LB or LD50_UB is based on survival rate | Virulent | INTERMEDIATE |
| | BALB/C | rA/CA04(1245678)/HK483(3)(H1N1) | | 3 | | TCID50 | LD50_LB or LD50_UB is based on survival rate | Avirulent | LOW |
| | BALB/C | rA/CA04(1235678)/HK483(4)(H5N1) | 2.5 | | | TCID50 | Value given | Virulent | HIGH |
| | BALB/C | rA/CA04(1234678)/HK483(5)(H1N1) | | 3 | | TCID50 | LD50_LB or LD50_UB is based on survival rate | Avirulent | LOW |
| | BALB/C | rA/CA04(1234578)/HK483(6)(H1N1) | | 3 | | TCID50 | LD50_LB or LD50_UB is based on survival rate | Avirulent | LOW |
| | BALB/C | rA/CA04(1234568)/HK483(7)(H1N1) | | 3 | | TCID50 | LD50_LB or LD50_UB is based on survival rate | Avirulent | LOW |
| | BALB/C | rA/CA04(1234567)/HK483(8)(H1N1) | | 3 | | TCID50 | LD50_LB or LD50_UB is based on survival rate | Virulent | INTERMEDIATE |
| Cox *et al*., 2015 [11] | C57BL/6 | A/PR8/LAIV/1934(H1N1) | 4.5 | | | FFU | Using formula Reed and Muench method | Virulent | INTERMEDIATE |
| | C57BL/6 | A/PR8/11C/1934(H1N1) | | | 3 | FFU | LD50_LB or LD50_UB is based on survival rate | Virulent | HIGH |
| | C57BL/6 | A/PR8/LAIV11C/1934(H1N1) | 5.5 | | | FFU | Using formula Reed and Muench method | Virulent | INTERMEDIATE |
| de Jong *et al*., 2013 [12] | BALB/C | A/chicken/Netherlands/621557/2003(H7N7) | | 6 | | TCID50 | Value given | Avirulent | LOW |
| | BALB/C | maA/chicken/Netherlands/621557/2003(H7N7) | | | 2 | TCID50 | Value given | Virulent | HIGH |
| Driskell *et al*., 2010 [13] | BALB/C | A/ruddyturnstone/Delaware/650625/2002(H6N1) | | 4.4 | | PFU | LD50_LB or LD50_UB is based on weight loss | Avirulent | LOW |

| | | | | | | | | | |
|---|---|---|---|---|---|---|---|---|---|
| Elliott *et al*., 2017 [14] | BALB/C | A/California/07/2009(H1N1) | 4.5 | | | TCID50 | Value given | Virulent | INTERMEDIATE |
| Fan *et al*., 2009 [15] | BALB/C | A/duck/Guangxi/53/2002(H5N1) | 6.5 | | | EID50 | Value given | Avirulent | LOW |
| | BALB/C | A/duck/Fujian/01/2002(H5N1) | 0.5 | | | EID50 | Value given | Virulent | HIGH |
| | BALB/C | rA/duck/Guangxi/53/2002(H5N1) | 6.4 | | | EID50 | Value given | Avirulent | LOW |
| | BALB/C | rA/duck/Fujian/01/2002(H5N1) | 0.9 | | | EID50 | Value given | Virulent | HIGH |
| | BALB/C | rA/DKGX53(2345678)/DKFJ01(1)(H5N1) | 6.4 | | | EID50 | Value given | Avirulent | LOW |
| | BALB/C | rA/DKGX53(1345678)/DKFJ01(2)(H5N1) | 6.4 | | | EID50 | Value given | Avirulent | LOW |
| | BALB/C | rA/DKGX53(1245678)/DKFJ01(3)(H5N1) | 6.4 | | | EID50 | Value given | Avirulent | LOW |
| | BALB/C | rA/DKGX53(1235678)/DKFJ01(4)(H5N1) | 6.4 | | | EID50 | Value given | Avirulent | LOW |
| | BALB/C | rA/DKGX53(1234678)/DKFJ01(5)(H5N1) | 6.4 | | | EID50 | Value given | Avirulent | LOW |
| | BALB/C | rA/DKGX53(1234578)/DKFJ01(6)(H5N1) | 6.4 | | | EID50 | Value given | Avirulent | LOW |
| | BALB/C | rA/DKGX53(1234568)/DKFJ01(7)(H5N1) | 3 | | | EID50 | Value given | Virulent | HIGH |
| | BALB/C | rA/DKGX53(1234567)/DKFJ01(8)(H5N1) | 6.2 | | | EID50 | Value given | Avirulent | LOW |
| | BALB/C | rA/DKFJ01(2345678)/DKGX53(1)(H5N1) | 2.9 | | | EID50 | Value given | Virulent | HIGH |
| | BALB/C | rA/DKFJ01(1345678)/DKGX53(2)(H5N1) | 2.5 | | | EID50 | Value given | Virulent | HIGH |
| | BALB/C | rA/DKFJ01(1245678)/DKGX53(3)(H5N1) | 2.5 | | | EID50 | Value given | Virulent | HIGH |
| | BALB/C | rA/DKFJ01(1235678)/DKGX53(4)(H5N1) | 1.5 | | | EID50 | Value given | Virulent | HIGH |
| | BALB/C | rA/DKFJ01(1234678)/DKGX53(5)(H5N1) | 1.5 | | | EID50 | Value given | Virulent | HIGH |
| | BALB/C | rA/DKFJ01(1234578)/DKGX53(6)(H5N1) | 0.9 | | | EID50 | Value given | Virulent | HIGH |
| | BALB/C | rA/DKFJ01(1234568)/DKGX53(7)(H5N1) | 4.5 | | | EID50 | Value given | Virulent | INTERMEDIATE |
| | BALB/C | rA/DKFJ01(1234567)/DKGX53(8)(H5N1) | 0.9 | | | EID50 | Value given | Virulent | HIGH |
| | BALB/C | A/DKGX53/M1N30D(H5N1) | 5.8 | | | EID50 | Value given | Virulent | INTERMEDIATE |
| | BALB/C | A/DKGX53/M1S126G(H5N1) | 6.4 | | | EID50 | Value given | Avirulent | LOW |
| | BALB/C | A/DKGX53/M1T215A(H5N1) | 4.6 | | | EID50 | Value given | Virulent | INTERMEDIATE |
| | BALB/C | A/DKGX53/M1N30DT215A(H5N1) | 3.2 | | | EID50 | Value given | Virulent | INTERMEDIATE |
| | BALB/C | A/DKGX53/M1S126GT215A(H5N1) | 4.3 | | | EID50 | Value given | Virulent | INTERMEDIATE |
| | BALB/C | A/DKFJ01/M1D30NA215T(H5N1) | 4.8 | | | EID50 | Value given | Virulent | INTERMEDIATE |
| | BALB/C | A/DKFJ01/M1D30N(H5N1) | 2.5 | | | EID50 | Value given | Virulent | HIGH |
| | BALB/C | A/DKFJ01/M1A215T(H5N1) | 3.3 | | | EID50 | Value given | Virulent | INTERMEDIATE |
| Ferraris *et al*., 2012 [16] | BALB/C | A/Lyon/969/2009(H1N1) | | 6 | | TCID50 | Value given | Avirulent | LOW |
| | BALB/C | A/Lyon/1337/2007(H1N1) | | 6 | | TCID50 | Value given | Avirulent | LOW |
| | BALB/C | rA/L969(1234578)/L1337(6)(H1N1) | | 6 | | TCID50 | Value given | Avirulent | LOW |
| | BALB/C | rA/L969(123478)/L1337(56)(H1N1) | | 6 | | TCID50 | Value given | Avirulent | LOW |
| | BALB/C | rA/L969(123578)/L1337(46)(H1N1) | | 6 | | TCID50 | Value given | Avirulent | LOW |

| | | | | | | | | | |
|---|---|---|---|---|---|---|---|---|---|
| | BALB/C | A/Lyon/48.425/2009(H1N1) | | 6 | | TCID50 | Value given | Avirulent | LOW |
| Ferraris *et al*., 2018 [17] | BALB/C | A/Turkey/13/2006(H5N1) | 1 | | | TCID50 | Value given | Virulent | HIGH |
| | BALB/C | rA/H5TK13(1345678)/L969(2)(H5N1) | 2.3 | | | TCID50 | Value given | Virulent | HIGH |
| | BALB/C | rA/H5TK13(1234568)/L969(7)(H5N1) | 4.4 | | | TCID50 | Value given | Virulent | INTERMEDIATE |
| | BALB/C | rA/H5TK13(123458)/L969(67)(H5N1) | 4.6 | | | TCID50 | Value given | Virulent | INTERMEDIATE |
| | BALB/C | rA/H5TK13(1234567)/L969(8)(H5N1) | 0.3 | | | TCID50 | Value given | Virulent | HIGH |
| | BALB/C | rA/H5TK13(124567)/L969(38)(H5N1) | 1.8 | | | TCID50 | Value given | Virulent | HIGH |
| | BALB/C | rA/H5TK13(123457)/L969(68)(H5N1) | 3.7 | | | TCID50 | Value given | Virulent | INTERMEDIATE |
| Gabriel *et al*., 2005 [18] | BALB/C | A/seal/Massachussetts/1/1980(H7N7) | | 6 | | PFU | Value given | Avirulent | LOW |
| | BALB/C | A/seal/Massachussetts/1-SC35M/1980(H7N7) | 2.8 | | | PFU | Value given | Virulent | HIGH |
| | BALB/C | rA/SC35(2345678)/SC35M(1)(H7N7) | 5.6 | | | PFU | Value given | Virulent | INTERMEDIATE |
| | BALB/C | rA/SC35(1345678)/SC35M(2)(H7N7) | | 6 | | PFU | Value given | Avirulent | LOW |
| | BALB/C | rA/SC35(1245678)/SC35M(3)(H7N7) | 6 | | | PFU | Value given | Virulent | INTERMEDIATE |
| | BALB/C | rA/SC35(1235678)/SC35M(4)(H7N7) | | 6 | | PFU | Value given | Avirulent | LOW |
| | BALB/C | rA/SC35(1234678)/SC35M(5)(H7N7) | | 6 | | PFU | Value given | Avirulent | LOW |
| | BALB/C | rA/SC35(1234578)/SC35M(6)(H7N7) | | 6 | | PFU | Value given | Avirulent | LOW |
| | BALB/C | rA/SC35/PB1-13P(H7N7) | | 6 | | PFU | Value given | Avirulent | LOW |
| | BALB/C | rA/SC35/PB1-678N(H7N7) | | 6 | | PFU | Value given | Avirulent | LOW |
| | BALB/C | rA/SC35/PB2-701N(H7N7) | 6 | | | PFU | Value given | Virulent | INTERMEDIATE |
| | BALB/C | rA/SC35/PB2-714R(H7N7) | 6 | | | PFU | Value given | Virulent | INTERMEDIATE |
| | BALB/C | rA/SC35/PB2-701N-714R(H7N7) | 3.5 | | | PFU | Value given | Virulent | INTERMEDIATE |
| | BALB/C | rA/SC35/PB2-701N/SC35M(5)(H7N7) | 5.5 | | | PFU | Value given | Virulent | INTERMEDIATE |
| | BALB/C | rA/SC35/PB2-714R/SC35M(5)(H7N7) | 5.5 | | | PFU | Value given | Virulent | INTERMEDIATE |
| | BALB/C | rA/SC35/PB2-701N-714R/SC35M(5)(H7N7) | 6 | | | PFU | Value given | Virulent | INTERMEDIATE |
| | BALB/C | rA/SC35M/PB1-13L(H7N7) | 4.2 | | | PFU | Value given | Virulent | INTERMEDIATE |
| | BALB/C | rA/SC35M/PB1-678S(H7N7) | 3.5 | | | PFU | Value given | Virulent | INTERMEDIATE |
| | BALB/C | rA/SC35M/PB2-333T(H7N7) | 1.4 | | | PFU | Value given | Virulent | HIGH |
| | BALB/C | rA/SC35M/PB2-701D(H7N7) | 3.3 | | | PFU | Value given | Virulent | INTERMEDIATE |
| | BALB/C | rA/SC35M/PB2-714S(H7N7) | | 6 | | PFU | Value given | Avirulent | LOW |
| Gariglianv *et al*., 2010 [19] | FVB/NJ | A/swine/Iowa/4/1976(H1N1) | | 6 | | TCID50 | Value given | Avirulent | LOW |
| | FVB/NJ | A/crestedeagle/Belgium/1/2004(H5N1) | | 6 | | TCID50 | Value given | Avirulent | LOW |
| Hatesuer *et al*., 2013 [20] | C57BL/6 | A/PR8M/1934(H1N1) | | | 5.3 | FFU | LD50_LB or LD50_UB is based on survival rate | Virulent | INTERMEDIATE |
| | C57BL/6 | A/Hamburg/4/2009(H1N1) | | | 5.3 | FFU | LD50_LB or LD50_UB is based on weight loss | Virulent | INTERMEDIATE |
| | C57BL/6 | A/PR8F/1934(H1N1) | | | 3.3 | FFU | LD50_LB or LD50_UB is based on weight loss | Virulent | HIGH |

| | | | | | | | | | |
|---|---|---|---|---|---|---|---|---|---|
| | C57BL/6 | maA/HongKong/1/1968(H3N2) | | 1 | | FFU | LD50_LB or LD50_UB is based on weight loss | Virulent | HIGH |
| | C57BL/6 | A/seal/Massachussetts/1-SC35M/1980(H7N7) | | | 4.3 | FFU | LD50_LB or LD50_UB is based on survival rate | Virulent | HIGH |
| Hatta *et al*., 2001 [21] | BALB/C | A/HongKong/483/1997(H5N1) | 0.26 | | | PFU | Value given | Virulent | HIGH |
| | BALB/C | rA/HongKong/483/1997(H5N1) | 0.23 | | | PFU | Value given | Virulent | HIGH |
| | BALB/C | A/HongKong/486/1997(H5N1) | | 3.88 | | PFU | Value given | Avirulent | LOW |
| | BALB/C | rA/HK486/HA-227I(H5N1) | | 6.11 | | PFU | Value given | Avirulent | LOW |
| | BALB/C | rA/HongKong/486/1997(H5N1) | 4.66 | | | PFU | Value given | Virulent | INTERMEDIATE |
| | BALB/C | rA/HK483/HK486-PA(H5N1) | 0.23 | | | PFU | Value given | Virulent | HIGH |
| | BALB/C | rA/HK483/HK486-PB1(H5N1) | 0.3 | | | PFU | Value given | Virulent | HIGH |
| | BALB/C | rA/HK483/HK486-PB2(H5N1) | 4 | | | PFU | Value given | Virulent | INTERMEDIATE |
| | BALB/C | rA/HK483/HK486-HA227I(H5N1) | 2.04 | | | PFU | Value given | Virulent | HIGH |
| | BALB/C | rA/HK483/HK486-HA227S(H5N1) | | | 0 | PFU | Value given | Virulent | HIGH |
| | BALB/C | rA/HK483/HK486-NP(H5N1) | 0.15 | | | PFU | Value given | Virulent | HIGH |
| | BALB/C | rA/HK483/HK486-NA(H5N1) | 0.61 | | | PFU | Value given | Virulent | HIGH |
| | BALB/C | rA/HK483/HK486-M(H5N1) | | | 0 | PFU | Value given | Virulent | HIGH |
| | BALB/C | rA/HK483/HK486-NS(H5N1) | 0.58 | | | PFU | Value given | Virulent | HIGH |
| | BALB/C | rA/HK486-HA227I/HK483-PA(H5N1) | | 5.6 | | PFU | Value given | Avirulent | LOW |
| | BALB/C | rA/HK486-HA227S/HK483-PA(H5N1) | | 5.04 | | PFU | Value given | Avirulent | LOW |
| | BALB/C | rA/HK486-HA227I/HK483-PB1(H5N1) | | 5.18 | | PFU | Value given | Avirulent | LOW |
| | BALB/C | rA/HK486-HA227S/HK483-PB1(H5N1) | | 4.52 | | PFU | Value given | Avirulent | LOW |
| | BALB/C | rA/HK486-HA227I/HK483-PB2(H5N1) | 3.53 | | | PFU | Value given | Virulent | INTERMEDIATE |
| | BALB/C | rA/HK486-HA227S/HK483-PB2(H5N1) | | | 0 | PFU | Value given | Virulent | HIGH |
| | BALB/C | rA/HK486/HK483-HA(H5N1) | 2.64 | | | PFU | Value given | Virulent | HIGH |
| | BALB/C | rA/HK486-HA227I/HK483-NP(H5N1) | | 5.9 | | PFU | Value given | Avirulent | LOW |
| | BALB/C | rA/HK486-HA227S/HK483-NP(H5N1) | | 4.57 | | PFU | Value given | Avirulent | LOW |
| | BALB/C | rA/HK486-HA227I/HK483-NA(H5N1) | | 5.9 | | PFU | Value given | Avirulent | LOW |
| | BALB/C | rA/HK486-HA227S/HK483-NA(H5N1) | 2.28 | | | PFU | Value given | Virulent | HIGH |
| | BALB/C | rA/HK486-HA227I/HK483-M(H5N1) | | 5.36 | | PFU | Value given | Avirulent | LOW |
| | BALB/C | rA/HK486-HA227S/HK483-M(H5N1) | 2.73 | | | PFU | Value given | Virulent | HIGH |
| | BALB/C | rA/HK486-HA227I/HK483-NS(H5N1) | | 5.58 | | PFU | Value given | Avirulent | LOW |
| | BALB/C | rA/HK486-HA227S/HK483-NS(H5N1) | 3.81 | | | PFU | Value given | Virulent | INTERMEDIATE |
| | BALB/C | rA/HK483/HA-cleaved(H5N1) | | 5.91 | | PFU | Value given | Avirulent | LOW |
| | BALB/C | A/HongKong/486/1997(H5N1) | 4 | | | PFU | Value given | Virulent | INTERMEDIATE |
| | BALB/C | rA/HK483/PB2-675I(H5N1) | | | 0 | PFU | Value given | Virulent | HIGH |

| | | | | | | | | | |
|---|---|---|---|---|---|---|---|---|---|
| | BALB/C | rA/HK486/PB2-675L(H5N1) | 3.32 | | | PFU | Value given | Virulent | INTERMEDIATE |
| | BALB/C | rA/HK483/PB2-627E(H5N1) | 3.36 | | | PFU | Value given | Virulent | INTERMEDIATE |
| | BALB/C | rA/HK486/PB2-627K(H5N1) | 0.76 | | | PFU | Value given | Virulent | HIGH |
| Huo *et al.*, 2018 [22] | C57BL/6 | A/WSN/1933(H1N1) | | | 2.3 | PFU | LD50_LB or LD50_UB is based on survival rate | Virulent | HIGH |
| | C57BL/6 | A/chicken/Henan/01/2004(H5N1) | | | 2.3 | PFU | LD50_LB or LD50_UB is based on survival rate | Virulent | HIGH |
| Ilyushina *et al.*, 2010 [23] | BALB/C | A/California/04/2009(H1N1) | 5.1 | | | PFU | Value given | Virulent | INTERMEDIATE |
| | BALB/C | A/California/04/MA1/I/2009(H1N1) | 1.8 | | | PFU | Value given | Virulent | HIGH |
| | BALB/C | A/California/04/MA2/I/2009(H1N1) | 1.8 | | | PFU | Value given | Virulent | HIGH |
| | BALB/C | A/Tennessee/1-560/2009(H1N1) | 5 | | | PFU | Value given | Virulent | INTERMEDIATE |
| | BALB/C | A/Tennessee/1-560/MA1/I/2009(H1N1) | 2.6 | | | PFU | Value given | Virulent | HIGH |
| | BALB/C | A/Tennessee/1-560/MA2/I/2009(H1N1) | 0.6 | | | PFU | Value given | Virulent | HIGH |
| Imai *et al.*, 2017 [24] | BALB/C | A/Guangdong/Th008/2017(H7N9) | 2.7 | | | PFU | Value given | Virulent | HIGH |
| | BALB/C | A/Guangdong/Th008/NA294R/2017(H7N9) | 1.7 | | | PFU | Value given | Virulent | HIGH |
| | BALB/C | A/Guangdong/Th008/NA294K/2017(H7N9) | 4.8 | | | PFU | Value given | Virulent | INTERMEDIATE |
| | BALB/C | A/Anhui/1/2013(H7N9) | 4.5 | | | PFU | Value given | Virulent | INTERMEDIATE |
| Itoh *et al.*, 2009 [25] | BALB/C | A/California/04/2009(H1N1) | 5.8 | | | PFU | Value given | Virulent | INTERMEDIATE |
| | BALB/C | A/Kawasaki/UTK-4/2009(H1N1) | | 6.6 | | PFU | Value given | Avirulent | LOW |
| Jang *et al.*, 2013a [26] | BALB/C | rA/X-31(123578)/KOR09(46)(H1N1) | | 5 | | PFU | LD50_LB or LD50_UB is based on weight loss | Avirulent | LOW |
| | BALB/C | A/Seoul/Y-01/2009(H1N1) | 2.3 | | | PFU | Value given; In the text: " … the challenge dose of 103 PFU corresponds to 5 MLD50" | Virulent | HIGH |
| Jang *et al.*, 2013b [27] | BALB/C | rA/X-31(H3N2) | | 5 | | PFU | LD50_LB or LD50_UB is based on weight loss | Avirulent | LOW |
| | BALB/C | rA/X-31(123578)/INA5(46)(H5N1) | | 5 | | PFU | LD50_LB or LD50_UB is based on weight loss | Avirulent | LOW |
| | BALB/C | rA/X-31(123578)/KOR03(46)(H5N1) | | 5 | | PFU | LD50_LB or LD50_UB is based on weight loss | Avirulent | LOW |
| Jang *et al.*, 2014 [28] | BALB/C | A/NewCaledonia/20/1999(H1N1) | 6 | | | PFU | LD50_LB or LD50_UB is based on survival rate | Virulent | INTERMEDIATE |
| | BALB/C | rA/X-31(123578)/NC99(46)(H1N1) | | 6 | | PFU | LD50_LB or LD50_UB is based on survival rate | Avirulent | LOW |
| | BALB/C | A/Panama/2007/1999(H3N2) | | 6 | | PFU | LD50_LB or LD50_UB is based on survival rate | Avirulent | LOW |
| | BALB/C | rA/X-31(123578)/PAN99(46)(H3N2) | | 6 | | PFU | LD50_LB or LD50_UB is based on survival rate | Avirulent | LOW |
| Jang *et al.*, 2018 [29] | BALB/C | A/PuertoRico/8/1934(H1N1) | 3.7 | | | PFU | Value given | Virulent | INTERMEDIATE |
| | BALB/C | mA/aquaticbird/Korea/w81/2005(H5N2) | 4 | | | PFU | Value given | Virulent | INTERMEDIATE |
| | BALB/C | A/Philippines/2/1982(H3N2) | 4.7 | | | PFU | Value given | Virulent | INTERMEDIATE |
| | BALB/C | rA/PR8(1235678)/NL03(4)(H7N1) | 3.7 | | | PFU | Value given | Virulent | INTERMEDIATE |
| Jiao *et al.*, 2008 [30] | BALB/C | A/duck/Guangxi/12/2003(H5N1) | 6.4 | | | EID50 | Value given | Avirulent | LOW |
| | BALB/C | A/duck/Guangxi/27/2003(H5N1) | 0.6 | | | EID50 | Value given | Virulent | HIGH |
| | BALB/C | rA/duck/Guangxi/12/2003(H5N1) | 6.4 | | | EID50 | Value given | Avirulent | LOW |

| | | | | | | | | | |
|---|---|---|---|---|---|---|---|---|---|
| | BALB/C | rA/duck/Guangxi/27/2003(H5N1) | 0.6 | | | EID50 | Value given | Virulent | HIGH |
| | BALB/C | rA/DKGX12(2345678)/DKGX27(1)(H5N1) | 6.4 | | | EID50 | Value given | Avirulent | LOW |
| | BALB/C | rA/DKGX12(1245678)/DKGX27(3)(H5N1) | 6.4 | | | EID50 | Value given | Avirulent | LOW |
| | BALB/C | rA/DKGX12(1235678)/DKGX27(4)(H5N1) | 6.4 | | | EID50 | Value given | Avirulent | LOW |
| | BALB/C | rA/DKGX12(1234678)/DKGX27(5)(H5N1) | 6.4 | | | EID50 | Value given | Avirulent | LOW |
| | BALB/C | rA/DKGX12(1234567)/DKGX27(8)(H5N1) | 2 | | | EID50 | Value given | Virulent | HIGH |
| | BALB/C | rA/DKGX27(2345678)/DKGX12(1)(H5N1) | 0.6 | | | EID50 | Value given | Virulent | HIGH |
| | BALB/C | rA/DKGX27(1245678)/DKGX12(3)(H5N1) | 1.5 | | | EID50 | Value given | Virulent | HIGH |
| | BALB/C | rA/DKGX27(1235678)/DKGX12(4)(H5N1) | 1.5 | | | EID50 | Value given | Virulent | HIGH |
| | BALB/C | rA/DKGX27(1234678)/DKGX12(5)(H5N1) | 0.6 | | | EID50 | Value given | Virulent | HIGH |
| | BALB/C | rA/DKGX27(1234567)/DKGX12(8)(H5N1) | 6.4 | | | EID50 | Value given | Avirulent | LOW |
| | BALB/C | rA/DKGX12/NS1-P42S(H5N1) | 2.2 | | | EID50 | Value given | Virulent | HIGH |
| | BALB/C | rA/DKGX12/NS1-N48S(H5N1) | 6.4 | | | EID50 | Value given | Avirulent | LOW |
| | BALB/C | rA/DKGX27/NS1-S42P(H5N1) | 6.4 | | | EID50 | Value given | Avirulent | LOW |
| | BALB/C | rA/DKGX27/NS1-S48N(H5N1) | 0.8 | | | EID50 | Value given | Virulent | HIGH |
| | BALB/C | rA/DKGX27/NS1-R38A(H5N1) | 0.8 | | | EID50 | Value given | Virulent | HIGH |
| | BALB/C | rA/DKGX27/NS1-K41A(H5N1) | 0.6 | | | EID50 | Value given | Virulent | HIGH |
| | BALB/C | rA/DKGX27/NS1-R38A-K41A(H5N1) | 6.4 | | | EID50 | Value given | Avirulent | LOW |
| Joseph *et al*., 2007 [31] | BALB/C | A/Netherlands/219/2003(H7N7) | 0.8 | | | TCID50 | Value given | Virulent | HIGH |
| | BALB/C | A/turkey/England/1963(H7N3) | 0.5 | | | TCID50 | Value given | Virulent | HIGH |
| | BALB/C | A/turkey/VA/55/2002(H7N2) | | 5 | | TCID50 | LD50_LB or LD50_UB is based on survival rate | Avirulent | LOW |
| | BALB/C | A/turkey/Utah/24721-10/1995(H7N3) | | 5 | | TCID50 | LD50_LB or LD50_UB is based on survival rate | Virulent | INTERMEDIATE |
| Katz *et al*., 2000 [32] | BALB/C | A/HongKong/481/1997(H5N1) | 1.7 | | | EID50 | Value given | Virulent | HIGH |
| | BALB/C | A/HongKong/483/1997(H5N1) | 2.4 | | | EID50 | Value given | Virulent | HIGH |
| | BALB/C | A/HongKong/485/1997(H5N1) | 2.9 | | | EID50 | Value given | Virulent | HIGH |
| | BALB/C | A/HongKong/491/1997(H5N1) | 2 | | | EID50 | Value given | Virulent | HIGH |
| | BALB/C | A/HongKong/503/1997(H5N1) | 2 | | | EID50 | Value given | Virulent | HIGH |
| | BALB/C | A/HongKong/514/1997(H5N1) | | | 1.5 | EID50 | Value given | Virulent | HIGH |
| | BALB/C | A/HongKong/516/1997(H5N1) | | | 1.5 | EID50 | Value given | Virulent | HIGH |
| | BALB/C | A/HongKong/532/1997(H5N1) | | | 1.5 | EID50 | Value given | Virulent | HIGH |
| | BALB/C | A/HongKong/542/1997(H5N1) | | | 1.5 | EID50 | Value given | Virulent | HIGH |
| | BALB/C | A/HongKong/156/1997(H5N1) | 5.9 | | | EID50 | Value given | Virulent | INTERMEDIATE |
| | BALB/C | A/HongKong/486/1997(H5N1) | | 6.5 | | EID50 | Value given | Avirulent | LOW |
| | BALB/C | A/HongKong/488/1997(H5N1) | | 6.5 | | EID50 | Value given | Avirulent | LOW |

| | | | | | | | | | |
|---|---|---|---|---|---|---|---|---|---|
| | BALB/C | A/HongKong/507/1997(H5N1) | | 6.5 | | EID50 | Value given | Avirulent | LOW |
| | BALB/C | A/HongKong/538/1997(H5N1) | | 6.5 | | EID50 | Value given | Avirulent | LOW |
| | BALB/C | A/HongKong/97/1998(H5N1) | | 6.5 | | EID50 | Value given | Avirulent | LOW |
| Kim *et al*., 2013 [33] | BALB/C | A/Korea/01/2009(H1N1) | | 6 | | PFU | Value given | Avirulent | LOW |
| | DBA/2 | A/Korea/01/2009(H1N1) | 2.83 | | | PFU | Value given | Virulent | HIGH |
| Kobasa *et al*., 2004 [34] | BALB/C | rA/WSN33(123578)/SC18(46)(H1N1) | 3 | | | PFU | Value given | Virulent | HIGH |
| | BALB/C | A/WSN/1933(H1N1) | 3.3 | | | PFU | Value given | Virulent | INTERMEDIATE |
| | BALB/C | rA/M88(123578)/SC18(46)(H1N1) | 5.2 | | | PFU | Value given | Virulent | INTERMEDIATE |
| | BALB/C | rA/M88(123578)/SC18(4)/K173(6)(H1N1) | 4.4 | | | PFU | Value given | Virulent | INTERMEDIATE |
| | BALB/C | A/Memphis/8/1988(H3N2) | | 6.2 | | PFU | Value given | Avirulent | LOW |
| | BALB/C | rA/M88(123578)/WSN33(46)(H1N1) | | 6.7 | | PFU | Value given | Avirulent | LOW |
| | BALB/C | rA/K173(123578)/SC18(46)(H1N1) | 6.9 | | | PFU | Value given | Avirulent | LOW |
| | BALB/C | rA/K173(1235678)/SC18(4)(H1N1) | 5.2 | | | PFU | Value given | Virulent | INTERMEDIATE |
| | BALB/C | A/Kawasaki/173/2001(H1N1) | | 7.4 | | PFU | Value given | Avirulent | LOW |
| | BALB/C | rA/K173(123578)/WSN33(46)(H1N1) | | 6.9 | | PFU | Value given | Avirulent | LOW |
| Kwon *et al*., 2018 [35] | BALB/C | A/environment/Korea/W541/2016(H5N6) | 5.5 | | | TCID50 | Value given | Virulent | INTERMEDIATE |
| | BALB/C | A/commonteal/Korea/W555/2017(H5N8) | 5.2 | | | TCID50 | Value given | Virulent | INTERMEDIATE |
| Lee *et al*., 2018 [36] | BALB/C | A/broilerduck/Korea/Buan2/2014(H5N8) | 2.7 | | | TCID50 | Value given | Virulent | HIGH |
| | BALB/C | rA/broilerduck/Korea/Buan2/2014(H5N8) | 3.4 | | | TCID50 | Value given | Virulent | INTERMEDIATE |
| | BALB/C | rA/broilerduck/Korea/Buan2-QRET/2014(H5N8) | | 4.7 | | TCID50 | Value given | Avirulent | LOW |
| | BALB/C | rA/broilerduck/Korea/Buan2-LRET/2014(H5N8) | | 4.7 | | TCID50 | Value given | Avirulent | LOW |
| Leist *et al*., 2016 [37] | C57BL/6 | maA/HongKong/1/1968(H3N2) | | | 1 | FFU | LD50_LB or LD50_UB is based on survival rate | Virulent | HIGH |
| | A/J | maA/HongKong/1/1968(H3N2) | | | 1 | FFU | LD50_LB or LD50_UB is based on survival rate | Virulent | HIGH |
| | 129S1/SvImJ | maA/HongKong/1/1968(H3N2) | | | 1 | FFU | LD50_LB or LD50_UB is based on survival rate | Virulent | HIGH |
| | NOD/ShiLtJ | maA/HongKong/1/1968(H3N2) | 1.38 | | | FFU | Using formula Reed and Muench method | Virulent | HIGH |
| | NZO/HILtJ | maA/HongKong/1/1968(H3N2) | | 5 | | FFU | LD50_LB or LD50_UB is based on survival rate | Avirulent | LOW |
| | CAST/EiJ | maA/HongKong/1/1968(H3N2) | | | 1 | FFU | LD50_LB or LD50_UB is based on survival rate | Virulent | HIGH |
| | PWK/PhJ | maA/HongKong/1/1968(H3N2) | | 5 | | FFU | LD50_LB or LD50_UB is based on survival rate | Avirulent | LOW |
| | WSB/EiJ | maA/HongKong/1/1968(H3N2) | | | 1 | FFU | LD50_LB or LD50_UB is based on survival rate | Virulent | HIGH |
| Li *et al*., 2005 [38] | BALB/C | A/duck/Guangxi/22/2001(H5N1) | | 6.5 | | EID50 | Value given; Karber Spearman method for LD50 | Avirulent | LOW |
| | BALB/C | A/duck/Guangxi/35/2001(H5N1) | 1.8 | | | EID50 | Value given; Karber Spearman method for LD50 | Virulent | HIGH |
| | BALB/C | rA/duck/Guangxi/22/2001(H5N1) | | 6.5 | | EID50 | Value given; Karber Spearman method for LD50 | Avirulent | LOW |

| | | | | | | | | | |
|---|---|---|---|---|---|---|---|---|---|
| | BALB/C | rA/duck/Guangxi/35/2001(H5N1) | 2.3 | | | EID50 | Value given; Karber Spearman method for LD50 | Virulent | HIGH |
| | BALB/C | rA/DKGX22(2345678)/DKGX35(1)(H5N1) | 4.3 | | | EID50 | Value given; Karber Spearman method for LD50 | Virulent | INTERMEDIATE |
| | BALB/C | rA/DKGX22(1345678)/DKGX35(2)(H5N1) | | 6.5 | | EID50 | Value given; Karber Spearman method for LD50 | Avirulent | LOW |
| | BALB/C | rA/DKGX22(1245678)/DKGX35(3)(H5N1) | | 6.5 | | EID50 | Value given; Karber Spearman method for LD50 | Avirulent | LOW |
| | BALB/C | rA/DKGX22(1235678)/DKGX35(4)(H5N1) | | 6.5 | | EID50 | Value given; Karber Spearman method for LD50 | Avirulent | LOW |
| | BALB/C | rA/DKGX22(1234678)/DKGX35(5)(H5N1) | | 6.5 | | EID50 | Value given; Karber Spearman method for LD50 | Avirulent | LOW |
| | BALB/C | rA/DKGX22(1234578)/DKGX35(6)(H5N1) | | 6.5 | | EID50 | Value given; Karber Spearman method for LD50 | Avirulent | LOW |
| | BALB/C | rA/DKGX22(1234568)/DKGX35(7)(H5N1) | | 6.5 | | EID50 | Value given; Karber Spearman method for LD50 | Avirulent | LOW |
| | BALB/C | rA/DKGX22(1234567)/DKGX35(8)(H5N1) | | 6.5 | | EID50 | Value given; Karber Spearman method for LD50 | Avirulent | LOW |
| | BALB/C | rA/DKGX35(2345678)/DKGX22(1)(H5N1) | 6.3 | | | EID50 | Value given; Karber Spearman method for LD50 | Avirulent | LOW |
| | BALB/C | rA/DKGX35(1345678)/DKGX22(2)(H5N1) | 1.7 | | | EID50 | Value given; Karber Spearman method for LD50 | Virulent | HIGH |
| | BALB/C | rA/DKGX35(1245678)/DKGX22(3)(H5N1) | 5.2 | | | EID50 | Value given; Karber Spearman method for LD50 | Virulent | INTERMEDIATE |
| | BALB/C | rA/DKGX35(1235678)/DKGX22(4)(H5N1) | 2.8 | | | EID50 | Value given; Karber Spearman method for LD50 | Virulent | HIGH |
| | BALB/C | rA/DKGX35(1234678)/DKGX22(5)(H5N1) | 3.2 | | | EID50 | Value given; Karber Spearman method for LD50 | Virulent | INTERMEDIATE |
| | BALB/C | rA/DKGX35(1234578)/DKGX22(6)(H5N1) | 3.7 | | | EID50 | Value given; Karber Spearman method for LD50 | Virulent | INTERMEDIATE |
| | BALB/C | rA/DKGX35(1234568)/DKGX22(7)(H5N1) | 2.8 | | | EID50 | Value given; Karber Spearman method for LD50 | Virulent | HIGH |
| | BALB/C | rA/DKGX35(1234567)/DKGX22(8)(H5N1) | 4.5 | | | EID50 | Value given; Karber Spearman method for LD50 | Virulent | INTERMEDIATE |
| Liedmann *et al*., 2014 [39] | C57BL/6 | A/PuertoRico/8/1934(H1N1) | | | 3 | PFU | LD50_LB or LD50_UB is based on survival rate | Virulent | HIGH |
| | C57BL/6 | A/PR8/Mutant(H1N1) | | 3 | | PFU | LD50_LB or LD50_UB is based on survival rate | Virulent | INTERMEDIATE |
| | BALB/C | A/PuertoRico/8/1934(H1N1) | | | 2.5 | PFU | LD50_LB or LD50_UB is based on survival rate | Virulent | HIGH |
| | BALB/C | rA/PR8(2345678)/PR8M(1)(H1N1) | | 2.5 | | PFU | LD50_LB or LD50_UB is based on survival rate | Virulent | INTERMEDIATE |
| | BALB/C | rA/PR8(1234678)/PR8M(5)(H1N1) | | 2.5 | | PFU | LD50_LB or LD50_UB is based on survival rate | Virulent | INTERMEDIATE |
| | BALB/C | rA/PR8(1234578)/PR8M(6)(H1N1) | | | 2.5 | PFU | LD50_LB or LD50_UB is based on survival rate | Virulent | HIGH |
| | BALB/C | rA/PR8(1234567)/PR8M(8)(H1N1) | | | 2.5 | PFU | LD50_LB or LD50_UB is based on survival rate | Virulent | HIGH |
| | DBA/2 | rA/PR8(4678)/PR8M(1235)(H1N1) | | 1 | | PFU | LD50_LB or LD50_UB is based on survival rate | Virulent | HIGH |
| | DBA/2 | A/PuertoRico/8/1934(H1N1) | | | 1 | PFU | LD50_LB or LD50_UB is based on survival rate | Virulent | HIGH |
| Long *et al*., 2008 [40] | BALB/C | A/duck/Shandong/093/2004(H5N1) | 6.2 | | | EID50 | Value given | Avirulent | LOW |
| | BALB/C | rA/WSN33(12357)/SD093(468)(H5N1) | 3.17 | | | EID50 | Value given | Virulent | INTERMEDIATE |
| | BALB/C | rA/WSN33(12357)/SD093(468)/NSdel(H5N1) | 2.5 | | | EID50 | Value given | Virulent | HIGH |
| | BALB/C | rA/WSN33(12357)/SD093(46)/YZ232(8)(H5N1) | 3.1 | | | EID50 | Value given | Virulent | INTERMEDIATE |
| | BALB/C | rA/WSN33(12357)/SD093(46)/YZ232(8)/NSins(H5N1) | 4.5 | | | EID50 | Value given | Virulent | INTERMEDIATE |
| | BALB/C | rA/WSN33(12357)/SD093(468)/NS1-E97D(H5N1) | 4.27 | | | EID50 | Value given | Virulent | INTERMEDIATE |
| | BALB/C | rA/WSN33(12357)/SD093(468)/NSdel/NS1-E92D(H5N1) | 4.58 | | | EID50 | Value given | Virulent | INTERMEDIATE |
| Lu et al., 1999 [41] | BALB/C | A/HongKong/483/1997(H5N1) | 2.4 | | | EID50 | Value given | Virulent | HIGH |

| | | | | | | | | | |
|---|---|---|---|---|---|---|---|---|---|
| | BALB/C | A/HongKong/485/1997(H5N1) | 2.9 | | | EID50 | Value given | Virulent | HIGH |
| | BALB/C | A/HongKong/156/1997(H5N1) | 5.9 | | | EID50 | Value given | Virulent | INTERMEDIATE |
| | BALB/C | A/HongKong/486/1997(H5N1) | | 7 | | EID50 | Value given | Avirulent | LOW |
| | BALB/C | rA/X-31(H3N2) | | 5.2 | | EID50 | Value given | Avirulent | LOW |
| Lu *et al*., 2018 [42] | BALB/C | A/NorthernShoveler/Ningxia/488-53/2015(H5N6) | | 6 | | EID50 | LD50_LB or LD50_UB is based on survival rate | Avirulent | LOW |
| Maines *et al*., 2005 [43] | BALB/C | A/HongKong/483/1997(H5N1) | 1.6 | | | EID50 | Value given | Virulent | HIGH |
| | BALB/C | A/Thailand/16/2004(H5N1) | 1.7 | | | EID50 | Value given | Virulent | HIGH |
| | BALB/C | A/VietNam/1203/2004(H5N1) | 2.2 | | | EID50 | Value given | Virulent | HIGH |
| | BALB/C | A/VietNam/1204/2004(H5N1) | 3.8 | | | EID50 | Value given | Virulent | INTERMEDIATE |
| | BALB/C | A/Thailand/SP/83/2004(H5N1) | 5.5 | | | EID50 | Value given | Virulent | INTERMEDIATE |
| | BALB/C | A/chicken/Korea/es/2003(H5N1) | | 7 | | EID50 | Value given | Avirulent | LOW |
| | BALB/C | A/chicken/Indonesia/7/2003(H5N1) | | 7 | | EID50 | Value given | Avirulent | LOW |
| | BALB/C | A/chicken/VietNam/8/2003(H5N1) | 5.8 | | | EID50 | Value given | Virulent | INTERMEDIATE |
| Manicassamy *et al*., 2010 [44] | C57BL/6 | A/California/04/2009(H1N1) | 4.7 | | | PFU | LD50_LB or LD50_UB is based on survival rate | Virulent | INTERMEDIATE |
| | C57BL/6 | A/Netherlands/602/2009(H1N1) | 4.2 | | | PFU | Value given | Virulent | INTERMEDIATE |
| | BALB/C | A/California/04/2009(H1N1) | 4.7 | | | PFU | LD50_LB or LD50_UB is based on survival rate | Virulent | INTERMEDIATE |
| Mase *et al*., 2005 [45] | BALB/C | A/duck/Yokohama/aq10/2003(H5N1) | 6.7 | | | EID50 | Value given | Avirulent | LOW |
| | BALB/C | A/duck/Yokohama/aq10/VAR1/2003(H5N1) | 1.97 | | | EID50 | Value given | Virulent | HIGH |
| | BALB/C | A/duck/Yokohama/aq10/VAR2/2003(H5N1) | 2 | | | EID50 | Value given | Virulent | HIGH |
| Metreveli *et al*., 2014 [46] | C57BL/6 | rA/swine/Sweden/1021/2009(H1N2) | 4.9 | | | PFU | Value given; It is written as MID50 in the paper | Virulent | INTERMEDIATE |
| | C57BL/6 | rA/swine/Sweden/9706/2010(H1N2) | 6.1 | | | PFU | Value given; It is written as MID50 in the paper | Avirulent | LOW |
| | C57BL/6 | rA/SWE1021(1345678)/SWE9706(2)(H1N2) | | 6 | | PFU | LD50_LB or LD50_UB is based on weight loss | Avirulent | LOW |
| | C57BL/6 | rA/SWE1021(2)/SWE9706(1345678)(H1N2) | | 6 | | PFU | LD50_LB or LD50_UB is based on weight loss | Avirulent | LOW |
| Mifsud *et al*., 2015 [47] | C57BL/6 | A/Memphis/1/1971(H3N2) | | 4.5 | | PFU | LD50_LB or LD50_UB is based on weight loss | Avirulent | LOW |
| Na *et al*., 2016 [48] | C57BL/6 | rA/PuertoRico/8/1934(H1N1) | 3.9 | | | EID50 | Value given | Virulent | INTERMEDIATE |
| | C57BL/6 | rA/PR8(1234567)/MI63(8)(H1N1) | 4.3 | | | EID50 | Value given | Virulent | INTERMEDIATE |
| | C57BL/6 | rA/PR8(1234567)/KYG11(8)(H1N1) | 7.8 | | | EID50 | Value given | Avirulent | LOW |
| Nurnberger *et al*., 2016 [49] | C57BL/6 | A/swan/Germany/R65/2006(H5N1) | | | 2 | PFU | LD50 less than 10E1.0 based on Bogs et al., 2011 | Virulent | HIGH |
| | C57BL/6 | A/seal/Massachussetts/1-SC35M/1980(H7N7) | | | 3 | PFU | LD50_LB or LD50_UB is based on survival rate | Virulent | HIGH |
| O'Neill *et al*., 2000 [50] | C57BL/6 | A/HongKong/156/1997(H5N1) | | | 2.8 | EID50 | LD50_LB or LD50_UB is based on weight loss | Virulent | HIGH |
| | BALB/C | A/HongKong/156/1997(H5N1) | | | 4.2 | EID50 | LD50_LB or LD50_UB is based on weight loss | Virulent | HIGH |
| | BALB/C | A/goose/HongKong/437-6/1999(H5N1) | | 4.2 | | EID50 | LD50_LB or LD50_UB is based on weight loss | Avirulent | LOW |
| | BALB/C | A/HongKong/1073/1999(H9N2) | | 4.2 | | EID50 | LD50_LB or LD50_UB is based on weight loss | Virulent | INTERMEDIATE |
| | BALB/C | rA/goose/HongKong/437-6/1999(H5N1) | | 4.2 | | EID50 | LD50_LB or LD50_UB is based on weight loss | Avirulent | LOW |

| | | | | | | | | | |
|---|---|---|---|---|---|---|---|---|---|
| | BALB/C | rA/HK1073(15)/GSHK437(234678)(H5N1) | | | 4.2 | EID50 | LD50_LB or LD50_UB is based on weight loss | Virulent | HIGH |
| | BALB/C | rA/HongKong/1073/1999(H9N2) | | 4.2 | | EID50 | LD50_LB or LD50_UB is based on weight loss | Virulent | INTERMEDIATE |
| | BALB/C | A/quail/HongKong/G1/1997(H9N2) | | | | | MILD: "QHKG1 typically results in loss of up to 15% of the initial weight, but the mice recover and become healthy again." | Virulent | INTERMEDIATE |
| Otte *et al.*, 2011 [51] | BALB/C | A/SolomonIslands/3/2006(H1N1) | | 6 | | PFU | Value given | Avirulent | LOW |
| | BALB/C | A/Hamburg/05/2009(H1N1) | | 6 | | PFU | Value given | Avirulent | LOW |
| | BALB/C | A/Hamburg/NY1580/2009(H1N1) | | 6 | | PFU | Value given | Avirulent | LOW |
| | BALB/C | A/Thailand/KAN-1/2004(H5N1) | 0.3 | | | PFU | Value given | Virulent | HIGH |
| | C57BL/6 | A/SolomonIslands/3/2006(H1N1) | | 6 | | PFU | Value given | Avirulent | LOW |
| | C57BL/6 | A/Hamburg/05/2009(H1N1) | 5.2 | | | PFU | Value given | Virulent | INTERMEDIATE |
| | C57BL/6 | A/Hamburg/NY1580/2009(H1N1) | 3.5 | | | PFU | Value given | Virulent | INTERMEDIATE |
| | C57BL/6 | A/Thailand/KAN-1/2004(H5N1) | 1.8 | | | PFU | Value given | Virulent | HIGH |
| Otte *et al.*, 2015 [52] | C57BL/6 | rA/Hamburg/05/2009(H1N1) | 5.2 | | | PFU | Value given | Virulent | INTERMEDIATE |
| | C57BL/6 | A/Hamburg/05/2009(H1N1) | 5.2 | | | PFU | Value given | Virulent | INTERMEDIATE |
| | C57BL/6 | rA/Hamburg/NY1580/2009(H1N1) | 3.2 | | | PFU | Value given | Virulent | INTERMEDIATE |
| | C57BL/6 | A/Hamburg/NY1580/2009(H1N1) | 3.5 | | | PFU | Value given | Virulent | INTERMEDIATE |
| | C57BL/6 | rA/HH05(2345678)/HH15(1)(H1N1) | | 5 | | PFU | Value given; "… the PB2, NA, and NS genes of HH15 did not significantly affect pathogenicity in mice." | Virulent | INTERMEDIATE |
| | C57BL/6 | rA/HH05(1345678)/HH15(3)(H1N1) | 4.5 | | | PFU | Value given | Virulent | INTERMEDIATE |
| | C57BL/6 | rA/HH05(1234678)/HH15(5)(H1N1) | 3.2 | | | PFU | Value given | Virulent | INTERMEDIATE |
| | C57BL/6 | rA/HH05(1235678)/HH15(4)(H1N1) | 3.6 | | | PFU | Value given | Virulent | INTERMEDIATE |
| | C57BL/6 | rA/HH05(1234578)/HH15(6)(H1N1) | 4.4 | | | PFU | Value given; "… the PB2, NA, and NS genes of HH15 did not significantly affect pathogenicity in mice." | Virulent | INTERMEDIATE |
| | C57BL/6 | rA/HH05(1234567)/HH15(8)(H1N1) | | 5 | | PFU | Value given; "… the PB2, NA, and NS genes of HH15 did not significantly affect pathogenicity in mice." | Virulent | INTERMEDIATE |
| | C57BL/6 | rA/HH05(24678)/HH15(135)(H1N1) | 3.5 | | | PFU | Value given | Virulent | INTERMEDIATE |
| | C57BL/6 | rA/HH05(123578)/HH15(46)(H1N1) | 3.8 | | | PFU | Value given | Virulent | INTERMEDIATE |
| | C57BL/6 | A/Hamburg/05/NP100I/2009(H1N1) | 3.4 | | | PFU | Value given | Virulent | INTERMEDIATE |
| | C57BL/6 | A/Hamburg/05/NP133L/2009(H1N1) | 2.9 | | | PFU | Value given | Virulent | HIGH |
| | C57BL/6 | A/Hamburg/05/NP373T/2009(H1N1) | 3.9 | | | PFU | Value given | Virulent | INTERMEDIATE |

| | | | | | | | | | |
|---|---|---|---|---|---|---|---|---|---|
| Pan *et al*., 2018 [53] | BALB/C | A/Guangzhou/39715/2014(H5N6) | 0.7 | | | PFU | Value given; Inferred from "… intranasally infected with 2 MLD50 (10 pfu) of H5N6/GZ14 …" | Virulent | HIGH |
| Pica *et al*., 2011 [54] | C57BL/6 | A/HongKong/1/1968(H3N2) | | 6 | | PFU | Value given | Avirulent | LOW |
| | C57BL/6 | A/Netherlands/602/2009(H1N1) | 4.3 | | | PFU | Value given | Virulent | INTERMEDIATE |
| | C57BL/6 | A/Brisbane/10/2007(H3N2) | | 6 | | PFU | Value given | Avirulent | LOW |
| | C57BL/6 | A/Wisconsin/67/2005(H3N2) | | 6 | | PFU | Value given | Avirulent | LOW |
| | C57BL/6 | A/Panama/2007/1999(H3N2) | | 6 | | PFU | Value given | Avirulent | LOW |
| | C57BL/6 | A/Brisbane/59/2007(H1N1) | | 6 | | PFU | Value given | Avirulent | LOW |
| | C57BL/6 | A/NewCaledonia/20/1999(H1N1) | 5.9 | | | PFU | Value given; Average LD50 from 10E5.5 and 10E6.3 | Virulent | INTERMEDIATE |
| | C57BL/6 | A/SolomonIslands/3/2006(H1N1) | | 5.3 | | PFU | Value given; The stock titer of the A/Solomon Islands/03/06 virus did not allow inoculation at doses greater that 10E5.3 PFU. | Avirulent | LOW |
| | C57BL/6 | A/swine/Spain/53207/2004(H1N1) | 5.7 | | | PFU | Value given | Virulent | INTERMEDIATE |
| | C57BL/6 | A/swine/Kansas/77778/2007(H1N1) | 2.5 | | | PFU | Value given | Virulent | HIGH |
| | C57BL/6 | A/swine/Spain/40564/2002(H1N2) | 4.7 | | | PFU | Value given | Virulent | INTERMEDIATE |
| | C57BL/6 | A/swine/Spain/54008/2004(H3N2) | | 6 | | PFU | Value given | Avirulent | LOW |
| | C57BL/6 | A/swine/Texas/4199-2/1998(H3N2) | 6.2 | | | PFU | Value given | Avirulent | LOW |
| | C57BL/6 | A/duck/Alberta/35/1976(H1N1) | | 6 | | PFU | Value given | Avirulent | LOW |
| | C57BL/6 | A/duck/Ukraine/1/1963(H3N8) | | 6 | | PFU | Value given | Avirulent | LOW |
| | C57BL/6 | rA/PR8(123578)/ALB76(46)(H1N1) | 3.5 | | | PFU | Value given | Virulent | INTERMEDIATE |
| | C57BL/6 | rA/PR8(123578)/UKR63(46)(H3N8) | | 6 | | PFU | Value given | Avirulent | LOW |
| | C57BL/6 | A/PuertoRico/8/1934(H1N1) | 1.4 | | | PFU | Value given; Average LD50 from 10E1.3 and 10E1.5 | Virulent | HIGH |
| | C57BL/6 | rA/X-31(H3N2) | 5.3 | | | PFU | Value given | Virulent | INTERMEDIATE |
| | DBA/2 | A/HongKong/1/1968(H3N2) | 5.5 | | | PFU | Value given | Virulent | INTERMEDIATE |
| | DBA/2 | A/Netherlands/602/2009(H1N1) | 0.5 | | | PFU | Value given | Virulent | HIGH |
| | DBA/2 | A/Brisbane/10/2007(H3N2) | | 6 | | PFU | Value given | Avirulent | LOW |
| | DBA/2 | A/Wisconsin/67/2005(H3N2) | | 6 | | PFU | Value given | Avirulent | LOW |
| | DBA/2 | A/Panama/2007/1999(H3N2) | | 6 | | PFU | Value given | Avirulent | LOW |
| | DBA/2 | A/Brisbane/59/2007(H1N1) | 5 | | | PFU | Value given | Virulent | INTERMEDIATE |
| | DBA/2 | A/NewCaledonia/20/1999(H1N1) | 4.3 | | | PFU | Value given; Average LD50 from 10E3.8 and 10E4.7 | Virulent | INTERMEDIATE |
| | DBA/2 | A/SolomonIslands/3/2006(H1N1) | 3.9 | | | PFU | Value given | Virulent | INTERMEDIATE |
| | DBA/2 | A/swine/Spain/53207/2004(H1N1) | 2.2 | | | PFU | Value given | Virulent | HIGH |
| | DBA/2 | A/swine/Kansas/77778/2007(H1N1) | 1 | | | PFU | Value given | Virulent | HIGH |
| | DBA/2 | A/swine/Spain/40564/2002(H1N2) | 1.3 | | | PFU | Value given | Virulent | HIGH |

| | | | | | | | | | |
|---|---|---|---|---|---|---|---|---|---|
| | DBA/2 | A/swine/Spain/54008/2004(H3N2) | 5.5 | | | PFU | Value given | Virulent | INTERMEDIATE |
| | DBA/2 | A/swine/Texas/4199-2/1998(H3N2) | 5.7 | | | PFU | Value given | Virulent | INTERMEDIATE |
| | DBA/2 | A/duck/Alberta/35/1976(H1N1) | | 6 | | PFU | Value given | Avirulent | LOW |
| | DBA/2 | A/duck/Ukraine/1/1963(H3N8) | | 6 | | PFU | Value given | Avirulent | LOW |
| | DBA/2 | rA/PR8(123578)/ALB76(46)(H1N1) | 1.2 | | | PFU | Value given | Virulent | HIGH |
| | DBA/2 | rA/PR8(123578)/UKR63(46)(H3N8) | 5.5 | | | PFU | Value given | Virulent | INTERMEDIATE |
| | DBA/2 | A/PuertoRico/8/1934(H1N1) | 0.4 | | | PFU | Value given; Average LD50 from 10E0.3 and 10E0.5 | Virulent | HIGH |
| | DBA/2 | rA/X-31(H3N2) | 0.7 | | | PFU | Value given | Virulent | HIGH |
| Ping *et al.*, 2011 [55] | CD-1 | A/HongKong/1/1968(H3N2) | | 7.7 | | PFU | Value given | Avirulent | LOW |
| | CD-1 | A/HongKong/1-1/1968(H3N2) | | 7.7 | | PFU | Value given | Avirulent | LOW |
| | CD-1 | A/HongKong/1-2/1968(H3N2) | | 7.7 | | PFU | Value given | Avirulent | LOW |
| | CD-1 | A/HongKong/1-4/1968(H3N2) | | 7.7 | | PFU | Value given | Avirulent | LOW |
| | CD-1 | A/HongKong/1-5/1968(H3N2) | | 7.7 | | PFU | Value given | Avirulent | LOW |
| | CD-1 | A/HongKong/1-6/1968(H3N2) | | 7.7 | | PFU | Value given | Avirulent | LOW |
| | CD-1 | A/HongKong/1-8/1968(H3N2) | | 7.7 | | PFU | Value given | Avirulent | LOW |
| | CD-1 | A/HongKong/1-9/1968(H3N2) | | 7.7 | | PFU | Value given | Avirulent | LOW |
| | CD-1 | A/HongKong/1-11/1968(H3N2) | | 7.7 | | PFU | Value given | Avirulent | LOW |
| | CD-1 | A/HongKong/1-12/1968(H3N2) | | 7.7 | | PFU | Value given | Avirulent | LOW |
| | CD-1 | A/HongKong/1-1-MA-12/1968(H3N2) | 4 | | | PFU | Value given | Virulent | INTERMEDIATE |
| | CD-1 | A/HongKong/1-1-MA-12A/1968(H3N2) | 3.9 | | | PFU | Value given | Virulent | INTERMEDIATE |
| | CD-1 | A/HongKong/1-1-MA-12B/1968(H3N2) | 3.6 | | | PFU | Value given | Virulent | INTERMEDIATE |
| | CD-1 | A/HongKong/1-1-MA-12C/1968(H3N2) | 4.6 | | | PFU | Value given | Virulent | INTERMEDIATE |
| | CD-1 | A/HongKong/1-1-MA-12D/1968(H3N2) | 5.1 | | | PFU | Value given | Virulent | INTERMEDIATE |
| | CD-1 | A/HongKong/1-1-MA-12E/1968(H3N2) | 5.4 | | | PFU | Value given | Virulent | INTERMEDIATE |
| | CD-1 | A/HongKong/1-1-MA-20/1968(H3N2) | 2.9 | | | PFU | Value given | Virulent | HIGH |
| | CD-1 | A/HongKong/1-1-MA-20A/1968(H3N2) | 4 | | | PFU | Value given | Virulent | INTERMEDIATE |
| | CD-1 | A/HongKong/1-1-MA-20B/1968(H3N2) | 3 | | | PFU | Value given | Virulent | HIGH |
| | CD-1 | A/HongKong/1-1-MA-20C/1968(H3N2) | 2.6 | | | PFU | Value given | Virulent | HIGH |
| | CD-1 | A/HongKong/1-1-MA-20D/1968(H3N2) | 3.5 | | | PFU | Value given | Virulent | INTERMEDIATE |
| | CD-1 | A/HongKong/1-1-MA-20E/1968(H3N2) | 3.6 | | | PFU | Value given | Virulent | INTERMEDIATE |
| | CD-1 | A/HongKong/1-4-MA21-1/1968(H3N2) | 4.5 | | | PFU | Value given | Virulent | INTERMEDIATE |
| | CD-1 | A/HongKong/1-4-MA21-3/1968(H3N2) | 2.8 | | | PFU | Value given | Virulent | HIGH |
| | CD-1 | A/HongKong/1-5-MA21-1/1968(H3N2) | 5.5 | | | PFU | Value given | Virulent | INTERMEDIATE |
| | CD-1 | A/HongKong/1-5-MA21-3/1968(H3N2) | 5.5 | | | PFU | Value given | Virulent | INTERMEDIATE |

| | | | | | | | | | |
|---|---|---|---|---|---|---|---|---|---|
| | CD-1 | A/HongKong/1-6-MA21-3/1968(H3N2) | 1.1 | | | PFU | Value given | Virulent | HIGH |
| | CD-1 | A/HongKong/1-9-MA21-3/1968(H3N2) | 6.5 | | | PFU | Value given | Avirulent | LOW |
| | CD-1 | A/HongKong/1-11-MA21-2/1968(H3N2) | 5.8 | | | PFU | Value given | Virulent | INTERMEDIATE |
| Ping *et al.*, 2018 [56] | BALB/C | A/PuertoRico/8/1934(H1N1) | 1.74 | | | PFU | Value given | Virulent | HIGH |
| | BALB/C | rA/PR8(1235678)/CH1(4)(H1N1) | | 6.72 | | PFU | Value given | Avirulent | LOW |
| | BALB/C | A/Sichuan/1/2009(H1N1) | 3.73 | | | PFU | Value given | Virulent | INTERMEDIATE |
| | BALB/C | A/NewCaledonia/20/1999(H1N1) | 5.68 | | | PFU | Value given | Virulent | INTERMEDIATE |
| | BALB/C | A/FortMonmouth/1/1947(H1N1) | 1.18 | | | PFU | Value given | Virulent | HIGH |
| Qi *et al.*, 2009 [57] | BALB/C | A/NewYork/312/2001(H1N1) | | 5.3 | | PFU | Value given | Avirulent | LOW |
| | BALB/C | rA/NY312(123578)/ALB76(46)(H1N1) | | 5.3 | | PFU | Value given | Avirulent | LOW |
| | BALB/C | rA/NY312(123578)/mutALB76(46)(H1N1) | | 5.3 | | PFU | Value given | Avirulent | LOW |
| | BALB/C | rA/NY312(123578)/SC18(46)(H1N1) | 3.8 | | | PFU | Value given | Virulent | INTERMEDIATE |
| | BALB/C | rA/NY312(123578)/SC18(46)/HA-D225G(H1N1) | 4.47 | | | PFU | Value given | Virulent | INTERMEDIATE |
| | BALB/C | rA/NY312(123578)/SC18(46)/HA-D190E-D225G(H1N1) | | 5.3 | | PFU | Value given | Virulent | INTERMEDIATE |
| Qi *et al.*, 2012 [58] | BALB/C | A/SouthCarolina/1/1918(H1N1) | 2.1 | | | PFU | Value given | Virulent | HIGH |
| | BALB/C | rA/SC18(2345678)/OH175(1)(H1N1) | | 5 | | PFU | Value given | Avirulent | LOW |
| | BALB/C | rA/SC18(1345678)/OH175(2)(H1N1) | 1.8 | | | PFU | Value given | Virulent | HIGH |
| | BALB/C | rA/SC18(1245678)/OH175(3)(H1N1) | 1.8 | | | PFU | Value given | Virulent | HIGH |
| | BALB/C | rA/SC18(1235678)/OH265(4)(H1N1) | 1.4 | | | PFU | Value given | Virulent | HIGH |
| | BALB/C | rA/SC18(1234678)/OH175(5)(H1N1) | 2.1 | | | PFU | Value given | Virulent | HIGH |
| | BALB/C | rA/SC18(1234578)/OH175(6)(H1N1) | 2.7 | | | PFU | Value given | Virulent | HIGH |
| | BALB/C | rA/SC18(1234568)/OH175(7)(H1N1) | 2.7 | | | PFU | Value given | Virulent | HIGH |
| | BALB/C | rA/SC18(1234567)/OH175(8)(H1N1) | 2.3 | | | PFU | Value given | Virulent | HIGH |
| | BALB/C | rA/OH175(1235678)/OH265(4)(H1N1) | | 5 | | PFU | Value given | Avirulent | LOW |
| | BALB/C | rA/SC18/PB2-E627(H1N1) | | 5 | | PFU | Value given | Avirulent | LOW |
| | BALB/C | rA/SC18/OH175-PB2-K627(H1N1) | 1.8 | | | PFU | Value given | Virulent | HIGH |
| Qi *et al.*, 2018 [59] | BALB/C | A/chicken/Guangdong/SW154/2015(H7N9) | | 6 | | EID50 | LD50_LB or LD50_UB is based on survival rate | Avirulent | LOW |
| | BALB/C | A/chicken/Heyuan/16876/2016(H7N9) | | 6 | | EID50 | LD50_LB or LD50_UB is based on survival rate | Avirulent | LOW |
| | BALB/C | A/chicken/Huizhou/HZ-3/2016(H7N9) | | 6 | | EID50 | LD50_LB or LD50_UB is based on survival rate | Avirulent | LOW |
| | BALB/C | A/Guangdong/Th005/2017(H7N9) | | | 6 | EID50 | LD50_LB or LD50_UB is based on survival rate | Virulent | HIGH |
| | BALB/C | A/Guangdong/Th008/2017(H7N9) | | | 6 | EID50 | LD50_LB or LD50_UB is based on survival rate | Virulent | HIGH |
| Quan *et al.*, 2008 [60] | BALB/C | A/PuertoRico/8/1934(H1N1) | 2.3 | | | PFU | Value given | Virulent | HIGH |
| | BALB/C | rA/X-31(H3N2) | 5.85 | | | PFU | Value given | Virulent | INTERMEDIATE |
| | BALB/C | A/WSN/1933(H1N1) | 2.3 | | | PFU | Value given | Virulent | HIGH |

| | | | | | | | | | |
|---|---|---|---|---|---|---|---|---|---|
| | BALB/C | A/Philippines/2/1982(H3N2) | 2.3 | | | PFU | Value given | Virulent | HIGH |
| Rodriguez *et al*., 2013 [61] | BALB/C | A/CastillaLaMancha/RR5661/2009(H1N1) | | 6 | | PFU | LD50_LB or LD50_UB is based on survival rate | Avirulent | LOW |
| | BALB/C | A/CastillaLaMancha/RR5911/2009(H1N1) | 6 | | | PFU | LD50_LB or LD50_UB is based on survival rate | Virulent | INTERMEDIATE |
| Shi *et al*., 2017 [62] | BALB/C | A/chicken/Guangdong/SD008/2017(H7N9) | | 7.5 | | EID50 | Value given | Avirulent | LOW |
| | BALB/C | A/chicken/Guangdong/SD008/PB2-627K/2017(H7N9) | 1.8 | | | EID50 | Value given | Virulent | HIGH |
| | BALB/C | A/chicken/Guangdong/SD008/PB2-701N/2017(H7N9) | 3.4 | | | EID50 | Value given | Virulent | INTERMEDIATE |
| Smee *et al*., 2012 [63] | BALB/C | A/California/04/2009(H1N1) | 3.5 | | | CCID50 | Value given | Virulent | INTERMEDIATE |
| | BALB/C | A/NWS/1933(H1N1) | 4 | | | CCID50 | Value given | Virulent | INTERMEDIATE |
| | BALB/C | A/Victoria/3/1975(H3N2) | 4.5 | | | CCID50 | Value given | Virulent | INTERMEDIATE |
| | BALB/C | A/duck/Minnesota/1525/1981(H5N1) | 3.5 | | | CCID50 | Value given | Virulent | INTERMEDIATE |
| Song *et al*., 2009 [64] | BALB/C | mA/aquaticbird/Korea/w81/2005(H5N2) | 2.6 | | | TCID50 | Value given | Virulent | HIGH |
| | BALB/C | rA/MA-w81(H5N2) | 2.7 | | | TCID50 | Value given | Virulent | HIGH |
| | BALB/C | rA/MA-w81(2345678)/w81(1)(H5N2) | 3 | | | TCID50 | Value given | Virulent | HIGH |
| | BALB/C | rA/MA-w81(1345678)/w81(2)(H5N2) | 2.9 | | | TCID50 | Value given | Virulent | HIGH |
| | BALB/C | rA/MA-w81(1245678)/w81(3)(H5N2) | | 5.5 | | TCID50 | Value given | Avirulent | LOW |
| | BALB/C | rA/MA-w81(1235678)/w81(4)(H5N2) | 3 | | | TCID50 | Value given | Virulent | HIGH |
| | BALB/C | rA/MA-w81(1234578)/w81(6)(H5N2) | 3.5 | | | TCID50 | Value given | Virulent | INTERMEDIATE |
| | BALB/C | rA/MA-w81(1234568)/w81(7)(H5N2) | 2.4 | | | TCID50 | Value given | Virulent | HIGH |
| | BALB/C | rA/MA-w81/PB2-627K(H5N2) | 2 | | | TCID50 | Value given | Virulent | HIGH |
| | BALB/C | A/aquaticbird/Korea/w81/2005(H5N2) | | 5.5 | | TCID50 | Value given | Avirulent | LOW |
| | BALB/C | rA/aquaticbird/Korea/w81/2005(H5N2) | | 5.5 | | TCID50 | Value given | Avirulent | LOW |
| | BALB/C | rA/w81(2345678)/MA-w81(1)(H5N2) | | 5.5 | | TCID50 | Value given | Avirulent | LOW |
| | BALB/C | rA/w81(1345678)/MA-w81(2)(H5N2) | | 5.5 | | TCID50 | Value given | Avirulent | LOW |
| | BALB/C | rA/w81(1245678)/MA-w81(3)(H5N2) | 4.7 | | | TCID50 | Value given | Virulent | INTERMEDIATE |
| | BALB/C | rA/w81(1235678)/MA-w81(4)(H5N2) | | 5.5 | | TCID50 | Value given | Avirulent | LOW |
| | BALB/C | rA/w81(1234578)/MA-w81(6)(H5N2) | 5.4 | | | TCID50 | Value given | Virulent | INTERMEDIATE |
| | BALB/C | rA/w81(1234568)/MA-w81(7)(H5N2) | | 5.5 | | TCID50 | Value given | Avirulent | LOW |
| | BALB/C | rA/w81(124578)/MA-w81(36)(H5N2) | 4.1 | | | TCID50 | Value given | Virulent | INTERMEDIATE |
| | BALB/C | rA/MA-w81/PA-22K(H5N2) | 3.1 | | | TCID50 | Value given | Virulent | INTERMEDIATE |
| | BALB/C | rA/MA-w81/PA-97T(H5N2) | | 5.5 | | TCID50 | Value given | Avirulent | LOW |
| | BALB/C | rA/MA-w81/PA-155M(H5N2) | 2.5 | | | TCID50 | Value given | Virulent | HIGH |
| | BALB/C | rA/MA-w81/PA-216D(H5N2) | 2.7 | | | TCID50 | Value given | Virulent | HIGH |
| | BALB/C | rA/MA-w81/PA-22K-97T(H5N2) | | 5.5 | | TCID50 | Value given | Avirulent | LOW |
| | BALB/C | rA/MA-w81/PA-97T-155M(H5N2) | | 5.5 | | TCID50 | Value given | Avirulent | LOW |

| | | | | | | | | | |
|---|---|---|---|---|---|---|---|---|---|
| | BALB/C | rA/MA-w81/PA-97T-216D(H5N2) | | 5.5 | | TCID50 | Value given | Avirulent | LOW |
| | BALB/C | rA/MA-w81/w81PA-97I(H5N2) | 2.7 | | | TCID50 | Value given | Virulent | HIGH |
| | BALB/C | rA/w81/PA-97I(H5N2) | 4.5 | | | TCID50 | Value given | Virulent | INTERMEDIATE |
| | BALB/C | rA/w81/PA-97I/MA-w81NA(H5N2) | 3.8 | | | TCID50 | Value given | Virulent | INTERMEDIATE |
| | BALB/C | rA/w81/NA-106V/PA-97I(H5N2) | 4.8 | | | TCID50 | Value given | Virulent | INTERMEDIATE |
| | BALB/C | rA/w81/NA-316Y/PA-97I(H5N2) | 4.1 | | | TCID50 | Value given | Virulent | INTERMEDIATE |
| | BALB/C | rA/w81/NA-436A/PA-97I(H5N2) | 5 | | | TCID50 | Value given | Virulent | INTERMEDIATE |
| | BALB/C | rA/w81/PB2-627K(H5N2) | 4.3 | | | TCID50 | Value given | Virulent | INTERMEDIATE |
| | BALB/C | rA/w81/PB2-627K/PA-97I(H5N2) | 4 | | | TCID50 | Value given | Virulent | INTERMEDIATE |
| | BALB/C | rA/MA-w81/PB2-627K/PA-97T(H5N2) | 4.1 | | | TCID50 | Value given | Virulent | INTERMEDIATE |
| Song *et al.*, 2013 [65] | BALB/C | A/California/04/2009(H1N1) | | 5.5 | | TCID50 | Value given | Virulent | INTERMEDIATE |
| | BALB/C | maA/California/04A/2009(H1N1) | 2 | | | TCID50 | Value given | Virulent | HIGH |
| | BALB/C | maA/California/04B/2009(H1N1) | | | 4 | TCID50 | LD50_LB or LD50_UB is based on survival rate | Virulent | HIGH |
| | BALB/C | maA/California/04C/2009(H1N1) | | | 4 | TCID50 | LD50_LB or LD50_UB is based on survival rate | Virulent | HIGH |
| | BALB/C | maA/California/04D/2009(H1N1) | | 4 | | TCID50 | LD50_LB or LD50_UB is based on survival rate | Virulent | INTERMEDIATE |
| | BALB/C | A/California/04/H274Y/2009(H1N1) | | 5.5 | | TCID50 | Value given | Avirulent | LOW |
| | BALB/C | maA/California/04A/H274Y/2009(H1N1) | | | 4 | TCID50 | LD50_LB or LD50_UB is based on survival rate | Virulent | HIGH |
| | BALB/C | maA/California/04B/H274Y/2009(H1N1) | | | 4 | TCID50 | LD50_LB or LD50_UB is based on survival rate | Virulent | HIGH |
| | BALB/C | maA/California/04C/H274Y/2009(H1N1) | 1.5 | | | TCID50 | Value given | Virulent | HIGH |
| | BALB/C | maA/California/04D/H274Y/2009(H1N1) | | | 4 | TCID50 | LD50_LB or LD50_UB is based on survival rate | Virulent | HIGH |
| Srivastava *et al.*, 2009 [66] | C57BL/6 | A/PuertoRico/8/1934(H1N1) | 5.3 | | | FFU | Value given | Virulent | INTERMEDIATE |
| | C57BL/6 | A/seal/Massachussetts/1-SC35M/1980(H7N7) | 4.07 | | | FFU | Using formula Reed and Muench method | Virulent | INTERMEDIATE |
| | BALB/C | A/PuertoRico/8/1934(H1N1) | | 3.3 | | FFU | LD50_LB or LD50_UB is based on weight loss | Avirulent | LOW |
| | FVB/NJ | A/PuertoRico/8/1934(H1N1) | | 3.3 | | FFU | LD50_LB or LD50_UB is based on weight loss | Avirulent | LOW |
| | A/J | A/PuertoRico/8/1934(H1N1) | | | 3.3 | FFU | LD50_LB or LD50_UB is based on weight loss | Virulent | HIGH |
| | CBA/J | A/PuertoRico/8/1934(H1N1) | | 3.3 | | FFU | LD50_LB or LD50_UB is based on weight loss | Virulent | INTERMEDIATE |
| | SJL/JOrlCrl | A/PuertoRico/8/1934(H1N1) | | 3.3 | | FFU | LD50_LB or LD50_UB is based on weight loss | Avirulent | LOW |
| | DBA/2 | A/PuertoRico/8/1934(H1N1) | 1.56 | | | FFU | Value given | Virulent | HIGH |
| | DBA/2 | A/seal/Massachussetts/1-SC35M/1980(H7N7) | | | 3.3 | FFU | LD50_LB or LD50_UB is based on survival rate | Virulent | HIGH |
| Sun *et al.*, 2016 [67] | BALB/C | A/duck/EasternChina/S0711/2014(H5N6) | 5.3 | | | EID50 | Value given | Virulent | INTERMEDIATE |
| | BALB/C | A/duck/EasternChina/S0322/2014(H5N6) | 5 | | | EID50 | Value given | Virulent | INTERMEDIATE |
| | BALB/C | A/goose/EasternChina/S0513/2013(H5N6) | 5.3 | | | EID50 | Value given | Virulent | INTERMEDIATE |
| | BALB/C | A/duck/EasternChina/S0908/2014(H5N6) | 5.3 | | | EID50 | Value given | Virulent | INTERMEDIATE |
| Sutton *et al.*, 2017 [68] | BALB/C | A/Anhui/1/2013(H7N9) | 5 | | | TCID50 | Value given | Virulent | INTERMEDIATE |

| | | | | | | | | | |
|---|---|---|---|---|---|---|---|---|---|
| | BALB/C | rA/X-79(H3N2) | 1.2 | | | TCID50 | Value given | Virulent | HIGH |
| Tate *et al*., 2011a [69] | C57BL/6 | rA/X-31(H3N2) | | 5 | | PFU | LD50_LB or LD50_UB is based on weight loss | Avirulent | LOW |
| | C57BL/6 | rA/BJ89(123578)/PR8(46)(H1N1) | | 5 | | PFU | LD50_LB or LD50_UB is based on weight loss | Avirulent | LOW |
| | C57BL/6 | A/PuertoRico/8/1934(H1N1) | | | 5 | PFU | LD50_LB or LD50_UB is based on weight loss | Virulent | HIGH |
| Tate *et al*., 2011b [70] | C57BL/6 | A/PuertoRico/8/1934/MountSinai(H1N1) | | | 5 | PFU | LD50_LB or LD50_UB is based on weight loss | Virulent | HIGH |
| | C57BL/6 | A/Brazil/11/1978(H1N1) | | 5 | | PFU | LD50_LB or LD50_UB is based on weight loss | Avirulent | LOW |
| | C57BL/6 | rA/PR8vMountSinai(1235678)/BRAZ78(4)(H1N1) | | 5 | | PFU | LD50_LB or LD50_UB is based on weight loss | Avirulent | LOW |
| Tumpey *et al*., 2005 [71] | BALB/C | A/SouthCarolina/1/1918(H1N1) | 3.38 | | | PFU | Value given; Average LD50 from 10E3.25 and 10E3.5 | Virulent | INTERMEDIATE |
| | BALB/C | A/Texas/36/1991(H1N1) | | 6 | | PFU | LD50_LB or LD50_UB is based on survival rate | Avirulent | LOW |
| | BALB/C | rA/SC18(45678)/TX91(123)(H1N1) | 5.5 | | | PFU | Value given | Virulent | INTERMEDIATE |
| | BALB/C | rA/SC18(1235678)/TX91(4)(H1N1) | | 6 | | PFU | LD50_LB or LD50_UB is based on survival rate | Avirulent | LOW |
| Vasilijevic *et al*., 2017 [72] | BALB/C | rA/California/04/2009(H1N1) | 5 | | | PFU | Value given | Virulent | INTERMEDIATE |
| | BALB/C | rA/CA04/PB2mut(H1N1) | | 6 | | PFU | Value given | Avirulent | LOW |
| | BALB/C | rA/CA04/PAmut(H1N1) | 3.48 | | | PFU | Value given | Virulent | INTERMEDIATE |
| | BALB/C | rA/CA04/PB2-PAmut(H1N1) | 4.54 | | | PFU | Value given | Virulent | INTERMEDIATE |
| | BALB/C | rA/CA04/Mmut(H1N1) | | 5.7 | | PFU | Value given | Avirulent | LOW |
| | BALB/C | rA/CA04/M-PAmut(H1N1) | | 5.7 | | PFU | Value given | Avirulent | LOW |
| Wang *et al*., 2018 [73] | BALB/C | rA/GSH7/673HA-673NA(H5N6) | 2.83 | | | EID50 | Value given | Virulent | HIGH |
| | BALB/C | rA/GSH7/673HA-674NA(H5N6) | 4.5 | | | EID50 | Value given | Virulent | INTERMEDIATE |
| | BALB/C | rA/GSH7/674HA-673NA(H5N6) | 4.5 | | | EID50 | Value given | Virulent | INTERMEDIATE |
| | BALB/C | rA/GSH7/674HA-674NA(H5N6) | 3.2 | | | EID50 | Value given | Virulent | INTERMEDIATE |
| Yang *et al*., 2018 [74] | BALB/C | A/chicken/Hunan/S1220/2017(H7N9) | 3.2 | | | EID50 | Value given | Virulent | INTERMEDIATE |
| Ye *et al*., 2010 [75] | DBA/2 | A/California/04/2009(H1N1) | | | 5.7 | TCID50 | LD50_LB or LD50_UB is based on survival rate | Virulent | HIGH |
| | BALB/C | A/California/04/2009(H1N1) | | 6 | | TCID50 | Value given | Avirulent | LOW |
| | BALB/C | A/California/04/MA1/Y/2009(H1N1) | 3.3 | | | TCID50 | Value given | Virulent | INTERMEDIATE |
| | BALB/C | rA/CA04(1235678)/CA04MA1Y(4)(H1N1) | 6 | | | TCID50 | Value given | Virulent | INTERMEDIATE |
| | BALB/C | rA/CA04(125678)/CA04MA1Y(34)(H1N1) | 4.6 | | | TCID50 | Value given | Virulent | INTERMEDIATE |
| | BALB/C | rA/CA04(123678)/CA04MA1Y(45)(H1N1) | 4.8 | | | TCID50 | Value given | Virulent | INTERMEDIATE |
| | BALB/C | rA/CA04(12678)/CA04MA1Y(345)(H1N1) | 3.8 | | | TCID50 | Value given | Virulent | INTERMEDIATE |
| | BALB/C | A/Netherlands/602/2009(H1N1) | 6 | | | TCID50 | Value given | Virulent | INTERMEDIATE |
| | BALB/C | rA/NL602(1235678)/NY18(4)(H1N1) | | 6 | | TCID50 | Value given | Avirulent | LOW |
| | BALB/C | rA/NL602(1235678)/CA04MA1Y(4)(H1N1) | 3.4 | | | TCID50 | Value given | Virulent | INTERMEDIATE |
| Yu *et al*., 2017 [76] | BALB/C | rA/goose/Guangdong/SH7/2013(H5N1) | 2.5 | | | EID50 | Value given | Virulent | HIGH |
| | BALB/C | rA/GSH7/LZFNA(H5N2) | 3.3 | | | EID50 | Value given | Virulent | INTERMEDIATE |

| | | | | | | | | | |
|---|---|---|---|---|---|---|---|---|---|
| | BALB/C | rA/GSH7/673NA(H5N6) | 2.68 | | | EID50 | Value given | Virulent | HIGH |
| | BALB/C | rA/GSH7/674NA(H5N6) | 3.16 | | | EID50 | Value given | Virulent | INTERMEDIATE |
| | BALB/C | rA/GSH7/JS1306NA(H5N8) | 2.5 | | | EID50 | Value given | Virulent | HIGH |
| Yu et al., 2018a [77] | BALB/C | A/duck/Liaoning/LN/2011(H5N5) | 6.25 | | | EID50 | Value given | Avirulent | LOW |
| | BALB/C | mA/duck/Liaoning/LNP1/2011(H5N5) | 5.75 | | | EID50 | Value given | Virulent | INTERMEDIATE |
| | BALB/C | mA/duck/Liaoning/LNP2/2011(H5N5) | 2.75 | | | EID50 | Value given | Virulent | HIGH |
| Yu *et al*., 2018b [78] | BALB/C | A/mallard/Shanghai/SH-9/2013(H5N8) | 5.75 | | | EID50 | Value given | Virulent | INTERMEDIATE |
| | BALB/C | mA/Shanghai/SH-9/L1P5/2013(H5N8) | 1.25 | | | EID50 | Value given | Virulent | HIGH |
| | BALB/C | mA/Shanghai/SH-9/L2P5/2013(H5N8) | 1.5 | | | EID50 | Value given | Virulent | HIGH |
| | BALB/C | mA/Shanghai/SH-9/L1P2/2013(H5N8) | 5.5 | | | EID50 | Value given | Virulent | INTERMEDIATE |
| | BALB/C | mA/Shanghai/SH-9/L1P4/2013(H5N8) | 2.5 | | | EID50 | Value given | Virulent | HIGH |
| Zhang *et al*., 2013 [79] | BALB/C | A/chicken/Shanghai/S1053/2013(H7N9) | | 6 | | EID50 | LD50_LB or LD50_UB is based on survival rate | Avirulent | LOW |
| | BALB/C | A/pigeon/Shanghai/S1069/2013(H7N9) | | 6 | | EID50 | LD50_LB or LD50_UB is based on survival rate | Avirulent | LOW |
| | BALB/C | A/pigeon/Shanghai/S1421/2013(H7N9) | | 6 | | EID50 | LD50_LB or LD50_UB is based on survival rate | Avirulent | LOW |
| | BALB/C | A/Shanghai/1/2013(H7N9) | 5.4 | | | EID50 | | Virulent | INTERMEDIATE |
| | BALB/C | A/Shanghai/02/2013(H7N9) | | 6 | | EID50 | LD50_LB or LD50_UB is based on survival rate | Avirulent | LOW |
| | BALB/C | A/Anhui/1/2013(H7N9) | | 6 | | EID50 | LD50_LB or LD50_UB is based on survival rate | Avirulent | LOW |
| Zhou *et al*., 2013 [80] | BALB/C | rA/NewYork/1682/2009(H1N1) | | 6.78 | | TCID50 | Value given | Avirulent | LOW |
| | BALB/C | rA/NY1682/PB2-E158G(H1N1) | 2.78 | | | TCID50 | Value given | Virulent | HIGH |
| | BALB/C | rA/NY1682/PB2-E627K(H1N1) | | 6.78 | | TCID50 | Value given | Avirulent | LOW |
| | BALB/C | rA/NY1682/PB2-D701N(H1N1) | 4.3 | | | TCID50 | Value given | Virulent | INTERMEDIATE |
| Zhou *et al*., 2016 [81] | C57BL/6 | A/PuertoRico/8/1934(H1N1) | | 6 | | PFU | LD50_LB or LD50_UB is based on survival rate | Avirulent | LOW |
| | BALB/C | A/PuertoRico/8/1934(H1N1) | | | 6 | PFU | LD50_LB or LD50_UB is based on survival rate | Virulent | HIGH |
| | C3H | A/PuertoRico/8/1934(H1N1) | | | 6 | PFU | LD50_LB or LD50_UB is based on survival rate | Virulent | HIGH |
| | 129S1/SvPasCrlVr | A/PuertoRico/8/1934(H1N1) | | | 6 | PFU | LD50_LB or LD50_UB is based on survival rate | Virulent | HIGH |
| | DBA/2 | A/PuertoRico/8/1934(H1N1) | | | 6 | PFU | LD50_LB or LD50_UB is based on survival rate | Virulent | INTERMEDIATE |
| | FVB/NJ | A/PuertoRico/8/1934(H1N1) | | 6 | | PFU | LD50_LB or LD50_UB is based on survival rate | Avirulent | LOW |
| Zhu *et al*., 2013 [82] | C57BL/6 | A/Shanghai/4664T/2013(H7N9) | | | 5.6 | TCID50 | LD50_LB or LD50_UB is based on weight loss | Virulent | INTERMEDIATE |
| | BALB/C | A/Shanghai/4664T/2013(H7N9) | | | 5.6 | TCID50 | LD50_LB or LD50_UB is based on weight loss | Virulent | HIGH |
| | ICR | A/Shanghai/4664T/2013(H7N9) | | | 5.6 | TCID50 | LD50_LB or LD50_UB is based on weight loss | Virulent | INTERMEDIATE |
| Zhu *et al*., 2015 [83] | C57BL/6 | A/Anhui/1/2013(H7N9) | 4.5 | | | TCID50 | Value given | Virulent | INTERMEDIATE |
| | C57BL/6 | A/AH1/KN(H7N9) | 3.3 | | | TCID50 | Value given | Virulent | INTERMEDIATE |
| | C57BL/6 | A/AH1/EN(H7N9) | | 6.3 | | TCID50 | Value given | Avirulent | LOW |

| | | | | | | | | | |
|---|---|---|---|---|---|---|---|---|---|
| Zhu *et al*., 2016 [84] | C57BL/6 | A/Hunan/42443/2015(H1N1) | 4.5 | | | TCID50 | Value given; It is written as MID50 in the paper | Virulent | INTERMEDIATE |
| | C57BL/6 | A/Jiangsu/1/2011(H1N1) | | 6.5 | | TCID50 | Value given; It is written as MID50 in the paper | Avirulent | LOW |

**Table S2.** Reduction of multiple records for infection involving specific influenza A virus (IAV) and mouse strains into a single record. The records colored in red are the records being selected, with their LD50 values highlighted in bold if they are updated.

| Reference | Host strain | Influenza strain | LD50 point estimate | LD50 lower bound | LD50 upper bound | Infection unit | Two-class virulence level | Three-class virulence level |
|---|---|---|---|---|---|---|---|---|
| Bi *et al.*, 2015 [4] | BALB/C | A/Anhui/1/2013(H7N9) | 7.83 | | | EID50 | LOW | Avirulent |
| Imai *et al.*, 2017 [24] | BALB/C | A/Anhui/1/2013(H7N9) | 4.5 | | | PFU | INTERMEDIATE | Virulent |
| Sutton *et al.*, 2017 [68] | BALB/C | A/Anhui/1/2013(H7N9) | 5 | | | TCID50 | INTERMEDIATE | Virulent |
| Zhang *et al.*, 2013 [79] | BALB/C | A/Anhui/1/2013(H7N9) | | 6 | | EID50 | LOW | Avirulent |
| Song *et al.*, 2009 [64] | BALB/C | A/aquaticbird/Korea/w81/2005(H5N2) | | 5.5 | | TCID50 | LOW | Avirulent |
| Song *et al.*, 2009 [64] | BALB/C | A/aquaticbird/Korea/w81/2005(H5N2) | | 5.5 | | TCID50 | LOW | Avirulent |
| Lee *et al.*, 2018 [36] | BALB/C | A/broilerduck/Korea/Buan2/2014(H5N8) | 2.7 | | | TCID50 | HIGH | Virulent |
| Lee *et al.*, 2018 [36] | BALB/C | A/broilerduck/Korea/Buan2/2014(H5N8) | 3.4 | | | TCID50 | INTERMEDIATE | Virulent |
| Belser *et al.*, 2010 [3] | BALB/C | A/California/04/2009(H1N1) | | 6 | | EID50 | LOW | Avirulent |
| Cline *et al.*, 2011 [10] | BALB/C | A/California/04/2009(H1N1) | | 5 | | TCID50 | INTERMEDIATE | Virulent |
| Ilyushina *et al.*, 2010 [23] | BALB/C | A/California/04/2009(H1N1) | 5.1 | | | PFU | INTERMEDIATE | Virulent |
| Itoh *et al.*, 2009 [25] | BALB/C | A/California/04/2009(H1N1) | 5.8 | | | PFU | INTERMEDIATE | Virulent |
| Manicassamy *et al.*, 2010 [44] | BALB/C | A/California/04/2009(H1N1) | 4.7 | | | PFU | INTERMEDIATE | Virulent |
| Smee *et al.*, 2012 [63] | BALB/C | A/California/04/2009(H1N1) | 3.5 | | | CCID50 | INTERMEDIATE | Virulent |
| Song *et al.*, 2013 [65] | BALB/C | A/California/04/2009(H1N1) | | 5.5 | | TCID50 | INTERMEDIATE | Virulent |
| Vasilijevic *et al.*, 2017 [72] | BALB/C | A/California/04/2009(H1N1) | 5 | | | PFU | INTERMEDIATE | Virulent |
| Ye *et al.*, 2010 [75] | BALB/C | A/California/04/2009(H1N1) | | 6 | | TCID50 | LOW | Avirulent |
| Fan *et al.*, 2009 [15] | BALB/C | A/duck/Fujian/01/2002(H5N1) | 0.5 | | | EID50 | HIGH | Virulent |
| Fan *et al.*, 2009 [15] | BALB/C | A/duck/Fujian/01/2002(H5N1) | 0.9 | | | EID50 | HIGH | Virulent |
| Jiao *et al.*, 2008 [30] | BALB/C | A/duck/Guangxi/12/2003(H5N1) | 6.4 | | | EID50 | LOW | Avirulent |
| Jiao *et al.*, 2008 [30] | BALB/C | A/duck/Guangxi/12/2003(H5N1) | 6.4 | | | EID50 | LOW | Avirulent |
| Li *et al.*, 2005 [38] | BALB/C | A/duck/Guangxi/22/2001(H5N1) | | 6.5 | | EID50 | LOW | Avirulent |
| Li *et al.*, 2005 [38] | BALB/C | A/duck/Guangxi/22/2001(H5N1) | | 6.5 | | EID50 | LOW | Avirulent |
| Jiao *et al.*, 2008 [30] | BALB/C | A/duck/Guangxi/27/2003(H5N1) | 0.6 | | | EID50 | HIGH | Virulent |
| Jiao *et al.*, 2008 [30] | BALB/C | A/duck/Guangxi/27/2003(H5N1) | 0.6 | | | EID50 | HIGH | Virulent |
| Li *et al.*, 2005 [38] | BALB/C | A/duck/Guangxi/35/2001(H5N1) | 1.8 | | | EID50 | HIGH | Virulent |
| Li *et al.*, 2005 [38] | BALB/C | A/duck/Guangxi/35/2001(H5N1) | 2.3 | | | EID50 | HIGH | Virulent |
| Fan *et al.*, 2009 [15] | BALB/C | A/duck/Guangxi/53/2002(H5N1) | 6.5 | | | EID50 | LOW | Avirulent |
| Fan *et al.*, 2009 [15] | BALB/C | A/duck/Guangxi/53/2002(H5N1) | 6.4 | | | EID50 | LOW | Avirulent |

| | | | | | | | | |
|---|---|---|---|---|---|---|---|---|
| O'Neill *et al.*, 2000 [50] | BALB/C | A/goose/HongKong/437-6/1999(H5N1) | | 4.2 | | EID50 | LOW | Avirulent |
| O'Neill *et al.*, 2000 [50] | BALB/C | A/goose/HongKong/437-6/1999(H5N1) | | 4.2 | | EID50 | LOW | Avirulent |
| Imai *et al.*, 2017 [24] | BALB/C | A/Guangdong/Th008/2017(H7N9) | 2.7 | | | PFU | HIGH | Virulent |
| Qi *et al.*, 2018 [59] | BALB/C | A/Guangdong/Th008/2017(H7N9) | | | 6 | EID50 | HIGH | Virulent |
| O'Neill *et al.*, 2000 [50] | BALB/C | A/HongKong/1073/1999(H9N2) | | 4.2 | | EID50 | INTERMEDIATE | Virulent |
| O'Neill *et al.*, 2000 [50] | BALB/C | A/HongKong/1073/1999(H9N2) | | 4.2 | | EID50 | INTERMEDIATE | Virulent |
| Katz *et al.*, 2000 [32] | BALB/C | A/HongKong/156/1997(H5N1) | 5.9 | | | EID50 | INTERMEDIATE | Virulent |
| Lu *et al.*, 1999 [41] | BALB/C | A/HongKong/156/1997(H5N1) | 5.9 | | | EID50 | INTERMEDIATE | Virulent |
| O'Neill *et al.*, 2000 [50] | BALB/C | A/HongKong/156/1997(H5N1) | | | 4.2 | EID50 | HIGH | Virulent |
| Chen *et al.*, 2007 [8] | BALB/C | A/HongKong/483/1997(H5N1) | 1.9 | | | EID50 | HIGH | Virulent |
| Cline *et al.*, 2011 [10] | BALB/C | A/HongKong/483/1997(H5N1) | 1.5 | | | TCID50 | HIGH | Virulent |
| Hatta *et al.*, 2001 [21] | BALB/C | A/HongKong/483/1997(H5N1) | 0.26 | | | PFU | HIGH | Virulent |
| Hatta *et al.*, 2001 [21] | BALB/C | A/HongKong/483/1997(H5N1) | 0.23 | | | PFU | HIGH | Virulent |
| Katz *et al.*, 2000 [32] | BALB/C | A/HongKong/483/1997(H5N1) | 2.4 | | | EID50 | HIGH | Virulent |
| Lu *et al.*, 1999 [41] | BALB/C | A/HongKong/483/1997(H5N1) | 2.4 | | | EID50 | HIGH | Virulent |
| Maines *et al.*, 2005 [43] | BALB/C | A/HongKong/483/1997(H5N1) | 1.6 | | | EID50 | HIGH | Virulent |
| Katz *et al.*, 2000 [32] | BALB/C | A/HongKong/485/1997(H5N1) | 2.9 | | | EID50 | HIGH | Virulent |
| Lu *et al.*, 1999 [41] | BALB/C | A/HongKong/485/1997(H5N1) | 2.9 | | | EID50 | HIGH | Virulent |
| Chen *et al.*, 2007 [8] | BALB/C | A/HongKong/486/1997(H5N1) | 5.98 | | | EID50 | INTERMEDIATE | Virulent |
| Hatta *et al.*, 2001 [21] | BALB/C | A/HongKong/486/1997(H5N1) | | 3.88 | | PFU | LOW | Avirulent |
| Hatta *et al.*, 2001 [21] | BALB/C | A/HongKong/486/1997(H5N1) | 4.66 | | | PFU | INTERMEDIATE | Virulent |
| Hatta *et al.*, 2001 [21] | BALB/C | A/HongKong/486/1997(H5N1) | 4 | | | PFU | INTERMEDIATE | Virulent |
| Katz *et al.*, 2000 [32] | BALB/C | A/HongKong/486/1997(H5N1) | | 6.5 | | EID50 | LOW | Avirulent |
| Lu *et al.*, 1999 [41] | BALB/C | A/HongKong/486/1997(H5N1) | | 7 | | EID50 | LOW | Avirulent |
| Casalegno *et al.*, 2014 [7] | BALB/C | A/Lyon/969/2009(H1N1) | 3.2 | | | TCID50 | INTERMEDIATE | Virulent |
| Ferraris *et al.*, 2012 [16] | BALB/C | A/Lyon/969/2009(H1N1) | | 6 | | TCID50 | LOW | Avirulent |
| Belser *et al.*, 2007 [2] | BALB/C | A/Netherlands/219/2003(H7N7) | 2.5 | | | EID50 | HIGH | Virulent |
| Joseph *et al.*, 2007 [31] | BALB/C | A/Netherlands/219/2003(H7N7) | 0.8 | | | TCID50 | HIGH | Virulent |
| Jang *et al.*, 2014 [28] | BALB/C | A/NewCaledonia/20/1999(H1N1) | 6 | | | PFU | INTERMEDIATE | Virulent |
| Ping *et al.*, 2018 [56] | BALB/C | A/NewCaledonia/20/1999(H1N1) | 5.68 | | | PFU | INTERMEDIATE | Virulent |
| Jang *et al.*, 2018 [29] | BALB/C | A/Philippines/2/1982(H3N2) | 4.7 | | | PFU | INTERMEDIATE | Virulent |
| Quan *et al.*, 2008 [60] | BALB/C | A/Philippines/2/1982(H3N2) | 2.3 | | | PFU | HIGH | Virulent |
| Jang *et al.*, 2018 [29] | BALB/C | A/PuertoRico/8/1934(H1N1) | 3.7 | | | PFU | INTERMEDIATE | Virulent |
| Liedmann *et al.*, 2014 [39] | BALB/C | A/PuertoRico/8/1934(H1N1) | | | 2.5 | PFU | HIGH | Virulent |

| | | | | | | | | |
|---|---|---|---|---|---|---|---|---|
| Ping *et al*., 2018 [56] | BALB/C | A/PuertoRico/8/1934(H1N1) | 1.74 | | | PFU | HIGH | Virulent |
| Quan *et al*., 2008 [60] | BALB/C | A/PuertoRico/8/1934(H1N1) | 2.3 | | | PFU | HIGH | Virulent |
| Srivastava *et al*., 2009 [66] | BALB/C | A/PuertoRico/8/1934(H1N1) | | 3.3 | | FFU | LOW | Avirulent |
| Zhou *et al*., 2016 [81] | BALB/C | A/PuertoRico/8/1934(H1N1) | | | 6 | PFU | HIGH | Virulent |
| Belser *et al*., 2010 [3] | BALB/C | A/SouthCarolina/1/1918(H1N1) | 3.5 | | | EID50 | INTERMEDIATE | Virulent |
| Qi *et al*., 2012 [58] | BALB/C | A/SouthCarolina/1/1918(H1N1) | 2.1 | | | PFU | HIGH | Virulent |
| Tumpey *et al*., 2005 [71] | BALB/C | A/SouthCarolina/1/1918(H1N1) | 3.38 | | | PFU | INTERMEDIATE | Virulent |
| Belser *et al*., 2010 [3] | BALB/C | A/VietNam/1203/2004(H5N1) | 1.3 | | | EID50 | HIGH | Virulent |
| Maines *et al*., 2005 [43] | BALB/C | A/VietNam/1203/2004(H5N1) | 2.2 | | | EID50 | HIGH | Virulent |
| Kobasa *et al*., 2004 [34] | BALB/C | A/WSN/1933(H1N1) | 3.3 | | | PFU | INTERMEDIATE | Virulent |
| Quan *et al*., 2008 [60] | BALB/C | A/WSN/1933(H1N1) | 2.3 | | | PFU | HIGH | Virulent |
| Jang *et al*., 2018 [29] | BALB/C | mA/aquaticbird/Korea/w81/2005(H5N2) | 4 | | | PFU | INTERMEDIATE | Virulent |
| Song *et al*., 2009 [64] | BALB/C | mA/aquaticbird/Korea/w81/2005(H5N2) | 2.6 | | | TCID50 | HIGH | Virulent |
| Chen *et al*., 2007 [8] | BALB/C | rA/HK486/PB2-627K(H5N1) | 2.25 | | | EID50 | HIGH | Virulent |
| Hatta *et al*., 2001 [21] | BALB/C | rA/HK486/PB2-627K(H5N1) | 0.76 | | | PFU | HIGH | Virulent |
| Jang *et al*., 2013b [27] | BALB/C | rA/X-31(H3N2) | | 5 | | PFU | LOW | Avirulent |
| Lu *et al*., 1999 [41] | BALB/C | rA/X-31(H3N2) | | 5.2 | | EID50 | LOW | Avirulent |
| Quan *et al*., 2008 [60] | BALB/C | rA/X-31(H3N2) | 5.85 | | | PFU | INTERMEDIATE | Virulent |
| Otte *et al*., 2011 [51] | C57BL/6 | A/Hamburg/05/2009(H1N1) | 5.2 | | | PFU | INTERMEDIATE | Virulent |
| Otte *et al*., 2015 [52] | C57BL/6 | A/Hamburg/05/2009(H1N1) | 5.2 | | | PFU | INTERMEDIATE | Virulent |
| Otte *et al*., 2015 [52] | C57BL/6 | A/Hamburg/05/2009(H1N1) | 5.2 | | | PFU | INTERMEDIATE | Virulent |
| Otte *et al*., 2011 [51] | C57BL/6 | A/Hamburg/NY1580/2009(H1N1) | 3.5 | | | PFU | INTERMEDIATE | Virulent |
| Otte *et al*., 2015 [52] | C57BL/6 | A/Hamburg/NY1580/2009(H1N1) | 3.2 | | | PFU | INTERMEDIATE | Virulent |
| Otte *et al*., 2015 [52] | C57BL/6 | A/Hamburg/NY1580/2009(H1N1) | 3.5 | | | PFU | INTERMEDIATE | Virulent |
| Manicassamy *et al*., 2010 [44] | C57BL/6 | A/Netherlands/602/2009(H1N1) | 4.2 | | | PFU | INTERMEDIATE | Virulent |
| Pica *et al*., 2011 [54] | C57BL/6 | A/Netherlands/602/2009(H1N1) | 4.3 | | | PFU | INTERMEDIATE | Virulent |
| Blazejewska *et al*., 2011 [5] | C57BL/6 | A/PR8F/1934(H1N1) | | | 3.3 | PFU | HIGH | Virulent |
| Hatesuer *et al*., 2013 [20] | C57BL/6 | A/PR8F/1934(H1N1) | | | 3.3 | FFU | HIGH | Virulent |
| Blazejewska *et al*., 2011 [5] | C57BL/6 | A/PR8M/1934(H1N1) | **4.3** | 3.3 | | PFU | INTERMEDIATE | Virulent |
| Hatesuer *et al*., 2013 [20] | C57BL/6 | A/PR8M/1934(H1N1) | | | 5.3 | FFU | INTERMEDIATE | Virulent |
| Liedmann *et al*., 2014 [39] | C57BL/6 | A/PuertoRico/8/1934(H1N1) | | | 3 | PFU | HIGH | Virulent |
| Na *et al*., 2016 [48] | C57BL/6 | A/PuertoRico/8/1934(H1N1) | 3.9 | | | EID50 | INTERMEDIATE | Virulent |
| Pica *et al*., 2011 [54] | C57BL/6 | A/PuertoRico/8/1934(H1N1) | 1.4 | | | PFU | HIGH | Virulent |
| Srivastava *et al*., 2009 [66] | C57BL/6 | A/PuertoRico/8/1934(H1N1) | 5.3 | | | FFU | INTERMEDIATE | Virulent |

| | | | | | | | | |
|---|---|---|---|---|---|---|---|---|
| Tate *et al*., 2011a [69] | C57BL/6 | A/PuertoRico/8/1934(H1N1) | | | 5 | PFU | HIGH | Virulent |
| Zhou *et al*., 2016 [81] | C57BL/6 | A/PuertoRico/8/1934(H1N1) | | 6 | | PFU | LOW | Avirulent |
| Hatesuer *et al*., 2013 [20] | C57BL/6 | A/seal/Massachussetts/1-SC35M/1980(H7N7) | | | 4.3 | FFU | HIGH | Virulent |
| Nurnberger *et al*., 2016 [49] | C57BL/6 | A/seal/Massachussetts/1-SC35M/1980(H7N7) | | | 3 | PFU | HIGH | Virulent |
| Srivastava *et al*., 2009 [66] | C57BL/6 | A/seal/Massachussetts/1-SC35M/1980(H7N7) | 4.07 | | | FFU | INTERMEDIATE | Virulent |
| Otte *et al*., 2011 [51] | C57BL/6 | A/SolomonIslands/3/2006(H1N1) | | 6 | | PFU | LOW | Avirulent |
| Pica *et al*., 2011 [54] | C57BL/6 | A/SolomonIslands/3/2006(H1N1) | | 5.3 | | PFU | LOW | Avirulent |
| Hatesuer *et al*., 2013 [20] | C57BL/6 | maA/HongKong/1/1968(H3N2) | **1** | 1 | | FFU | HIGH | Virulent |
| Leist *et al*., 2016 [37] | C57BL/6 | maA/HongKong/1/1968(H3N2) | | | 1 | FFU | HIGH | Virulent |
| Pica *et al*., 2011 [54] | C57BL/6 | rA/X-31(H3N2) | 5.3 | | | PFU | INTERMEDIATE | Virulent |
| Tate *et al*, 2011a [69] | C57BL/6 | rA/X-31(H3N2) | | 5 | | PFU | LOW | Avirulent |
| Liedmann *et al*., 2014 [39] | DBA/2 | A/PuertoRico/8/1934(H1N1) | | | 1 | PFU | HIGH | Virulent |
| Pica *et al*., 2011 [54] | DBA/2 | A/PuertoRico/8/1934(H1N1) | 0.4 | | | PFU | HIGH | Virulent |
| Srivastava *et al*., 2009 [66] | DBA/2 | A/PuertoRico/8/1934(H1N1) | 1.56 | | | FFU | HIGH | Virulent |
| Zhou *et al*., 2016 [81] | DBA/2 | A/PuertoRico/8/1934(H1N1) | | | 6 | PFU | INTERMEDIATE | Virulent |
| Srivastava *et al*., 2009 [66] | FVB/NJ | A/PuertoRico/8/1934(H1N1) | | 3.3 | | FFU | LOW | Avirulent |
| Zhou *et al*., 2016 [81] | FVB/NJ | A/PuertoRico/8/1934(H1N1) | | 6 | | PFU | LOW | Avirulent |

**Table S3.** Reduction of multiple records for infection of a specific influenza A virus (IAV) strain in different mouse strains into a single record. The records colored in red are the records being selected, with their LD50 values highlighted in bold if they are updated.

| Reference | Host strain | Influenza strain | LD50 point estimate | LD50 lower bound | LD50 upper bound | Infection unit | Two-class virulence level | Three-class virulence level |
|---|---|---|---|---|---|---|---|---|
| Imai *et al.*, 2017 [24] | BALB/C | A/Anhui/1/2013(H7N9) | 4.5 | | | PFU | INTERMEDIATE | Virulent |
| Zhu *et al.*, 2015 [83] | C57BL/6 | A/Anhui/1/2013(H7N9) | 4.5 | | | TCID50 | INTERMEDIATE | Virulent |
| Pica *et al.*, 2011 [54] | C57BL/6 | A/Brisbane/10/2007(H3N2) | | 6 | | PFU | LOW | Avirulent |
| Pica *et al.*, 2011 [54] | DBA/2 | A/Brisbane/10/2007(H3N2) | | 6 | | PFU | LOW | Avirulent |
| Pica *et al.*, 2011 [54] | C57BL/6 | A/Brisbane/59/2007(H1N1) | | 6 | | PFU | LOW | Avirulent |
| Pica *et al.*, 2011 [54] | DBA/2 | A/Brisbane/59/2007(H1N1) | 5 | | | PFU | INTERMEDIATE | Virulent |
| Manicassamy *et al.*, 2010 [44] | C57BL/6 | A/California/04/2009(H1N1) | 4.7 | | | PFU | INTERMEDIATE | Virulent |
| Smee *et al.*, 2012 [63] | BALB/C | A/California/04/2009(H1N1) | 3.5 | | | CCID50 | INTERMEDIATE | Virulent |
| Ye *et al.*, 2010 [75] | DBA/2 | A/California/04/2009(H1N1) | | | 5.7 | TCID50 | HIGH | Virulent |
| Pica *et al.*, 2011 [54] | C57BL/6 | A/duck/Alberta/35/1976(H1N1) | | 6 | | PFU | LOW | Avirulent |
| Pica *et al.*, 2011 [54] | DBA/2 | A/duck/Alberta/35/1976(H1N1) | | 6 | | PFU | LOW | Avirulent |
| Pica *et al.*, 2011 [54] | C57BL/6 | A/duck/Ukraine/1/1963(H3N8) | | 6 | | PFU | LOW | Avirulent |
| Pica *et al.*, 2011 [54] | DBA/2 | A/duck/Ukraine/1/1963(H3N8) | | 6 | | PFU | LOW | Avirulent |
| Choi *et al.*, 2017 [9] | C57BL/6 | A/environment/Korea/W468/2014(H5N8) | | 4 | | PFU | LOW | Avirulent |
| Choi *et al.*, 2017 [9] | BALB/C | A/environment/Korea/W468/2014(H5N8) | 7.3 | | | PFU | LOW | Avirulent |
| Otte *et al.*, 2011 [51] | BALB/C | A/Hamburg/05/2009(H1N1) | | 6 | | PFU | LOW | Avirulent |
| Otte *et al.*, 2011 [51] | C57BL/6 | A/Hamburg/05/2009(H1N1) | 5.2 | | | PFU | INTERMEDIATE | Virulent |
| Otte *et al.*, 2011 [51] | BALB/C | A/Hamburg/NY1580/2009(H1N1) | | 6 | | PFU | LOW | Avirulent |
| Otte *et al.*, 2015 [52] | C57BL/6 | A/Hamburg/NY1580/2009(H1N1) | 3.2 | | | PFU | INTERMEDIATE | Virulent |
| Pica *et al.*, 2011 [54] | C57BL/6 | A/HongKong/1/1968(H3N2) | | 6 | | PFU | LOW | Avirulent |
| Pica *et al.*, 2011 [54] | DBA/2 | A/HongKong/1/1968(H3N2) | 5.5 | | | PFU | INTERMEDIATE | Virulent |
| Ping *et al.*, 2011 [55] | CD-1 | A/HongKong/1/1968(H3N2) | | 7.7 | | PFU | LOW | Avirulent |
| Lu *et al.*, 1999 [41] | BALB/C | A/HongKong/156/1997(H5N1) | 5.9 | | | EID50 | INTERMEDIATE | Virulent |
| O'Neill *et al.*, 2000 [50] | C57BL/6 | A/HongKong/156/1997(H5N1) | | | 2.8 | EID50 | HIGH | Virulent |
| Blazejewska *et al.*, 2011 [5] | C57BL/6 | A/HongKong/156/1997(H5N1) | | | 3.3 | PFU | HIGH | Virulent |
| Blazejewska *et al.*, 2011 [5] | DBA/2 | A/hvPR8/1934(H1N1) | | | 3.3 | PFU | HIGH | Virulent |
| Kim *et al.*, 2013 [33] | BALB/C | A/Korea/01/2009(H1N1) | | 6 | | PFU | LOW | Avirulent |
| Kim *et al.*, 2013 [33] | DBA/2 | A/Korea/01/2009(H1N1) | 2.83 | | | PFU | HIGH | Virulent |

| | | | | | | | | |
|---|---|---|---|---|---|---|---|---|
| Choi *et al.*, 2017 [9] | C57BL/6 | A/ma452-G1-1/2014(H5N8) | | | 4 | PFU | HIGH | Virulent |
| Choi *et al.*, 2017 [9] | BALB/C | A/ma452-G1-1/2014(H5N8) | 1 | | | PFU | HIGH | Virulent |
| Choi *et al.*, 2017 [9] | C57BL/6 | A/ma452-G3-1/2014(H5N8) | | | 4 | PFU | HIGH | Virulent |
| Choi *et al.*, 2017 [9] | BALB/C | A/ma452-G3-1/2014(H5N8) | 2 | | | PFU | HIGH | Virulent |
| Choi *et al.*, 2017 [9] | C57BL/6 | A/ma452-G3-2/2014(H5N8) | | | 4 | PFU | HIGH | Virulent |
| Choi *et al.*, 2017 [9] | BALB/C | A/ma452-G3-2/2014(H5N8) | 0.5 | | | PFU | HIGH | Virulent |
| Choi *et al.*, 2017 [9] | C57BL/6 | A/ma452-G4-1/2014(H5N8) | | | 4 | PFU | HIGH | Virulent |
| Choi *et al.*, 2017 [9] | BALB/C | A/ma452-G4-1/2014(H5N8) | 1.3 | | | PFU | HIGH | Virulent |
| Choi *et al.*, 2017 [9] | C57BL/6 | A/ma468-G1-1/2014(H5N8) | | | 4 | PFU | HIGH | Virulent |
| Choi *et al.*, 2017 [9] | BALB/C | A/ma468-G1-1/2014(H5N8) | 1.7 | | | PFU | HIGH | Virulent |
| Choi *et al.*, 2017 [9] | C57BL/6 | A/ma468-G1-2/2014(H5N8) | | | 4 | PFU | HIGH | Virulent |
| Choi *et al.*, 2017 [9] | BALB/C | A/ma468-G1-2/2014(H5N8) | 1 | | | PFU | HIGH | Virulent |
| Choi *et al.*, 2017 [9] | C57BL/6 | A/ma468-G2-1/2014(H5N8) | | | 4 | PFU | HIGH | Virulent |
| Choi *et al.*, 2017 [9] | BALB/C | A/ma468-G2-1/2014(H5N8) | 0.7 | | | PFU | HIGH | Virulent |
| Choi *et al.*, 2017 [9] | C57BL/6 | A/ma468-G2-2/2014(H5N8) | | | 4 | PFU | HIGH | Virulent |
| Choi *et al.*, 2017 [9] | BALB/C | A/ma468-G2-2/2014(H5N8) | 1.5 | | | PFU | HIGH | Virulent |
| Choi *et al.*, 2017 [9] | C57BL/6 | A/ma468-G2-3/2014(H5N8) | | 4 | | PFU | LOW | Avirulent |
| Choi *et al.*, 2017 [9] | BALB/C | A/ma468-G2-3/2014(H5N8) | 4.8 | | | PFU | INTERMEDIATE | Virulent |
| Choi *et al.*, 2017 [9] | C57BL/6 | A/ma468-G4-2/2014(H5N8) | | | 4 | PFU | HIGH | Virulent |
| Choi *et al.*, 2017 [9] | BALB/C | A/ma468-G4-2/2014(H5N8) | 0.5 | | | PFU | HIGH | Virulent |
| Choi *et al.*, 2017 [9] | C57BL/6 | A/mallard/Korea/W452/2014(H5N8) | | 4 | | PFU | LOW | Avirulent |
| Choi *et al.*, 2017 [9] | BALB/C | A/mallard/Korea/W452/2014(H5N8) | 7.5 | | | PFU | LOW | Avirulent |
| Manicassamy *et al.*, 2010 [44] | C57BL/6 | A/Netherlands/602/2009(H1N1) | 4.2 | | | PFU | INTERMEDIATE | Virulent |
| Pica *et al.*, 2011 [54] | DBA/2 | A/Netherlands/602/2009(H1N1) | 0.5 | | | PFU | HIGH | Virulent |
| Ye *et al.*, 2010 [75] | BALB/C | A/Netherlands/602/2009(H1N1) | 6 | | | TCID50 | INTERMEDIATE | Virulent |
| Pica *et al.*, 2011 [54] | C57BL/6 | A/NewCaledonia/20/1999(H1N1) | 5.9 | | | PFU | INTERMEDIATE | Virulent |
| Pica *et al.*, 2011 [54] | DBA/2 | A/NewCaledonia/20/1999(H1N1) | 4.3 | | | PFU | INTERMEDIATE | Virulent |
| Ping *et al.*, 2018 [56] | BALB/C | A/NewCaledonia/20/1999(H1N1) | 5.68 | | | PFU | INTERMEDIATE | Virulent |
| Jang *et al.*, 2014 [28] | BALB/C | A/Panama/2007/1999(H3N2) | | 6 | | PFU | LOW | Avirulent |
| Pica *et al.*, 2011 [54] | C57BL/6 | A/Panama/2007/1999(H3N2) | | 6 | | PFU | LOW | Avirulent |
| Pica *et al.*, 2011 [54] | DBA/2 | A/Panama/2007/1999(H3N2) | | 6 | | PFU | LOW | Avirulent |
| Blazejewska *et al.*, 2011 [5] | C57BL/6 | A/PR8F/1934(H1N1) | | | 3.3 | PFU | HIGH | Virulent |
| Blazejewska *et al.*, 2011 [5] | DBA/2 | A/PR8F/1934(H1N1) | | | 3.3 | PFU | HIGH | Virulent |
| Blazejewska *et al.*, 2011 [5] | C57BL/6 | A/PR8M/1934(H1N1) | **3.3** | 3.3 | | PFU | INTERMEDIATE | Virulent |

| | | | | | | | | |
|---|---|---|---|---|---|---|---|---|
| Blazejewska *et al.*, 2011 [5] | DBA/2 | A/PR8M/1934(H1N1) | | | 3.3 | PFU | HIGH | Virulent |
| Pica *et al.*, 2011 [54] | C57BL/6 | A/PuertoRico/8/1934(H1N1) | 1.4 | | | PFU | HIGH | Virulent |
| Pica *et al.*, 2011 [54] | DBA/2 | A/PuertoRico/8/1934(H1N1) | 0.4 | | | PFU | HIGH | Virulent |
| Ping *et al.*, 2018 [56] | BALB/C | A/PuertoRico/8/1934(H1N1) | 1.74 | | | PFU | HIGH | Virulent |
| Srivastava *et al.*, 2009 [66] | A/J | A/PuertoRico/8/1934(H1N1) | | | 3.3 | FFU | HIGH | Virulent |
| Srivastava *et al.*, 2009 [66] | CBA/J | A/PuertoRico/8/1934(H1N1) | | 3.3 | | FFU | INTERMEDIATE | Virulent |
| Srivastava *et al.*, 2009 [66] | SJL/JOrlCrl | A/PuertoRico/8/1934(H1N1) | | 3.3 | | FFU | LOW | Avirulent |
| Zhou *et al.*, 2016 [81] | C3H | A/PuertoRico/8/1934(H1N1) | | | 6 | PFU | HIGH | Virulent |
| Zhou *et al.*, 2016 [81] | 129S1/SvPasCrlVr | A/PuertoRico/8/1934(H1N1) | | | 6 | PFU | HIGH | Virulent |
| Zhou *et al.*, 2016 [81] | FVB/NJ | A/PuertoRico/8/1934(H1N1) | | 6 | | PFU | LOW | Avirulent |
| Gabriel *et al.*, 2005 [18] | BALB/C | A/seal/Massachussetts/1-SC35M/1980(H7N7) | 2.8 | | | PFU | HIGH | Virulent |
| Nurnberger *et al.*, 2016 [49] | C57BL/6 | A/seal/Massachussetts/1-SC35M/1980(H7N7) | | | 3 | PFU | HIGH | Virulent |
| Srivastava *et al.*, 2009 [66] | DBA/2 | A/seal/Massachussetts/1-SC35M/1980(H7N7) | | | 3.3 | FFU | HIGH | Virulent |
| Zhu *et al.*, 2013 [82] | C57BL/6 | A/Shanghai/4664T/2013(H7N9) | | | 5.6 | TCID50 | INTERMEDIATE | Virulent |
| Zhu *et al.*, 2013 [82] | BALB/C | A/Shanghai/4664T/2013(H7N9) | | | 5.6 | TCID50 | HIGH | Virulent |
| Zhu *et al.*, 2013 [82] | ICR | A/Shanghai/4664T/2013(H7N9) | | | 5.6 | TCID50 | INTERMEDIATE | Virulent |
| Otte *et al.*, 2011 [51] | BALB/C | A/SolomonIslands/3/2006(H1N1) | | 6 | | PFU | LOW | Avirulent |
| Otte *et al.*, 2011 [51] | C57BL/6 | A/SolomonIslands/3/2006(H1N1) | | 6 | | PFU | LOW | Avirulent |
| Pica *et al.*, 2011 [54] | DBA/2 | A/SolomonIslands/3/2006(H1N1) | 3.9 | | | PFU | INTERMEDIATE | Virulent |
| Pica *et al.*, 2011 [54] | C57BL/6 | A/swine/Kansas/77778/2007(H1N1) | 2.5 | | | PFU | HIGH | Virulent |
| Pica *et al.*, 2011 [54] | DBA/2 | A/swine/Kansas/77778/2007(H1N1) | 1 | | | PFU | HIGH | Virulent |
| Pica *et al.*, 2011 [54] | C57BL/6 | A/swine/Spain/40564/2002(H1N2) | 4.7 | | | PFU | INTERMEDIATE | Virulent |
| Pica *et al.*, 2011 [54] | DBA/2 | A/swine/Spain/40564/2002(H1N2) | 1.3 | | | PFU | HIGH | Virulent |
| Pica *et al.*, 2011 [54] | C57BL/6 | A/swine/Spain/53207/2004(H1N1) | 5.7 | | | PFU | INTERMEDIATE | Virulent |
| Pica *et al.*, 2011 [54] | DBA/2 | A/swine/Spain/53207/2004(H1N1) | 2.2 | | | PFU | HIGH | Virulent |
| Pica *et al.*, 2011 [54] | C57BL/6 | A/swine/Spain/54008/2004(H3N2) | | 6 | | PFU | LOW | Avirulent |
| Pica *et al.*, 2011 [54] | DBA/2 | A/swine/Spain/54008/2004(H3N2) | 5.5 | | | PFU | INTERMEDIATE | Virulent |
| Pica *et al.*, 2011 [54] | C57BL/6 | A/swine/Texas/4199-2/1998(H3N2) | 6.2 | | | PFU | LOW | Avirulent |
| Pica *et al.*, 2011 [54] | DBA/2 | A/swine/Texas/4199-2/1998(H3N2) | 5.7 | | | PFU | INTERMEDIATE | Virulent |
| Otte *et al.*, 2011 [51] | BALB/C | A/Thailand/KAN-1/2004(H5N1) | 0.3 | | | PFU | HIGH | Virulent |
| Otte *et al.*, 2011 [51] | C57BL/6 | A/Thailand/KAN-1/2004(H5N1) | 1.8 | | | PFU | HIGH | Virulent |
| Pica *et al.*, 2011 [54] | C57BL/6 | A/Wisconsin/67/2005(H3N2) | | 6 | | PFU | LOW | Avirulent |
| Pica *et al.*, 2011 [54] | DBA/2 | A/Wisconsin/67/2005(H3N2) | | 6 | | PFU | LOW | Avirulent |

| | | | | | | | | |
|---|---|---|---|---|---|---|---|---|
| Huo *et al.*, 2018 [22] | C57BL/6 | A/WSN/1933(H1N1) | | | 2.3 | PFU | HIGH | Virulent |
| Quan *et al.*, 2008 [60] | BALB/C | A/WSN/1933(H1N1) | 2.3 | | | PFU | HIGH | Virulent |
| Hatesuer *et al.*, 2013 [20] | C57BL/6 | maA/HongKong/1/1968(H3N2) | **1** | 1 | | FFU | HIGH | Virulent |
| Leist *et al.*, 2016 [37] | A/J | maA/HongKong/1/1968(H3N2) | | | 1 | FFU | HIGH | Virulent |
| Leist *et al.*, 2016 [37] | 129S1/SvImJ | maA/HongKong/1/1968(H3N2) | | | 1 | FFU | HIGH | Virulent |
| Leist *et al.*, 2016 [37] | NOD/ShiLtJ | maA/HongKong/1/1968(H3N2) | 1.38 | | | FFU | HIGH | Virulent |
| Leist *et al.*, 2016 [37] | NZO/HILtJ | maA/HongKong/1/1968(H3N2) | | 5 | | FFU | LOW | Avirulent |
| Leist *et al.*, 2016 [37] | CAST/EiJ | maA/HongKong/1/1968(H3N2) | | | 1 | FFU | HIGH | Virulent |
| Leist *et al.*, 2016 [37] | PWK/PhJ | maA/HongKong/1/1968(H3N2) | | 5 | | FFU | LOW | Avirulent |
| Leist *et al.*, 2016 [37] | WSB/EiJ | maA/HongKong/1/1968(H3N2) | | | 1 | FFU | HIGH | Virulent |
| Pica *et al.*, 2011 [54] | C57BL/6 | rA/PR8(123578)/ALB76(46)(H1N1) | 3.5 | | | PFU | INTERMEDIATE | Virulent |
| Pica *et al.*, 2011 [54] | DBA/2 | rA/PR8(123578)/ALB76(46)(H1N1) | 1.2 | | | PFU | HIGH | Virulent |
| Pica *et al.*, 2011 [54] | C57BL/6 | rA/PR8(123578)/UKR63(46)(H3N8) | | 6 | | PFU | LOW | Avirulent |
| Pica *et al.*, 2011 [54] | DBA/2 | rA/PR8(123578)/UKR63(46)(H3N8) | 5.5 | | | PFU | INTERMEDIATE | Virulent |
| Lu *et al.*, 1999 [41] | BALB/C | rA/X-31(H3N2) | | 5.2 | | EID50 | LOW | Avirulent |
| Pica *et al.*, 2011 [54] | C57BL/6 | rA/X-31(H3N2) | 5.3 | | | PFU | INTERMEDIATE | Virulent |
| Pica *et al.*, 2011 [54] | DBA/2 | rA/X-31(H3N2) | 0.7 | | | PFU | HIGH | Virulent |

**Table S4.** Extrapolated incomplete IAV genomes.

| No. | Extrapolated incomplete genomes | | | | | | | | |
|---|---|---|---|---|---|---|---|---|---|
| | Query | | | | | Selected BLAST hit | | | |
| | Genome ID | Sequence ID | Virus name | Segment | Notes | Sequence ID | Virus name | Query cover | Identity |
| 1 | ASSEM0044 | | A/Kawasaki/173/2001(H1N1) | 1 | BLAST the HA and NA | CY003031 | A/New York/341/2001(H1N1) | | |
| | ASSEM0044 | | A/Kawasaki/173/2001(H1N1) | 2 | BLAST the HA and NA | CY003030 | A/New York/341/2001(H1N1) | | |
| | ASSEM0044 | | A/Kawasaki/173/2001(H1N1) | 3 | BLAST the HA and NA | CY003029 | A/New York/341/2001(H1N1) | | |
| | ASSEM0044 | | A/Kawasaki/173/2001(H1N1) | 5 | BLAST the HA and NA | CY003027 | A/New York/341/2001(H1N1) | | |
| | ASSEM0044 | | A/Kawasaki/173/2001(H1N1) | 7 | BLAST the HA and NA | CY003025 | A/New York/341/2001(H1N1) | | |
| | ASSEM0044 | | A/Kawasaki/173/2001(H1N1) | 8 | BLAST the HA and NA | CY003028 | A/New York/341/2001(H1N1) | | |
| 2 | ASSEM0045 | | A/Kawasaki/UTK-4/2009(H1N1) | 1 | BLAST the HA and NA | CY043494 | A/Niigata/08F031/2009(H1N1) | | |
| | ASSEM0045 | | A/Kawasaki/UTK-4/2009(H1N1) | 2 | BLAST the HA and NA | CY043495 | A/Niigata/08F031/2009(H1N1) | | |
| | ASSEM0045 | | A/Kawasaki/UTK-4/2009(H1N1) | 3 | BLAST the HA and NA | CY043496 | A/Niigata/08F031/2009(H1N1) | | |
| | ASSEM0045 | | A/Kawasaki/UTK-4/2009(H1N1) | 5 | BLAST the HA and NA | CY043498 | A/Niigata/08F031/2009(H1N1) | | |
| | ASSEM0045 | | A/Kawasaki/UTK-4/2009(H1N1) | 7 | BLAST the HA and NA | CY043500 | A/Niigata/08F031/2009(H1N1) | | |
| | ASSEM0045 | | A/Kawasaki/UTK-4/2009(H1N1) | 8 | BLAST the HA and NA | CY043501 | A/Niigata/08F031/2009(H1N1) | | |
| 3 | ASSEM0046 | | A/VietNam/1204/2004(H5N1) | 5 | BLAST the HA and NA | HM006760 | A/Viet Nam/1203/2004(H5N1) | | |
| | ASSEM0046 | | A/VietNam/1204/2004(H5N1) | 7 | BLAST the HA and NA | HM006762 | A/Viet Nam/1203/2004(H5N1) | | |
| 4 | ASSEM0047 | | A/SolomonIslands/3/2006(H1N1) | 1 | BLAST the HA and NA | CY119233 | A/Malaysia/1706215/2007(H1N1) | | |
| | ASSEM0047 | | A/SolomonIslands/3/2006(H1N1) | 2 | BLAST the HA and NA | CY119232 | A/Malaysia/1706215/2007(H1N1) | | |
| | ASSEM0047 | | A/SolomonIslands/3/2006(H1N1) | 8 | BLAST the HA and NA | CY119230 | A/Malaysia/1706215/2007(H1N1) | | |
| 5 | ASSEM0048 | | A/Tennessee/1-560/2009(H1N1) | 5 | BLAST the HA and NA | KY926173 | A/Sao Gabriel/LACENRS-1626/2009(H1N1) | | |
| | ASSEM0048 | | A/Tennessee/1-560/2009(H1N1) | 7 | BLAST the HA and NA | KY926049 | A/Sao Gabriel/LACENRS-1626/2009(H1N1) | | |
| | ASSEM0048 | | A/Tennessee/1-560/2009(H1N1) | 8 | BLAST the HA and NA | KY925946 | A/Sao Gabriel/LACENRS-1626/2009(H1N1) | | |
| 6 | ASSEM0052 | | A/NWS/1933(H1N1) | 1 | BLAST the HA and NA | CY120991 | A/NWS/1934(H1N1) | | |
| | ASSEM0052 | | A/NWS/1933(H1N1) | 2 | BLAST the HA and NA | CY120990 | A/NWS/1934(H1N1) | | |
| | ASSEM0052 | | A/NWS/1933(H1N1) | 3 | BLAST the HA and NA | CY120989 | A/NWS/1934(H1N1) | | |
| | ASSEM0052 | | A/NWS/1933(H1N1) | 5 | BLAST the HA and NA | CY120987 | A/NWS/1934(H1N1) | | |
| 7 | ASSEM0053 | | A/duck/Minnesota/1525/1981(H5N1) | 6 | BLAST the HA | CY179413 | A/mallard/Wisconsin/568/1982(H5N1) | | |
| 8 | ASSEM0055 | | A/LaReunion/803/2010(H1N1) | 1 | BLAST the HA and NA | JX309663 | A/Singapore/GP3667/2010(H1N1) | | |
| 9 | ASSEM0056 | | A/Mississippi/03/2001(H1N1) | 1 | BLAST the NA | CY016267 | A/New South Wales/26/2000(H1N1) | | |
| | ASSEM0056 | | A/Mississippi/03/2001(H1N1) | 2 | BLAST the NA | CY016266 | A/New South Wales/26/2000(H1N1) | | |
| | ASSEM0056 | | A/Mississippi/03/2001(H1N1) | 3 | BLAST the NA | CY016265 | A/New South Wales/26/2000(H1N1) | | |
| | ASSEM0056 | | A/Mississippi/03/2001(H1N1) | 4 | BLAST the NA | CY016260 | A/New South Wales/26/2000(H1N1) | | |
| | ASSEM0056 | | A/Mississippi/03/2001(H1N1) | 5 | BLAST the NA | CY016263 | A/New South Wales/26/2000(H1N1) | | |
| | ASSEM0056 | | A/Mississippi/03/2001(H1N1) | 7 | BLAST the NA | CY016261 | A/New South Wales/26/2000(H1N1) | | |
| | ASSEM0056 | | A/Mississippi/03/2001(H1N1) | 8 | BLAST the NA | CY016264 | A/New South Wales/26/2000(H1N1) | | |
| 10 | ASSEM0059 | | A/Turkey/13/2006(H5N1) | 1 | Matching by the closest name | EF620011 | A/Turkey/15/2006(H5N1) | | |
| | ASSEM0059 | | A/Turkey/13/2006(H5N1) | 2 | Matching by the closest name | EF620010 | A/Turkey/15/2006(H5N1) | | |
| | ASSEM0059 | | A/Turkey/13/2006(H5N1) | 3 | Matching by the closest name | EF620009 | A/Turkey/15/2006(H5N1) | | |

| | | | | | | | |
|---|---|---|---|---|---|---|---|
| | ASSEM0059 | | A/Turkey/13/2006(H5N1) | 4 | Matching by the closest name | EF619989 | A/Turkey/15/2006(H5N1) |
| | ASSEM0059 | | A/Turkey/13/2006(H5N1) | 5 | Matching by the closest name | EF620007 | A/Turkey/15/2006(H5N1) |
| | ASSEM0059 | | A/Turkey/13/2006(H5N1) | 6 | Matching by the closest name | EF619988 | A/Turkey/15/2006(H5N1) |
| | ASSEM0059 | | A/Turkey/13/2006(H5N1) | 7 | Matching by the closest name | EF620006 | A/Turkey/15/2006(H5N1) |
| | ASSEM0059 | | A/Turkey/13/2006(H5N1) | 8 | Matching by the closest name | EF620008 | A/Turkey/15/2006(H5N1) |
| 11 | ASSEM0065 | | B/Shangdong/7/1997 | 1 | BLAST the HA | AY582052 | B/Nanchang/2/1997 |
| | ASSEM0065 | | B/Shangdong/7/1997 | 2 | BLAST the HA | AY582065 | B/Nanchang/2/1997 |
| | ASSEM0065 | | B/Shangdong/7/1997 | 3 | BLAST the HA | AY582039 | B/Nanchang/2/1997 |
| | ASSEM0065 | | B/Shangdong/7/1997 | 5 | BLAST the HA | AY582025 | B/Nanchang/2/1997 |
| | ASSEM0065 | | B/Shangdong/7/1997 | 6 | BLAST the HA | AY582003 | B/Nanchang/2/1997 |
| | ASSEM0065 | | B/Shangdong/7/1997 | 7 | BLAST the HA | AY582077 | B/Nanchang/2/1997 |
| | ASSEM0065 | | B/Shangdong/7/1997 | 8 | BLAST the HA | AY582078 | B/Nanchang/2/1997 |
| 12 | ASSEM0086 | | A/Netherlands/230/2003(H7N7) | 2 | BLAST the HA and NA | AB438939 | A/chicken/Netherlands/2586/2003(H7N7) |
| | ASSEM0086 | | A/Netherlands/230/2003(H7N7) | 3 | BLAST the HA and NA | AB438940 | A/chicken/Netherlands/2586/2003(H7N7) |
| | ASSEM0086 | | A/Netherlands/230/2003(H7N7) | 5 | BLAST the HA and NA | AB438942 | A/chicken/Netherlands/2586/2003(H7N7) |
| | ASSEM0086 | | A/Netherlands/230/2003(H7N7) | 7 | BLAST the HA and NA | AB438944 | A/chicken/Netherlands/2586/2003(H7N7) |
| | ASSEM0086 | | A/Netherlands/230/2003(H7N7) | 8 | BLAST the HA and NA | AB438945 | A/chicken/Netherlands/2586/2003(H7N7) |
| 13 | ASSEM0088 | | A/Netherlands/603/2009(H1N1) | 3 | BLAST the HA | MG856195 | A/Mexico/IBT25/2009(H1N1) |
| | ASSEM0088 | | A/Netherlands/603/2009(H1N1) | 6 | BLAST the HA | MG856198 | A/Mexico/IBT25/2009(H1N1) |
| | ASSEM0088 | | A/Netherlands/603/2009(H1N1) | 7 | BLAST the HA | MG856199 | A/Mexico/IBT25/2009(H1N1) |
| | ASSEM0088 | | A/Netherlands/603/2009(H1N1) | 8 | BLAST the HA | MG856200 | A/Mexico/IBT25/2009(H1N1) |
| 14 | ASSEM0090 | | A/Seoul/Y-01/2009(H1N1) | 1 | BLAST the HA and NA | CY128218 | A/Viet Nam/13032036/2009(H1N1) |
| | ASSEM0090 | | A/Seoul/Y-01/2009(H1N1) | 2 | BLAST the HA and NA | CY128217 | A/Viet Nam/13032036/2009(H1N1) |
| | ASSEM0090 | | A/Seoul/Y-01/2009(H1N1) | 3 | BLAST the HA and NA | CY128216 | A/Viet Nam/13032036/2009(H1N1) |
| | ASSEM0090 | | A/Seoul/Y-01/2009(H1N1) | 5 | BLAST the HA and NA | CY128214 | A/Viet Nam/13032036/2009(H1N1) |
| | ASSEM0090 | | A/Seoul/Y-01/2009(H1N1) | 7 | BLAST the HA and NA | CY128212 | A/Viet Nam/13032036/2009(H1N1) |
| | ASSEM0090 | | A/Seoul/Y-01/2009(H1N1) | 8 | BLAST the HA and NA | CY128215 | A/Viet Nam/13032036/2009(H1N1) |

**Table S5.** Extrapolated partial IAV nucleotide sequences.

| No. | Extrapolated partial sequences | | | | | | | | |
|---|---|---|---|---|---|---|---|---|---|
| | Query | | | | | BLAST hit | | | |
| | Genome ID | Sequence ID | Virus name | Segment | Notes | Sequence ID | Virus name | Query cover | Identity |
| 1 | ASSEM0001 | AF116575 | A/BrevigMission/1/1918(H1N1) | 4 | | AF117241 | A/South Carolina/1/18 (H1N1) | 100% | 99% |
| 2 | ASSEM0002 | KC853225 | A/Shanghai/4664T/2013(H7N9) | 5 | | MF988739 | A/duck/Anhui/S702/2013(H7N9) | 100% | 100% |
| 3 | ASSEM0003 | AF036361 | A/HongKong/156/1997(H5N1) | 3 | | AJ289874 | A/Hong Kong/156/97(H5N1) | 100% | 99% |
| | ASSEM0003 | AF036356 | A/HongKong/156/1997(H5N1) | 4 | | GU052127 | A/environment/Hong Kong/156/1997(H5N1) | 100% | 99% |
| 4 | ASSEM0004 | AF046093 | A/HongKong/156/1997(H5N1) | 1 | | GU052134 | A/environment/Hong Kong/156/1997(H5N1) | 100% | 100% |
| 5 | ASSEM0007 | HQ111367 | A/Hamburg/05/2009(H1N1) | 7 | | CY266344 | A/California/07-00019/2009(H1N1) | 100% | 100% |
| | ASSEM0007 | HQ111368 | A/Hamburg/05/2009(H1N1) | 8 | | CY266347 | A/California/07-00019/2009(H1N1) | 100% | 100% |
| 6 | ASSEM0008 | HQ104928 | A/Hamburg/NY1580/2009(H1N1) | 7 | | KP412332 | A/swine/Illinois/21-1230/2009(H1N1) | 100% | 100% |
| | ASSEM0008 | HQ104929 | A/Hamburg/NY1580/2009(H1N1) | 8 | | JX625532 | A/Northern Ireland/94480397/2009(H1N1) | 100% | 100% |
| 7 | ASSEM0014 | ID0007 | A/ma452-G1-1/2014(H5N8) | 7 | | MG965867 | A/turkey/Wisconsin/15-014298-1/2015(H5N2) | 100% | 100% |
| | ASSEM0014 | ID0008 | A/ma452-G1-1/2014(H5N8) | 8 | | MG964812 | A/red-tailed hawk/Washington/15-002551-2/2015(H5N2) | 100% | 100% |
| 8 | ASSEM0015 | ID0015 | A/ma452-G3-1/2014(H5N8) | 7 | | MG965867 | A/turkey/Wisconsin/15-014298-1/2015(H5N2) | 100% | 100% |
| | ASSEM0015 | ID0016 | A/ma452-G3-1/2014(H5N8) | 8 | | MG964812 | A/red-tailed hawk/Washington/15-002551-2/2015(H5N2) | 100% | 100% |
| 9 | ASSEM0016 | ID0023 | A/ma452-G3-2/2014(H5N8) | 7 | | MG965867 | A/turkey/Wisconsin/15-014298-1/2015(H5N2) | 100% | 100% |
| | ASSEM0016 | ID0024 | A/ma452-G3-2/2014(H5N8) | 8 | | MG964812 | A/red-tailed hawk/Washington/15-002551-2/2015(H5N2) | 100% | 100% |
| 10 | ASSEM0017 | ID0031 | A/ma452-G4-1/2014(H5N8) | 7 | | MG965867 | A/turkey/Wisconsin/15-014298-1/2015(H5N2) | 100% | 100% |
| | ASSEM0017 | ID0032 | A/ma452-G4-1/2014(H5N8) | 8 | | MG964812 | A/red-tailed hawk/Washington/15-002551-2/2015(H5N2) | 100% | 100% |
| 11 | ASSEM0018 | ID0039 | A/ma468-G1-1/2014(H5N8) | 7 | | KX297904 | A/environment/Korea/W477/2014(H5N8) | 100% | 100% |
| | ASSEM0018 | ID0040 | A/ma468-G1-1/2014(H5N8) | 8 | | KX297970 | A/environment/Korea/W468/2014(H5N8) | 100% | 100% |
| 12 | ASSEM0019 | ID0047 | A/ma468-G1-2/2014(H5N8) | 7 | | KX297904 | A/environment/Korea/W477/2014(H5N8) | 100% | 100% |
| | ASSEM0019 | ID0048 | A/ma468-G1-2/2014(H5N8) | 8 | | KX297970 | A/environment/Korea/W468/2014(H5N8) | 100% | 100% |
| 13 | ASSEM0020 | ID0055 | A/ma468-G2-1/2014(H5N8) | 7 | | KX297904 | A/environment/Korea/W477/2014(H5N8) | 100% | 100% |
| | ASSEM0020 | ID0056 | A/ma468-G2-1/2014(H5N8) | 8 | | KX297970 | A/environment/Korea/W468/2014(H5N8) | 100% | 100% |
| 14 | ASSEM0021 | ID0063 | A/ma468-G2-2/2014(H5N8) | 7 | | KX297904 | A/environment/Korea/W477/2014(H5N8) | 100% | 100% |
| | ASSEM0021 | ID0064 | A/ma468-G2-2/2014(H5N8) | 8 | | KX297970 | A/environment/Korea/W468/2014(H5N8) | 100% | 100% |
| 15 | ASSEM0022 | ID0071 | A/ma468-G2-3/2014(H5N8) | 7 | | KX297904 | A/environment/Korea/W477/2014(H5N8) | 100% | 100% |
| | ASSEM0022 | ID0072 | A/ma468-G2-3/2014(H5N8) | 8 | | KX297970 | A/environment/Korea/W468/2014(H5N8) | 100% | 100% |
| 16 | ASSEM0023 | ID0076 | A/ma468-G4-2/2014(H5N8) | 4 | | KX297879 | A/environment/Korea/W477/2014(H5N8) | 100% | 99% |
| | ASSEM0023 | ID0079 | A/ma468-G4-2/2014(H5N8) | 7 | | KX297904 | A/environment/Korea/W477/2014(H5N8) | 100% | 100% |
| | ASSEM0023 | ID0080 | A/ma468-G4-2/2014(H5N8) | 8 | | KX297970 | A/environment/Korea/W468/2014(H5N8) | 100% | 100% |
| 17 | ASSEM0024 | GQ495134 | A/swine/Sweden/1021/2009(H1N2) | 6 | | KR066629 | A/swine/Denmark/10832-1/2009(H1N2) | 100% | 99% |
| | ASSEM0024 | GQ495135 | A/swine/Sweden/1021/2009(H1N2) | 7 | | KR700052 | A/swine/Denmark/10-1725-1/2011(H1N2) | 100% | 98% |
| 18 | ASSEM0025 | HM626484 | A/swine/Sweden/9706/2010(H1N2) | 6 | | KR066629 | A/swine/Denmark/10832-1/2009(H1N2) | 100% | 98% |
| | ASSEM0025 | HM626485 | A/swine/Sweden/9706/2010(H1N2) | 7 | | KR700052 | A/swine/Denmark/10-1725-1/2011(H1N2) | 100% | 98% |
| 19 | ASSEM0027 | GU052022 | A/goose/HongKong/437-6/1999(H5N1) | 5 | | AF216720 | A/environment/Hong Kong/437-6/99 (H5N1) | 99% | 99% |
| 20 | ASSEM0033 | AF084280 | A/HongKong/483/1997(H5N1) | 4 | | AF046097 | A/Hong Kong/483/97(H5N1) | 100% | 99% |

| | | | | | | | | | |
|---|---|---|---|---|---|---|---|---|---|
| 21 | ASSEM0035 | GU052103 | A/HongKong/483/1997(H5N1) | 2 | | GU052182 | A/environment/Hong Kong/258/1997(H5) | 100% | 99% |
| | ASSEM0035 | GU052100 | A/HongKong/483/1997(H5N1) | 5 | | AF255746 | A/Hong Kong/483/97(H5N1) | 100% | 100% |
| 22 | ASSEM0036 | GU052143 | A/HongKong/485/1997(H5N1) | 7 | | AF046082 | A/Chicken/Hong Kong/220/97 (H5N1) | 100% | 99% |
| 23 | ASSEM0037 | AF084532 | A/HongKong/485/1997(H5N1) | 4 | | GU052142 | A/Hong Kong/485/1997(H5N1) | 100% | 100% |
| 24 | ASSEM0038 | AF084281 | A/HongKong/486/1997(H5N1) | 4 | | AF046098 | A/Hong Kong/482/97(H5N1) | 100% | 100% |
| 25 | ASSEM0039 | AF102671 | A/HongKong/486/1997(H5N1) | 4 | | AF046098 | A/Hong Kong/482/97(H5N1) | 100% | 100% |
| 26 | ASSEM0040 | EF473503 | A/Thailand/16/2004(H5N1) | 3 | | EU268218 | A/Thailand/16/2004(H5N1) | 100% | 99% |
| 27 | ASSEM0046 | EF473407 | A/VietNam/1204/2004(H5N1) | 3 | | HM006758 | A/Viet Nam/1203/2004(H5N1) | 100% | 99% |
| 28 | ASSEM0047 | CY047425 | A/SolomonIslands/3/2006(H1N1) | 3 | | CY119231 | A/Malaysia/1706215/2007(H1N1) | | |
| | ASSEM0047 | EU124136 | A/SolomonIslands/3/2006(H1N1) | 6 | | CY119228 | A/Malaysia/1706215/2007(H1N1) | | |
| 29 | ASSEM0048 | CY040457 | A/Tennessee/1-560/2009(H1N1) | 4 | | KY926119 | A/Teutonia/LACENRS-711/2009(H1N1) | 100% | 99% |
| | ASSEM0048 | CY040458 | A/Tennessee/1-560/2009(H1N1) | 6 | | KY925637 | A/Porto Alegre/LACENRS-1786/2009(H1N1) | 100% | 100% |
| 30 | ASSEM0049 | GU361156 | A/aquaticbird/Korea/w81/2005(H5N2) | 5 | | DQ323677 | A/chicken/Kurgan/3/2005(H5N1) | 100% | 99% |
| 31 | ASSEM0058 | KF897796 | A/StEtienne/1139/2010(H1N1) | 7 | | JX625745 | A/England/182/2010(H1N1) | 100% | 100% |
| 32 | ASSEM0059 | EF620011 | A/Turkey/13/2006(H5N1) | 1 | including insertion at position 2079 | EU146846 | A/Iraq/1/2006(H5N1) | 100% | 99% |
| | ASSEM0059 | EF620009 | A/Turkey/13/2006(H5N1) | 3 | | EF619995 | A/Turkey/651242/2006(H5N1) | 100% | 100% |
| | ASSEM0059 | EF619989 | A/Turkey/13/2006(H5N1) | 4 | | EF446779 | A/goose/Hungary/3413/2007(H5N1) | 100% | 99% |
| | ASSEM0059 | EF620007 | A/Turkey/13/2006(H5N1) | 5 | | DQ323677 | A/chicken/Kurgan/3/2005(H5N1) | 100% | 99% |
| | ASSEM0059 | EF619988 | A/Turkey/13/2006(H5N1) | 6 | | EF620000 | A/Turkey/65596/2006(H5N1) | 100% | 100% |
| | ASSEM0059 | EF620006 | A/Turkey/13/2006(H5N1) | 7 | | DQ234077 | A/grebe/Novosibirsk/29/2005(H5N1) | 100% | 99% |
| | ASSEM0059 | EF620008 | A/Turkey/13/2006(H5N1) | 8 | | EU599286 | A/chicken/Iraq/900845/2006(H5N1) | 100% | 99% |
| 33 | ASSEM0064 | KF897804 | A/Limoges/1159/2010(H1N1) | 7 | | JX625745 | A/England/182/2010(H1N1) | 100% | 100% |
| 34 | ASSEM0065 | EF541422 | B/Shangdong/7/1997 | 4 | | AY581956 | B/Nanchang/2/1997 | 100% | 99% |
| 35 | ASSEM0066 | KF897812 | A/Lyon/1.12/2011(H1N1) | 7 | | KY925952 | A/Viamao/LACENRS-1400/2011(H1N1) | 100% | 100% |
| 36 | ASSEM0067 | KF897788 | A/Lyon/52.16/2010(H1N1) | 7 | | KC488882 | A/Chita/RII08/2012(H1N1) | 100% | 100% |
| | ASSEM0067 | GQ166219 | A/Hamburg/4/2009(H1N1) | 7 | | KX571118 | A/swine/Quebec/1279825/2011(H3N2) | 100% | 100% |
| 37 | ASSEM0069 | AF102663 | A/HongKong/481/1997(H5N1) | 6 | | AF084271 | A/HongKong/481/1997(H5N1) | 100% | 99% |
| 38 | ASSEM0070 | AF084279 | A/HongKong/481/1997(H5N1) | 4 | | AF046096 | A/HongKong/481/1997(H5N1) | 100% | 99% |
| 39 | ASSEM0071 | AF258847 | A/HongKong/485/1997(H5N1) | 1 | | GU052149 | A/Hong Kong/485/1997(H5N1) | 100% | 99% |
| | ASSEM0071 | AF258828 | A/HongKong/485/1997(H5N1) | 2 | | GU052182 | A/environment/Hong Kong/258/1997(H5) | 100% | 99% |
| | ASSEM0071 | AF257203 | A/HongKong/485/1997(H5N1) | 3 | | AF046087 | A/Chicken/Hong Kong/220/97(H5N1) | 100% | 99% |
| | ASSEM0071 | AF102681 | A/HongKong/485/1997(H5N1) | 4 | | GU052142 | A/Hong Kong/485/1997(H5N1) | 100% | 100% |
| | ASSEM0071 | AH010699 | A/HongKong/485/1997(H5N1) | 5 | | GU052145 | A/Hong Kong/485/1997(H5N1) | 91% | 99% |
| | ASSEM0071 | AF102664 | A/HongKong/485/1997(H5N1) | 6 | | GU052144 | A/Hong Kong/485/1997(H5N1) | 100% | 99% |
| | ASSEM0071 | AH010696 | A/HongKong/485/1997(H5N1) | 7 | | JX465629 | A/chicken/Iran/ZMT-101/1998(H9N2) | 86% | 99% |
| | ASSEM0071 | AF256189 | A/HongKong/485/1997(H5N1) | 8 | | GU052146 | A/Hong Kong/485/1997(H5N1) | 100% | 100% |
| 40 | ASSEM0072 | AF258848 | A/HongKong/488/1997(H5N1) | 1 | | GU052041 | A/environment/Hong Kong/486/1997(H5N1) | 100% | 99% |
| | ASSEM0072 | AF258829 | A/HongKong/488/1997(H5N1) | 2 | | GU052040 | A/environment/Hong Kong/486/1997(H5N1) | 100% | 99% |
| | ASSEM0072 | AF257204 | A/HongKong/488/1997(H5N1) | 3 | | AF046087 | A/Chicken/Hong Kong/220/97(H5N1) | 100% | 99% |
| | ASSEM0072 | AF102672 | A/HongKong/488/1997(H5N1) | 4 | | AF046098 | A/Hong Kong/482/97(H5N1) | 100% | 99% |
| | ASSEM0072 | AH010700 | A/HongKong/488/1997(H5N1) | 5 | | GU052037 | A/environment/Hong Kong/486/1997(H5N1) | 90% | 100% |
| | ASSEM0072 | AF102657 | A/HongKong/488/1997(H5N1) | 6 | | GU052036 | A/environment/Hong Kong/486/1997(H5N1) | 100% | 99% |
| | ASSEM0072 | AH010697 | A/HongKong/488/1997(H5N1) | 7 | Replace Ns at the centre | AF255373 | A/Hong Kong/542/97(H5N1) | 86% | 99% |
| | ASSEM0072 | AF256190 | A/HongKong/488/1997(H5N1) | 8 | | AF084285 | A/HongKong/482/97(H5N1) | 100% | 100% |
| 41 | ASSEM0073 | AF258849 | A/HongKong/491/1997(H5N1) | 1 | | AF258845 | A/Hong Kong/542/97(H5N1) | 100% | 99% |

| | | | | | | | | | |
|---|---|---|---|---|---|---|---|---|---|
| | ASSEM0073 | AF258830 | A/HongKong/491/1997(H5N1) | 2 | | AF258826 | A/Hong Kong/542/97(H5N1) | 100% | 100% |
| | ASSEM0073 | AF257205 | A/HongKong/491/1997(H5N1) | 3 | | AF046087 | A/Chicken/Hong Kong/220/97 (H5N1) | 100% | 99% |
| | ASSEM0073 | AF102677 | A/HongKong/491/1997(H5N1) | 4 | | AF082034 | A/Chicken/Hong Kong/728/97 (H5N1) | 100% | 99% |
| | ASSEM0073 | AH010701 | A/HongKong/491/1997(H5N1) | 5 | | AF098619 | A/Chicken/Hong Kong/786/97 (H5N1) | 90% | 99% |
| | ASSEM0073 | AF102665 | A/HongKong/491/1997(H5N1) | 6 | | GU052122 | A/chicken/China/27402/1997(H5N1) | 100% | 99% |
| | ASSEM0073 | AH010698 | A/HongKong/491/1997(H5N1) | 7 | | AF255365 | A/Hong Kong/481/97(H5N1) | 87% | 99% |
| | ASSEM0073 | AF256191 | A/HongKong/491/1997(H5N1) | 8 | | GU052138 | A/chicken/Hong Kong/786-2/1997(H5N1) | 100% | 99% |
| 42 | ASSEM0074 | AF258850 | A/HongKong/503/1997(H5N1) | 1 | | AF258845 | A/Hong Kong/542/97(H5N1) | 100% | 99% |
| | ASSEM0074 | AF258831 | A/HongKong/503/1997(H5N1) | 2 | | AF258826 | A/Hong Kong/542/97(H5N1) | 100% | 99% |
| | ASSEM0074 | AF257206 | A/HongKong/503/1997(H5N1) | 3 | | AF046087 | A/Chicken/Hong Kong/220/97(H5N1) | 100% | 100% |
| | ASSEM0074 | AF102679 | A/HongKong/503/1997(H5N1) | 4 | | AF082034 | A/Chicken/Hong Kong/728/97(H5N1) | 100% | 99% |
| | ASSEM0074 | AH010702 | A/HongKong/503/1997(H5N1) | 5 | | AF098619 | A/Chicken/Hong Kong/786/97(H5N1) | 89% | 100% |
| | ASSEM0074 | AF102666 | A/HongKong/503/1997(H5N1) | 6 | | GU052122 | A/chicken/China/27402/1997(H5N1) | 100% | 99% |
| | ASSEM0074 | AF255381 | A/HongKong/503/1997(H5N1) | 7 | | AF255373 | A/Hong Kong/542/97(H5N1) | 100% | 99% |
| | ASSEM0074 | AF256192 | A/HongKong/503/1997(H5N1) | 8 | | GU052138 | A/chicken/Hong Kong/786-2/1997(H5N1) | 100% | 98% |
| 43 | ASSEM0075 | AF258851 | A/HongKong/507/1997(H5N1) | 1 | | GU052041 | A/environment/Hong Kong/486/1997(H5N1) | 100% | 100% |
| | ASSEM0075 | AF258832 | A/HongKong/507/1997(H5N1) | 2 | | GU052040 | A/environment/Hong Kong/486/1997(H5N1) | 100% | 99% |
| | ASSEM0075 | AF257207 | A/HongKong/507/1997(H5N1) | 3 | | AF046087 | A/Chicken/Hong Kong/220/97(H5N1) | 100% | 99% |
| | ASSEM0075 | AF102675 | A/HongKong/507/1997(H5N1) | 4 | | AF046098 | A/Hong Kong/482/97(H5N1) | 100% | 99% |
| | ASSEM0075 | AH010703 | A/HongKong/507/1997(H5N1) | 5 | | GU052037 | A/environment/Hong Kong/486/1997(H5N1) | 85% | 100% |
| | ASSEM0075 | AF102659 | A/HongKong/507/1997(H5N1) | 6 | | GU052036 | A/environment/Hong Kong/486/1997(H5N1) | 100% | 99% |
| | ASSEM0075 | AF255382 | A/HongKong/507/1997(H5N1) | 7 | | AF255372 | A/Hong Kong/538/97(H5N1) | 100% | 100% |
| | ASSEM0075 | AF256193 | A/HongKong/507/1997(H5N1) | 8 | | GU052038 | A/environment/Hong Kong/486/1997(H5N1) | 100% | 99% |
| 44 | ASSEM0076 | AF258852 | A/HongKong/514/1997(H5N1) | 1 | | AB586776 | A/quail/Hong Kong/NT342/2001(H6N1) | 100% | 99% |
| | ASSEM0076 | AF258833 | A/HongKong/514/1997(H5N1) | 2 | | GU052182 | A/environment/Hong Kong/258/1997(H5) | 100% | 99% |
| | ASSEM0076 | AF257208 | A/HongKong/514/1997(H5N1) | 3 | | AF046087 | A/Chicken/Hong Kong/220/97(H5N1) | 100% | 99% |
| | ASSEM0076 | AF102682 | A/HongKong/514/1997(H5N1) | 4 | | GU052142 | A/Hong Kong/485/1997(H5N1) | 100% | 99% |
| | ASSEM0076 | AH010704 | A/HongKong/514/1997(H5N1) | 5 | | AF046084 | A/Chicken/Hong Kong/220/97 (H5N1) | 90% | 99% |
| | ASSEM0076 | AF102669 | A/HongKong/514/1997(H5N1) | 6 | | GU052122 | A/chicken/China/27402/1997(H5N1) | 100% | 99% |
| | ASSEM0076 | AF255383 | A/HongKong/514/1997(H5N1) | 7 | | KY785902 | A/quail/Hong Kong/G1/1997(H9N2) | 100% | 100% |
| 45 | ASSEM0077 | AF258853 | A/HongKong/516/1997(H5N1) | 1 | | GU052041 | A/environment/Hong Kong/486/1997(H5N1) | 100% | 99% |
| | ASSEM0077 | AF258834 | A/HongKong/516/1997(H5N1) | 2 | | GU052040 | A/environment/Hong Kong/486/1997(H5N1) | 100% | 100% |
| | ASSEM0077 | AF257209 | A/HongKong/516/1997(H5N1) | 3 | | GU052039 | A/environment/Hong Kong/486/1997(H5N1) | 100% | 100% |
| | ASSEM0077 | AF102673 | A/HongKong/516/1997(H5N1) | 4 | | AF046098 | A/Hong Kong/482/97(H5N1) | 100% | 99% |
| | ASSEM0077 | AH010705 | A/HongKong/516/1997(H5N1) | 5 | | GU052037 | A/environment/Hong Kong/486/1997(H5N1) | 89% | 100% |
| | ASSEM0077 | AF102660 | A/HongKong/516/1997(H5N1) | 6 | | AF084272 | A/HongKong/482/97(H5N1) | 100% | 99% |
| | ASSEM0077 | AF255384 | A/HongKong/516/1997(H5N1) | 7 | | AF255372 | A/Hong Kong/538/97(H5N1) | 100% | 100% |
| | ASSEM0077 | AF256194 | A/HongKong/516/1997(H5N1) | 8 | | GU052038 | A/environment/Hong Kong/486/1997(H5N1) | 100% | 99% |
| 46 | ASSEM0078 | AF102680 | A/HongKong/532/1997(H5N1) | 4 | | GU052135 | A/chicken/Hong Kong/786-2/1997(H5N1) | 100% | 99% |
| | ASSEM0078 | AF102667 | A/HongKong/532/1997(H5N1) | 6 | | GU052122 | A/chicken/China/27402/1997(H5N1) | 100% | 99% |
| 47 | ASSEM0079 | AF102674 | A/HongKong/538/1997(H5N1) | 4 | | AF046098 | A/Hong Kong/482/97(H5N1) | 100% | 99% |
| | ASSEM0079 | AF102662 | A/HongKong/538/1997(H5N1) | 6 | | AF084272 | A/HongKong/482/97(H5N1) | 100% | 99% |
| 48 | ASSEM0080 | AF102678 | A/HongKong/542/1997(H5N1) | 4 | | AF082034 | A/Chicken/Hong Kong/728/97(H5N1) | 100% | 99% |
| | ASSEM0080 | AF102670 | A/HongKong/542/1997(H5N1) | 6 | | GU052122 | A/chicken/China/27402/1997(H5N1) | 100% | 99% |
| 49 | ASSEM0081 | AF102676 | A/HongKong/97/1998(H5N1) | 4 | | AF046098 | A/Hong Kong/482/97(H5N1) | 99% | 99% |
| | ASSEM0081 | AF102661 | A/HongKong/97/1998(H5N1) | 6 | | AF084272 | A/HongKong/482/97(H5N1) | 100% | 99% |

| | | | | | | | | | |
|---|---|---|---|---|---|---|---|---|---|
| 50 | ASSEM0085 | KP459007 | A/Lyon/1337/2007(H1N1) | 4 | | CY105086 | A/KhanhHoa/KH161/2008(H1N1) | 100% | 99% |
| 51 | ASSEM0086 | EPI319935 | A/Netherlands/230/2003(H7N7) | 1 | | EF015558 | A/chicken/Netherlands/03010132/03(H7N7) | 100% | 99% |
| | ASSEM0086 | EPI319937 | A/Netherlands/230/2003(H7N7) | 4 | | AB438941 | A/chicken/Netherlands/2586/2003(H7N7) | 100% | 99% |
| | ASSEM0086 | EPI319936 | A/Netherlands/230/2003(H7N7) | 6 | | AB438943 | A/chicken/Netherlands/2586/2003(H7N7) | 100% | 99% |
| 52 | ASSEM0087 | EU587371 | A/NewYork/107/2003(H7N2) | 5 | | CY031654 | A/guinea fowl/New York/22071/2005(H7N2) | 100% | 99% |
| 53 | ASSEM0089 | DQ492869 | A/chicken/VietNam/8/2003(H5N1) | 1 | | CY034205 | A/chicken/Vietnam/18/2004(H5N1) | 100% | 99% |
| | ASSEM0089 | DQ493393 | A/chicken/VietNam/8/2003(H5N1) | 2 | | CY028706 | A/chicken/Vietnam/10/2004(H5N1) | 100% | 99% |
| | ASSEM0089 | DQ493306 | A/chicken/VietNam/8/2003(H5N1) | 3 | Remove the first 24 nts (low quality??) | HM627920 | A/openbill stork/Thailand/VSMU-5-NSN/2004(H5N1) | 100% | 99% |
| | ASSEM0089 | DQ497693 | A/chicken/VietNam/8/2003(H5N1) | 4 | | AB440326 | A/quail/Angthong/71/2004(H5N1) | 100% | 99% |
| | ASSEM0089 | DQ493131 | A/chicken/VietNam/8/2003(H5N1) | 5 | | AY627895 | A/Thailand/2(SP-33)/2004(H5N1) | 100% | 100% |
| | ASSEM0089 | DQ492954 | A/chicken/VietNam/8/2003(H5N1) | 7 | | EF541449 | A/chicken/Viet Nam/1/2004(H5N1) | 100% | 100% |
| | ASSEM0089 | DQ493218 | A/chicken/VietNam/8/2003(H5N1) | 8 | | AY651555 | A/Viet Nam/3062/2004(H5N1) | 100% | 99% |
| 54 | EPIISL249281 | EPI919533 | A/chicken/Heyuan/16876/2016(H7N9) | 4 | | EF446779 | A/goose/Hungary/3413/2007(H5N1) | 100% | 99% |
| | EPIISL249281 | EPI919537 | A/chicken/Heyuan/16876/2016(H7N9) | 8 | | EU599286 | A/chicken/Iraq/900845/2006(H5N1) | 100% | 100% |
| 55 | EPIISL259927 | EPI1215865 | A/goose/Guangdong/SH7/2013(H5N1) | 7 | | KP732557 | A/chicken/AnNing/1/2014(H5N1) | 100% | 100% |
| 56 | EPIISL30054 | EPI291908 | A/Ohio/02/2007(H1N1) | 1 | | EU604691 | A/swine/OH/511445/2007(H1N1) | 100% | 99% |
| | EPIISL30054 | EPI338828 | A/Ohio/02/2007(H1N1) | 6 | | EU604690 | A/swine/OH/511445/2007(H1N1) | 100% | 100% |
| 57 | EPIISL3889 | EPI21222 | A/duck/Fujian/01/2002(H5N1) | 8 | | HQ259228 | A/mallard/Bavaria/185-8/2008(H1N1) | 100% | 94% |
| 58 | EPIISL3898 | EPI21248 | A/duck/Guangxi/53/2002(H5N1) | 8 | | DQ997525 | A/goose/Guangdong/xb/2001(H5N1) | 100% | 99% |

**Table S6.** Examples of rules generated by OneR, JRip and PART for (A) two-class and (B) three-class H1N1 dataset containing concatenated alignments of IAV proteins.

(A) Two-class virulence grouping

| Method | Rule(s) | Summary |
|---|---|---|
| OneR | NS2.107:<br>F -> Avirulent<br>L -> Virulent<br>(45/68 instances correct) | === Summary ===<br><br>Correctly Classified Instances 45 66.1765 %<br>Incorrectly Classified Instances 23 33.8235 %<br>Kappa statistic 0.3235<br>Mean absolute error 0.3382<br>Root mean squared error 0.5816<br>Relative absolute error 67.6471 %<br>Root relative squared error 116.316 %<br>Total Number of Instances 68<br><br>=== Confusion Matrix ===<br><br>a b <-- classified as<br>12 22 \| a = Avirulent<br>1 33 \| b = Virulent |
| JRip | JRIP rules:<br>===========<br><br>(NS2.107 = F) => Vir_two_classes=Avirulent (13.0/1.0)<br>=> Vir_two_classes=Virulent (55.0/22.0)<br><br>Number of Rules : 2 | === Summary ===<br><br>Correctly Classified Instances 45 66.1765 %<br>Incorrectly Classified Instances 23 33.8235 %<br>Kappa statistic 0.3235<br>Mean absolute error 0.4154<br>Root mean squared error 0.4557<br>Relative absolute error 83.0769 %<br>Root relative squared error 91.1465 %<br>Total Number of Instances 68<br><br>=== Confusion Matrix ===<br><br>a b <-- classified as<br>12 22 \| a = Avirulent<br>1 33 \| b = Virulent |
| PART | PART decision list<br>------------------<br><br>NS2.107 = L AND<br>NS2.3 = P: Virulent (14.0/2.0)<br><br>NS2.107 = F: Avirulent (13.0/1.0)<br><br>NS2.89 = I AND<br>NS2.60 = N: Virulent (6.0/2.0) | === Summary ===<br><br>Correctly Classified Instances 49 72.0588 %<br>Incorrectly Classified Instances 19 27.9412 %<br>Kappa statistic 0.4412<br>Mean absolute error 0.3426<br>Root mean squared error 0.4139<br>Relative absolute error 68.5183 %<br>Root relative squared error 82.7758 % |

| | | |
|---|---|---|
| | NS2.57 = Y: Virulent (31.0/14.0)<br><br>: Avirulent (4.0)<br><br>Number of Rules : 5 | Total Number of Instances 68<br><br>=== Confusion Matrix ===<br><br>a b <-- classified as<br>16 18 \| a = Avirulent<br>1 33 \| b = Virulent |

(B) Three-class virulence grouping

| Method | Rule(s) | Summary |
|---|---|---|
| OneR | PA.277:<br>F -> LOW<br>H -> INTERMEDIATE<br>S -> HIGH<br>Y -> LOW<br>(45/87 instances correct) | === Summary ===<br><br>Correctly Classified Instances 45 51.7241 %<br>Incorrectly Classified Instances 42 48.2759 %<br>Kappa statistic 0.2759<br>Mean absolute error 0.3218<br>Root mean squared error 0.5673<br>Relative absolute error 72.4138 %<br>Root relative squared error 120.3443 %<br>Total Number of Instances 87<br><br>=== Confusion Matrix ===<br><br>a b c <-- classified as<br>18 11 0 \| a = HIGH<br>9 17 3 \| b = INTERMEDIATE<br>6 13 10 \| c = LOW |
| JRip | JRIP rules:<br>===========<br><br>(NA.108 = T) => Vir_three_classes=LOW (18.0/6.0)<br>(PB2.82 = S) => Vir_three_classes=LOW (4.0/0.0)<br>=> Vir_three_classes=HIGH (65.0/38.0)<br><br>Number of Rules : 3 | === Summary ===<br><br>Correctly Classified Instances 43 49.4253 %<br>Incorrectly Classified Instances 44 50.5747 %<br>Kappa statistic 0.2414<br>Mean absolute error 0.3867<br>Root mean squared error 0.4397<br>Relative absolute error 86.9968 %<br>Root relative squared error 93.2721 %<br>Total Number of Instances 87<br><br>=== Confusion Matrix ===<br><br>a b c <-- classified as<br>27 0 2 \| a = HIGH<br>25 0 4 \| b = INTERMEDIATE<br>13 0 16 \| c = LOW |
| PART | PART decision list<br>------------------ | === Summary === |

```
PB2.108 = A: HIGH (5.0)

HA.453 = S AND
PB1.105 = N AND
NA.330 = S AND
M2.24 = D AND
NA.273 = G AND
HA.75 = E AND
PB2.215 = T AND
PB2.443 = K AND
NP.133 = I: INTERMEDIATE (4.0)

HA.453 = S AND
PB1.105 = N AND
NA.330 = S AND
M2.24 = D AND
PB1.391 = K AND
HA.113 = S AND
HA.383 = S AND
PA.529 = D AND
PB2.158 = E AND
NS1.101 = D AND
PB2.251 = R AND
HA.190 = D AND
HA.103 = A: LOW (10.0)

HA.453 = S AND
PB1.105 = N AND
PA.688 = E AND
M2.24 = D AND
PB1.391 = K AND
HA.113 = S AND
NS2.22 = E: HIGH (8.0/1.0)

HA.453 = S AND
PB1.105 = N AND
PA.688 = E AND
M2.24 = D AND
NS2.3 = P: INTERMEDIATE (7.0)

HA.453 = S AND
PB2.701 = D AND
PB1.105 = N AND
M2.24 = D AND
HA.420 = I AND
HA.225 = G AND
HA.186 = P: HIGH (4.0)

HA.453 = S AND
PB2.701 = D AND
PB1.105 = N AND
M2.24 = D AND
PB2.344 = V AND
PB2.187 = K AND
NA.451 = I AND
NA.300 = H AND
```

```
Correctly Classified Instances          69               79.3103
%
Incorrectly Classified Instances        18               20.6897
%
Kappa statistic                          0.6897
Mean absolute error                      0.1769
Root mean squared error                  0.3356
Relative absolute error                 39.7962 %
Root relative squared error             71.1988 %
Total Number of Instances               87

=== Confusion Matrix ===

  a  b  c   <-- classified as
 29  0  0 |  a = HIGH
  5 20  4 |  b = INTERMEDIATE
  4  5 20 |  c = LOW
```

| | |
|---|---|
| | NA.32 = I AND<br>HA.225 = D AND<br>HA.186 = S AND<br>PA.529 = D AND<br>PB2.82 = N AND<br>PB1.175 = N AND<br>PB1.353 = K: LOW (8.0/2.0)<br><br>HA.453 = S AND<br>PB2.701 = D AND<br>PB1.105 = N AND<br>M2.24 = D AND<br>HA.420 = I AND<br>NA.300 = H AND<br>NA.451 = I AND<br>PA.581 = M AND<br>HA.165 = S: INTERMEDIATE (15.0/2.0)<br><br>PB1.53 = G AND<br>PB2.82 = N AND<br>NA.300 = H AND<br>NA.376 = N: HIGH (15.0/2.0)<br><br>: LOW (11.0/1.0)<br><br>Number of Rules : 10 |

**Fig. S1. Accuracy distribution of 100 models learned independently from two-class and three-class BALB/C (A and B, respectively) and C57BL/6 (C and D, respectively) datasets using OneR, JRip and PART**. The datasets contain either the concatenated alignments or an individual alignment of IAV proteins. Wilcoxon signed-rank sum test is used to test the null hypothesis that the median of the accuracy is equal to the accuracy of zero rule learner (represented by the red dashed horizontal line). The level of significance of each test is flagged by the stars: * adjusted p-value <0.05, ** adjusted p-value <0.01 and *** adjusted p-value <0.001.

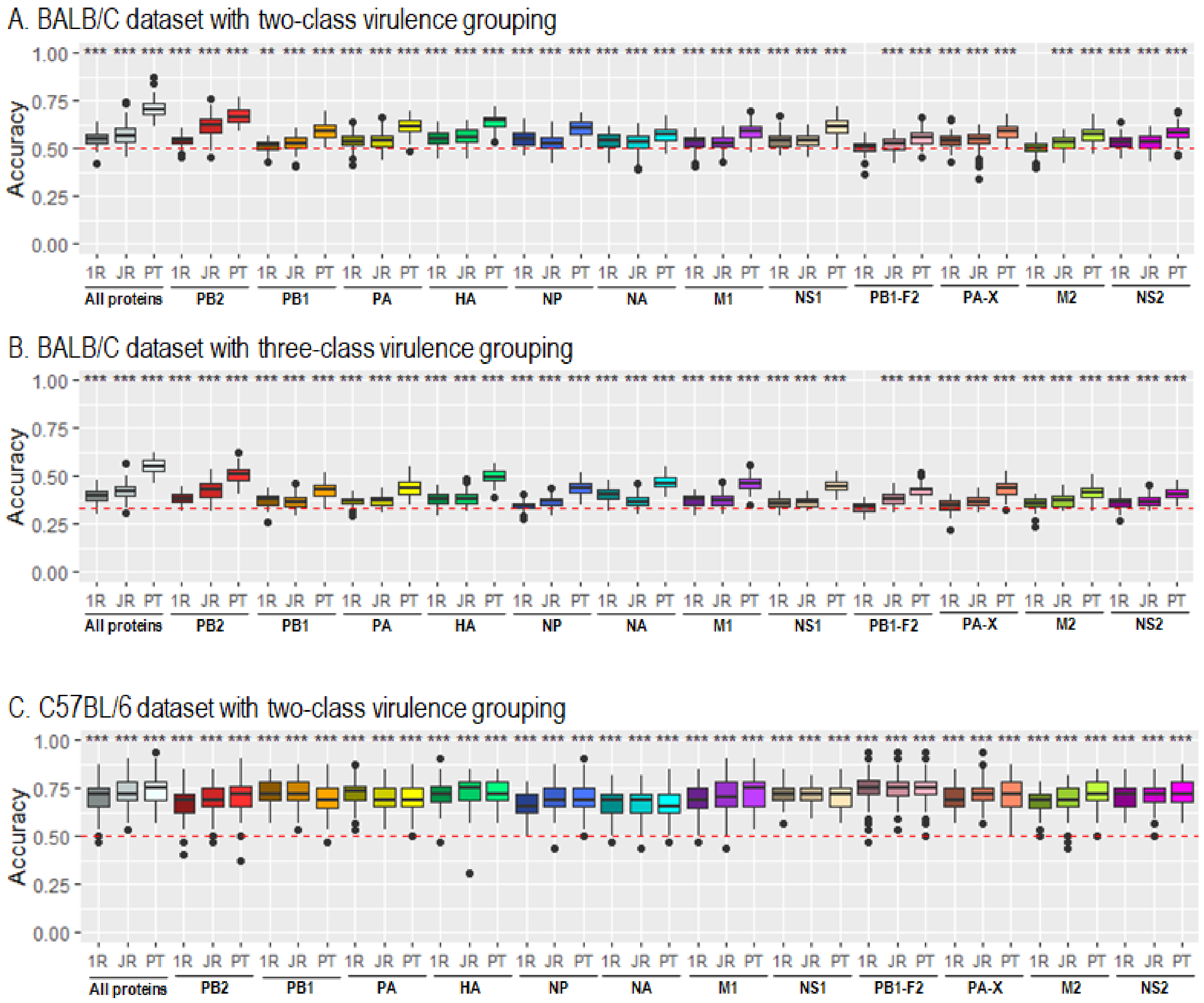

D. C57BL/6 dataset with three-class virulence grouping

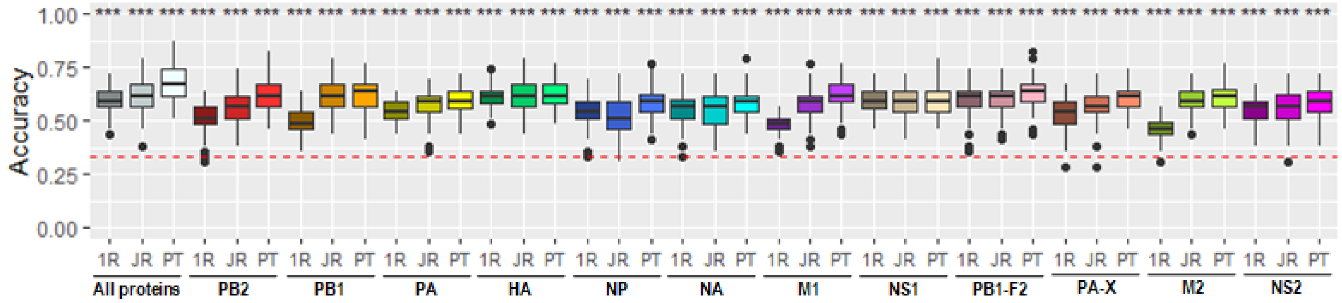

**Fig. S2. Accuracy distribution of 100 models learned independently from two-class and three-class H1N1 (A and B, respectively), H3N2 (C and D, respectively), and H5N1 (E and F, respectively) datasets using OneR, JRip and PART.** The datasets contain either the concatenated alignments or an individual alignment of IAV proteins. Wilcoxon signed-rank sum test is used to test the null hypothesis that the median of the accuracy is equal to the accuracy of zero rule learner (represented by the red dashed horizontal line). The level of significance of each test is flagged by the stars: * adjusted p-value <0.05, ** adjusted p-value <0.01 and *** adjusted p-value <0.001.

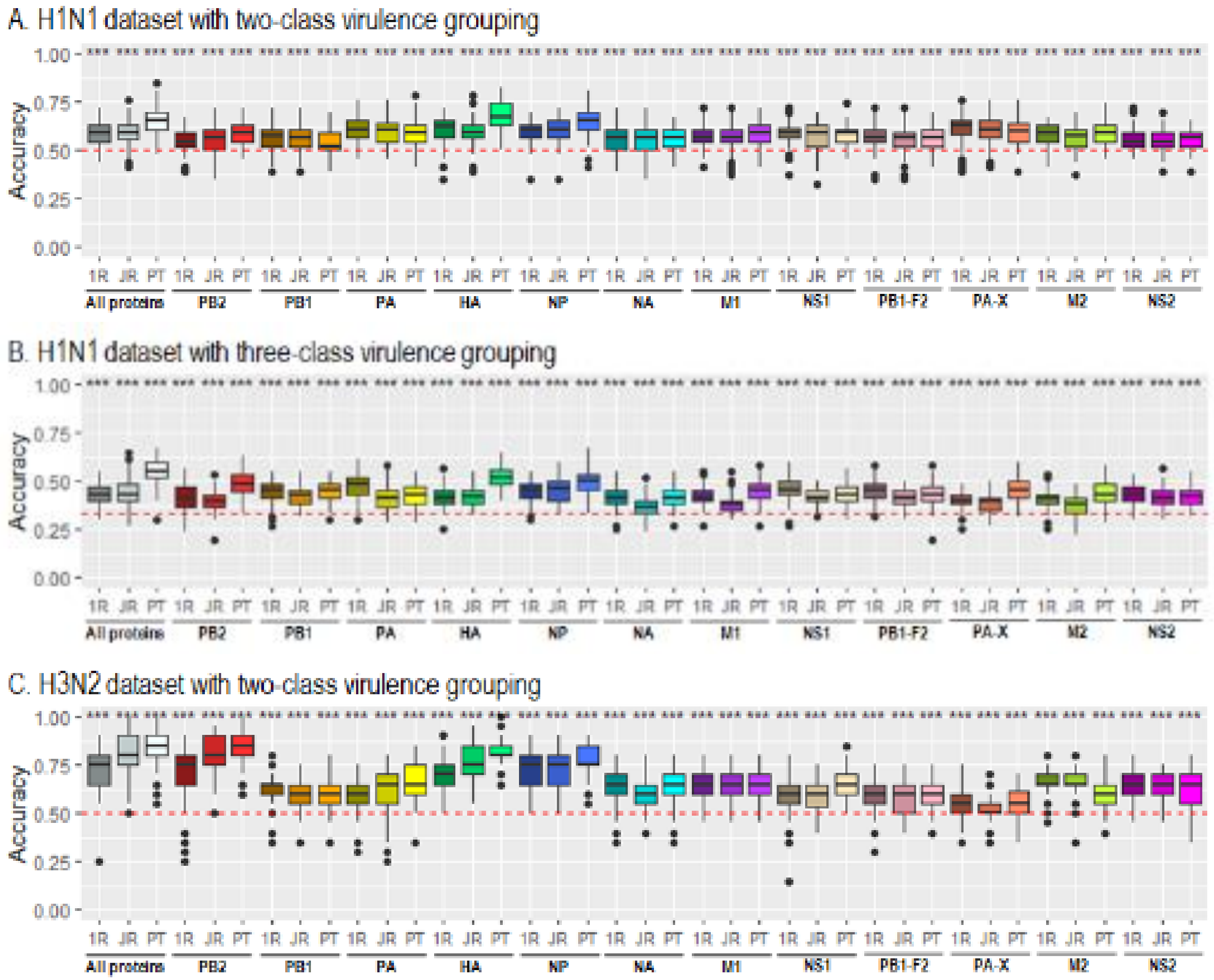

D. H3N2 dataset with three-class virulence grouping

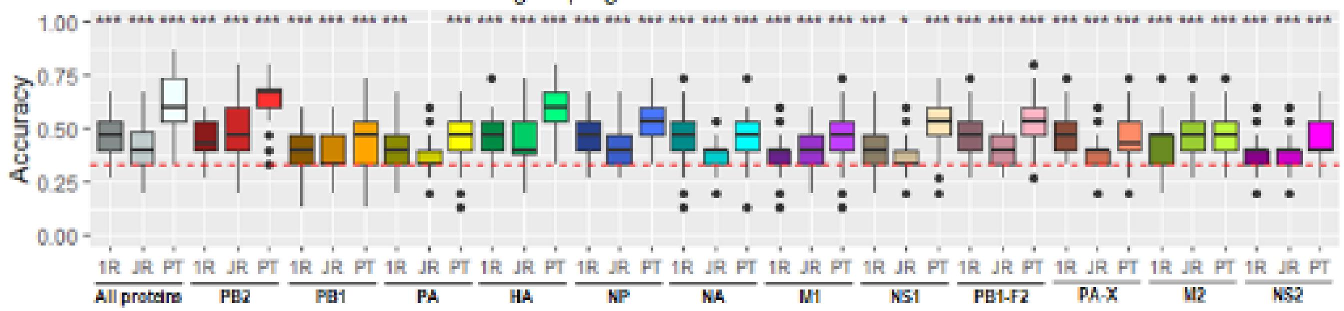

E. H5N1 dataset with two-class virulence grouping

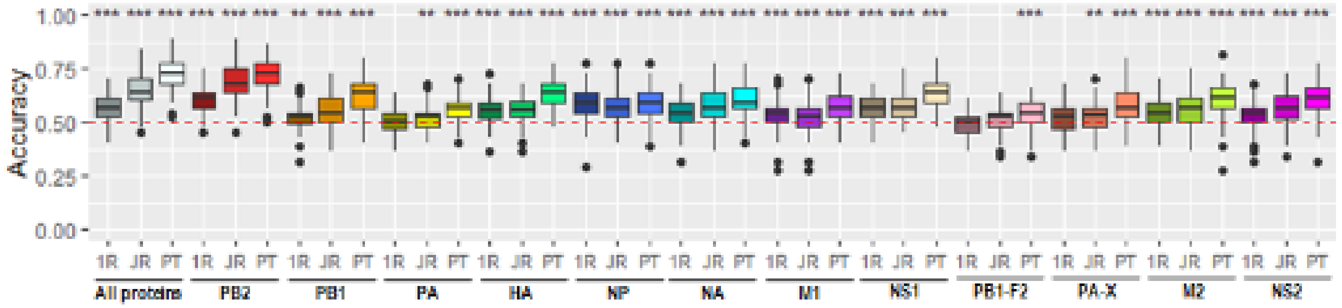

F. H5N1 dataset with three-class virulence grouping

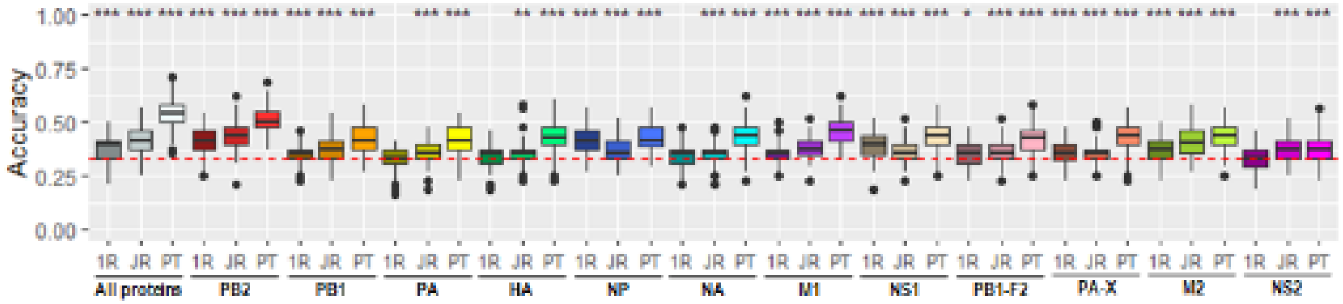

**Fig. S3. Accuracy distribution of 100 models learned independently from (A) two-class datasets and (B) three-class datasets using PART and random forest (RF).** The datasets contain the concatenated alignments of IAV proteins. Wilcoxon signed-rank sum test is used to test the null hypothesis that the median of the accuracy of PART models is greater than that of random forest (RF) models. The red dashed line indicates the accuracy of zero rule learner. The level of significance of each paired test is flagged by the stars: * adjusted p-value <0.05, ** adjusted p-value <0.01 and *** adjusted p-value <0.001.

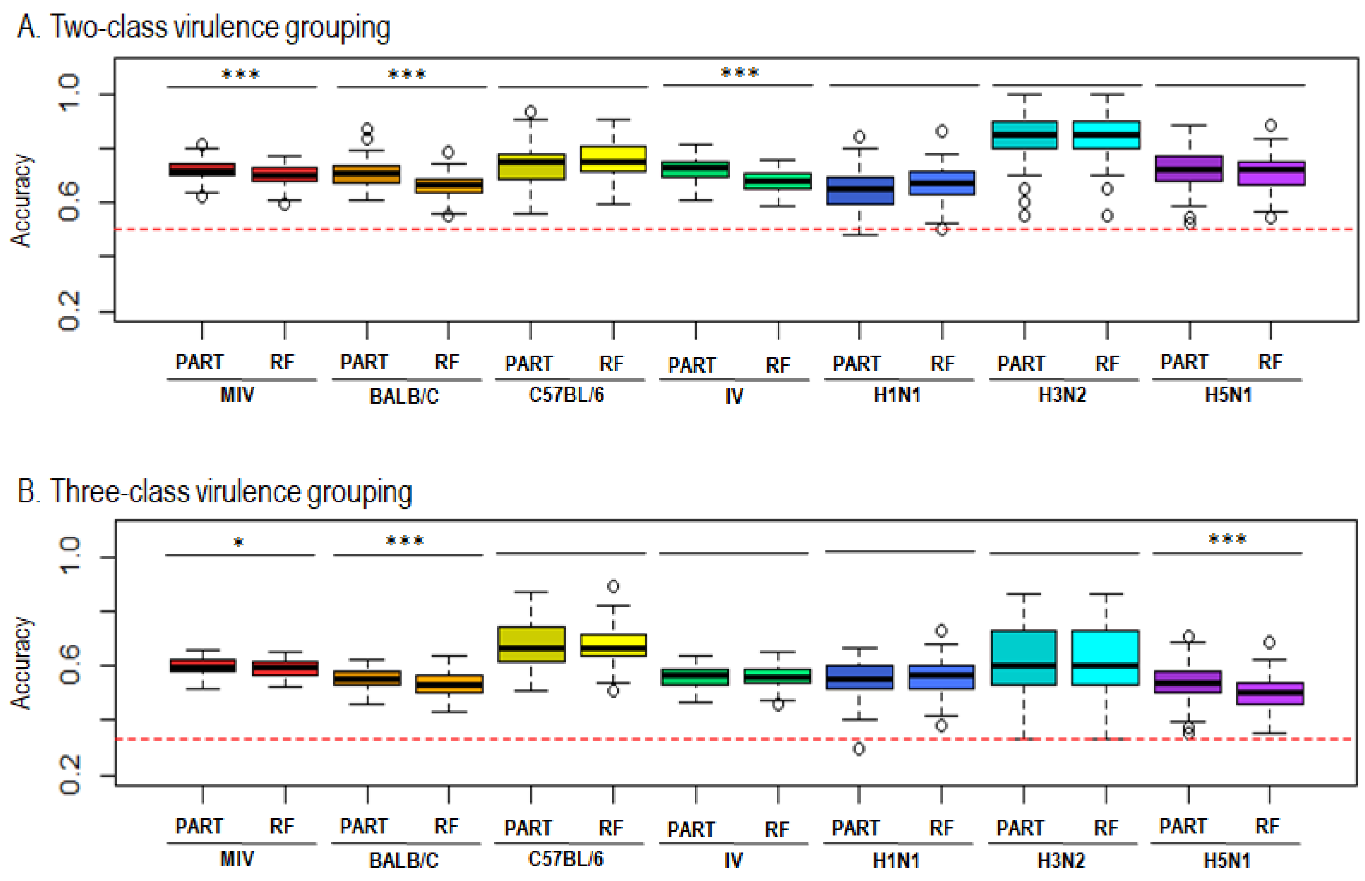
